# SPARK-ID: Dynamic DSB-sensor interactomes reveal modular nuclear repair networks coordinated by connector proteins

**DOI:** 10.64898/2026.07.27.740882

**Authors:** Alfredo Garcia-Venzor, Estefania De Allende-Becerra, Shai Kaluski-Kopach, Miguel Lurgi, Debra Toiber

## Abstract

DNA double-strand breaks (DSBs) activate repair pathways that must be coordinated with other cellular functions. Although DSB sensors SIRT6, Ku80, and MRE11 initiate repair, how they organize these nuclear processes remains unknown. SPARK-ID is a proximity-labeling strategy mapping DDR interactome dynamics. Using these sensors as baits, we resolved chromatin-associated interactomes from damage formation to recovery. The sensors shared an enriched repair core while capturing distinct interactors, allowing temporal specialization: SIRT6ID was biased toward RNA and chromatin regulation, Ku80ID toward telomere-associated and translational programs, and MRE11ID toward recombination and DNA synthesis. Modularity analysis showed these functions are organized into modules linked by “connectors”. Among them, Nucleolin linked DNA repair, RNA-metabolism, and nucleolar modules. Nucleolin depletion rewired DSB-sensor interactomes, altered repair-associated complex composition, expanded γH2AX domains, and reduced BRCA1, 53BP1, and phospho-ATM foci. Altogether, SPARK-ID reveals modular DSB-sensor interactomes whose robustness depends on connectors that integrate and constrain the DNA damage response.

**Graphical Abstract:** Three DSB sensors (MRE11, Ku80, SIRT6) orchestrate DNA repair through dynamic protein-protein interaction networks that expand upon DSB induction. Proximity labeling reveals distinct sensor interactomes that share a functional core of nuclear processes while incorporating sensor-specific interactors for pathway specialization. These networks contain connector nodes that coordinate multiple parallel nuclear functions, enabling functional diversification and conferring robustness to the DNA damage response. The sensors reshape their interactome structure and composition to integrate and constrain cellular responses through organized macromolecular complexes.

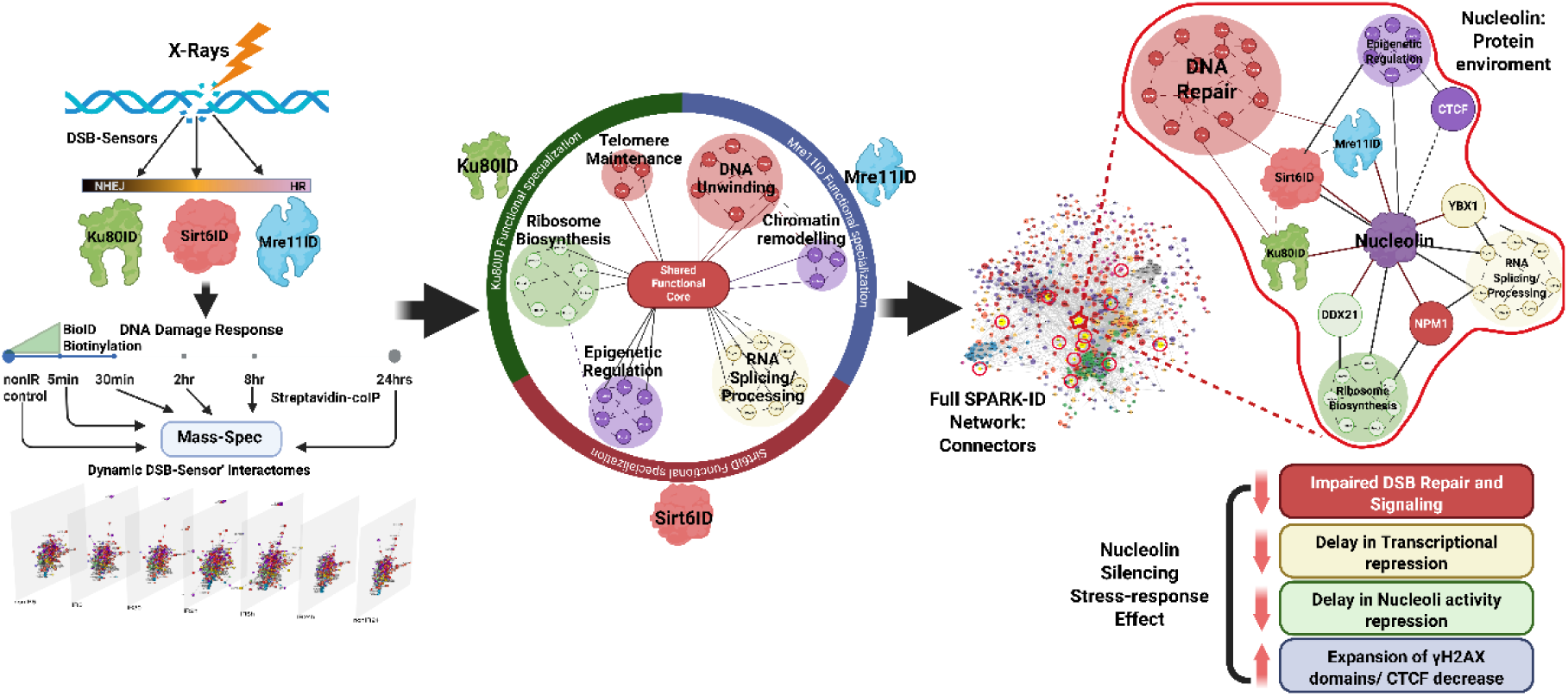

## INTRODUCTION

DNA double-strand breaks (DSBs) are among the most cytotoxic forms of DNA damage. They arise from endogenous sources, including replication and transcription stress^1^, as well as from exogenous insults such as ionizing radiation (IR) and chemotherapeutic agents. If not repaired correctly, DSBs promote mutations, deletions, chromosomal translocation, persistent checkpoint signaling, and large-scale genome instability^2^. Accordingly, defective DSB repair is linked to cancer, neurodegeneration, ageing-associated decline, and other age-related pathologies^3,4,4,5^.

Although DSBs are local DNA lesions, their repair requires the coordinated remodeling of multiple nuclear processes. DNA end recognition must be integrated with chromatin relaxation and restoration, transcriptional control, RNA processing, cell-cycle checkpoint signaling, protein turnover, and, eventually, the recovery of nuclear homeostasis ^6,7^. This coordination is dynamic: proteins are recruited, released, exchanged, and reorganized over time as the lesion transitions from detection to processing, pathway engagement, repair, and recovery. However, most experimental approaches capture either individual repair factors or static interaction snapshots, limiting our understanding of how DSB repair is organized as a temporally evolving nuclear network^8^.

DSB repair pathway choice is strongly influenced by the proteins that first recognize or organize the damaged DNA ends^9–11^. Non-homologous end joining (NHEJ) is initiated by the Ku70/80 complex and enables rapid ligation of DNA ends, whereas homologous recombination (HR) depends on the MRN complex, composed of MRE11, RAD50, and NBS1, to promote DNA end resection and template-directed repair ^9,12,13^. In addition to these canonical complexes, SIRT6 has emerged as a DSB sensor that binds DNA ends and contributes to both HR and NHEJ^11,14–16^. PARP1 also participates in DNA damage sensing and repair pathway coordination, particularly in the context of single-strand breaks and repair-associated poly-ADP-ribosylation^17–19^. Together, these sensors do more than recognize damage: they recruit signaling proteins, chromatin remodelers, repair enzymes, and effector complexes that shape the molecular environment surrounding the break. Despite extensive knowledge of individual repair pathways, how different DSB sensors organize their surrounding protein networks over time remains poorly understood. Direct comparisons between sensor-specific interactomes are limited, and many studies rely on loss-of-function systems that disrupt the sensor itself, making it difficult to distinguish the consequences of sensor absence from the normal organization of sensor-associated repair environments^20–22^. Moreover, transient and context-dependent protein interactions are difficult to capture using conventional affinity purification approaches^23,24^. A dynamic and comparative strategy is therefore needed to determine how DSB sensors coordinate shared and sensor-specific responses during repair progression. Here, we developed SPARK-ID (Sensor Proteome Alliance for Repair Kinetics) to map the dynamic protein-protein interaction networks of three DSB sensors: SIRT6, Ku80, and MRE11. Using proximity labeling coupled to chromatin enrichment and mass spectrometry, we followed sensor-associated interactomes from early damage recognition to late recovery after ionizing radiation^25,26^. This approach revealed that DSB sensors share a conserved repair-stress core composed of DNA repair, chromatin regulation, RNA metabolism, ribosome biogenesis, nuclear transport, and proteostasis-associated proteins. However, each sensor also recruited specific interactors that generated functional specialization: SIRT6 was preferentially associated with RNA and chromatin regulatory programs, Ku80 with telomere-associated and translation-related functions, and MRE11 with recombination and DNA synthesis-associated processes. Network modularity further showed that these functions are organized into structural modules connected by a limited set of connector proteins. Among these, Nucleolin emerged as a connector linking DNA repair, RNA metabolism, and nucleolar modules. Experimental Nucleolin depletion rewired DSB-sensor interactomes and impaired repair signaling, supporting a model in which DSB sensors coordinate modular nuclear repair networks through connector proteins that integrate and constrain the DNA damage response. To facilitate data exploration, we generated an online resource: SPARK, where the results presented here are displayed and accessible for public query (https://sparkid.bgu.ac.il/).

## RESULTS

### SPARK-ID maps dynamic DSB-sensor interactomes during repair progression

To determine how DSB-sensors organize their local protein environment during DSB repair, we developed SPARK-ID, a temporally controlled proximity-labeling strategy to reconstruct sensor-specific interactomes during the DNA damage response (DDR). SPARK-ID combines the inducible degron-based stabilization (via Shield1) ^27^ and the biotin-based labeling (BioID)^28^, confirming the induction and degradation with each of the three DSB-sensors: SIRT6 ^11^, Ku80, a component of the Ku-complex ^22^, and MRE11, the catalytic subunit of the MRN complex ^29^(**Figure 1A-B**), allowing the capture of early, intermediate, and late DSB-sensor interactors, while minimizing background labeling. Transduction into SH-SY5Y cells demonstrated that all SPARK-ID constructs retained proper nuclear/chromatin localization, interactions with known partners, a fast induction upon Shield1 addition, and high biotinylation activities (**Supplementary Figure 1A-K**).

**Figure 1.**
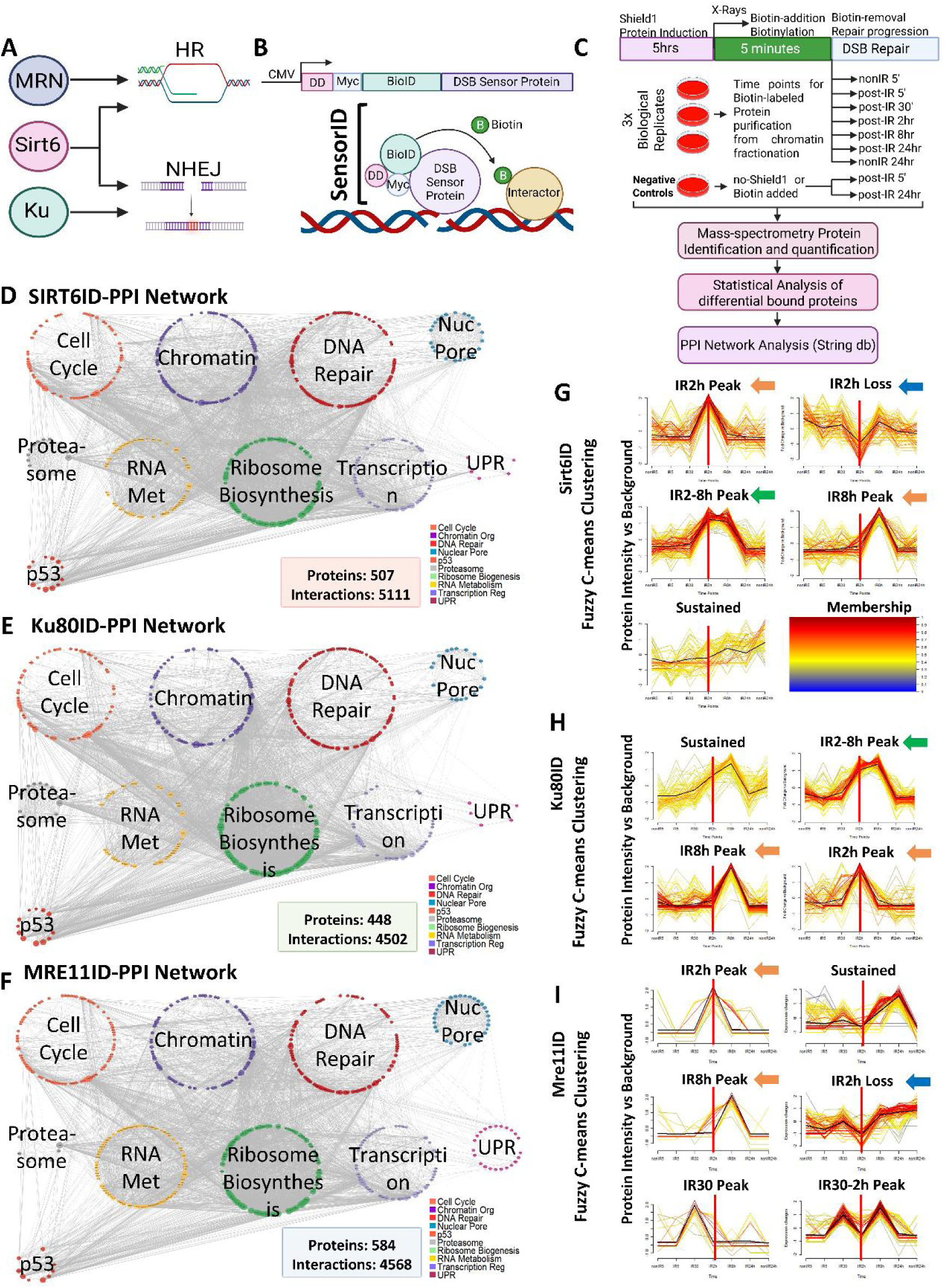
SPARK-ID captures the dynamic reorganization of DSB-repair networks. A) The three DSB sensors: MRN (MRE11–RAD50–NBS1; HR), Ku complex (Ku80/70; NHEJ), and SIRT6 (HR/NHEJ regulation). B) SPARK-ID chimera design, fusing each sensor to a biotin ligase (BioID) and destabilization domain (DD). C) Experimental timeline: interactomes mapped at basal (nonIR5′, nonIR24h), early (IR5, IR30), intermediate (IR2h), and late (IR8h, IR24h) stages post-irradiation (4 Gy). D–F) Global PPI networks for SIRT6, Ku80, and MRE11; nodes clustered and colored by nuclear core function. G–I) Fuzzy C-means clustering of interaction dynamics, with color-coded membership values.

Cells were exposed to 4Gy X-ray irradiation, and biotin was added immediately after damage induction. After a short labeling pulse, biotin and Shield were removed to reduce continued labeling and limit background signal. Chromatin-associated proteins were then purified by streptavidin co-immunoprecipitation at multiple time points comprising basal (non-irradiated), early (5- and 30-mins post-IR), intermediate (2h post-IR), and late restoration (8 and 24h post-IR) stages of the DDR. This design enabled temporal mapping of DSB-sensor-associated chromatin interactomes enriched for repair-proximal protein environments (**Figure 1C**). Because biotin labeling was confined to the initial 5-min pulse, later samples reflect persistence and redistribution of the initially labeled cohort, together with proteins co-purified through stable interactions, rather than direct de novo labeling at each collection time.

Using replicate mass spectrometry datasets and differential enrichment relative to negative controls, we identified dynamic interactomes composed of 507 SIRT6ID interactors, 448 Ku80ID interactors, and 584 MRE11ID interactors (**Figure 1D-F**). Interactor calls combined a large enrichment threshold (Log2FC>1.5) with empirical-Bayes moderated evidence (p_mod<0.05); FDR-adjusted values were retained in the complete results for transparency and sensitivity assessment. Manual annotation of these proteins revealed that all three networks contained major nuclear functional classes, including DNA repair and metabolism. These interactomes were highly connected and dynamic, displaying marked changes after irradiation; at IR5, all three interactomes showed an apparent contraction characterized by reduced recovery of peripheral interactors, followed by increased recovery and additional detectable associations at IR2h–IR8h. (**Supplementary Figures 1L-M**). This behavior was particularly evident at IR2h and IR8h, where the maximal deviation from the basal state was observed (**Supplementary Figure 1O**), whereas MRE11 showed a more prominent early response at IR30 and a second disconnection at IR2h, likely marking the transition towards active end-resection.

To identify temporal interaction patterns, we clustered protein abundance dynamics using Fuzzy C-means clustering (**Figure 1G-I**). The three sensors shared clusters with interaction peaks at IR2h and IR8h, suggesting that these stages represent common transition points during repair progression. SIRT6ID and Ku80ID also shared sustained IR2h-IR8h interaction patterns; SIRT6ID and MRE11ID shared an IR2h-disconnection pattern, whereas MRE11ID contained additional clusters with maximal recovery at IR30 or combined IR30-IR8h behavior, consistent with an earlier response linked to DNA end processing and recombination-associated functions.

Together, these results show that SPARK-ID captures dynamic DSB-sensor-associated chromatin interactomes and reveal that SIRT6, Ku80, and MRE11 undergo rapid and sensor-specific network remodeling after DNA damage. Although all three sensors engage broad nuclear repair-stress programs, their temporal interaction patterns suggest distinct modes of network expansion, specialization, and recovery during the DDR.

### DSB sensors mobilize a shared nuclear repair-stress core while acquiring sensor-specific functional programs

The broad functional composition of the SPARK-ID interactomes suggested that DSB sensors associate with multiple nuclear processes beyond canonical DNA repair. To define these programs more systematically, we performed time-resolved enrichment analysis using GO Biological Processes annotations. We identified a robust functional core, shared by all three sensors and sustained throughout the repair process, encompassing DNA repair, transcriptional regulation, chromatin organization, RNA metabolism, nucleocytoplasmic transport, and ribosome biogenesis (**Figure 2A**). These major functions constitute a conserved nuclear repair-stress core shared by the three sensors and highlight that DSB-sensor interactomes are not restricted to repair enzymes alone, but integrate repair with nuclear processes required to silence, remodel, process, and restore the damaged chromatin environment and achieve cell homeostasis.

**Figure 2.**
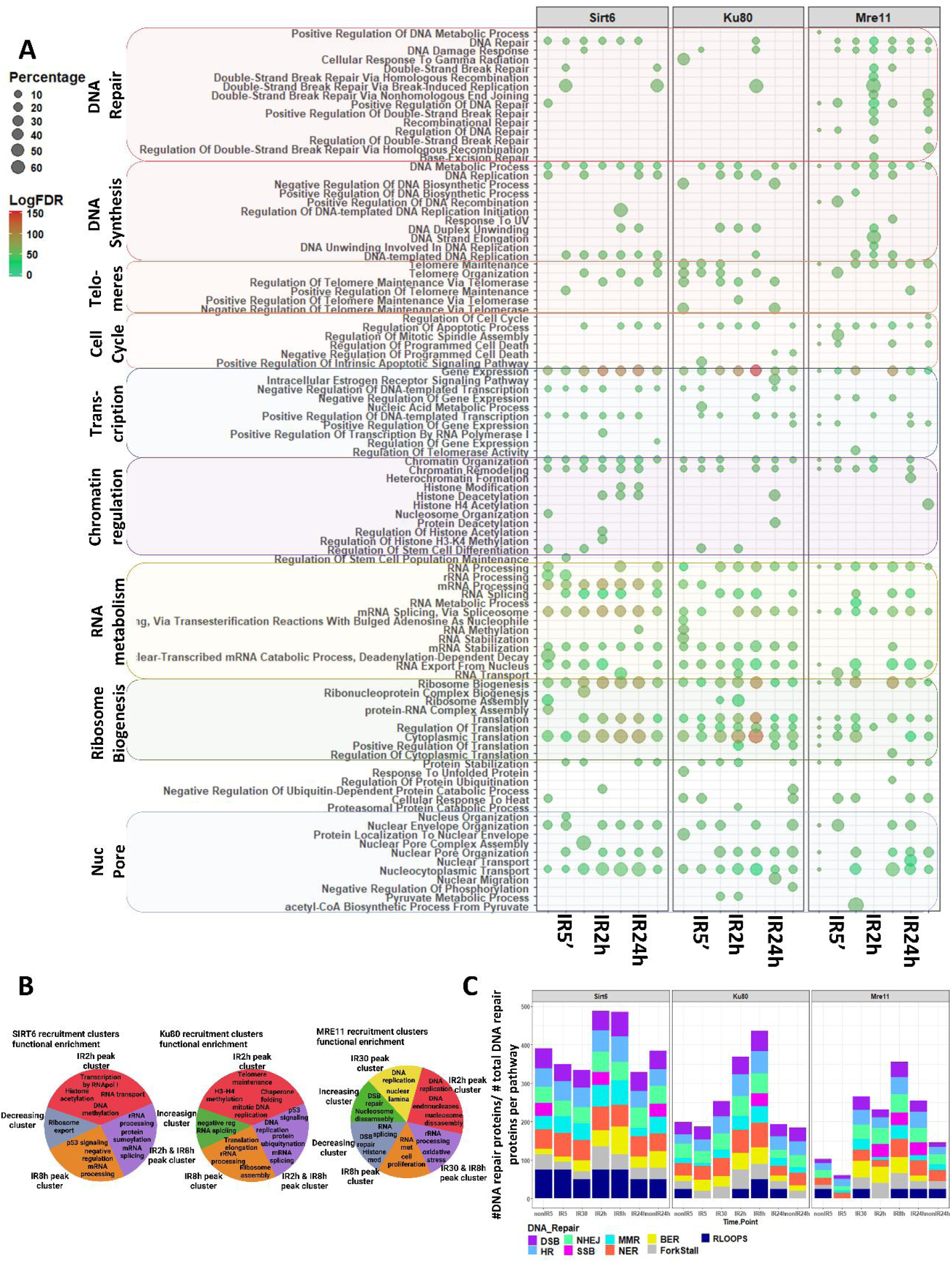
DSB sensors mobilize a shared nuclear repair-stress core and sensor-specific functional programs. A) Time-resolved GO Biological Process enrichment of the SIRT6ID, Ku80ID, and MRE11ID interactomes. The top 25 nonredundant terms per time point, selected by Benjamini–Hochberg significance and biological representativeness, are collapsed into major functional categories (DNA repair/synthesis, telomere/cell-cycle regulation, transcription/chromatin regulation, RNA metabolism, ribosome biogenesis, nucleocytoplasmic transport); broader networks are in the supplement. B) Biological processes enriched within each sensor’s Fuzzy C-means clusters, summarizing sensor-specific temporal programs. C) Recruitment of DNA repair factors: bars show the fraction of interactors previously identified in genome-wide screens for each repair pathway, normalized to known factors per pathway.

Despite this shared core, each sensor displayed a distinct functional bias that defines its regulatory specialization (**Figure 2A, supplementary Figure 2**). The SIRT6 interactome displayed a high specialization in transcriptional repression, histone modification, rRNA transcription, and mRNA splicing, consistent with the recruitment of Sin3A, NuRD, MLL1, HDAC1, and the spliceosome complexes. Ku80 showed a specialization in telomere maintenance, ribosome and translational regulation, as well as nucleocytoplasmic transport, partially explained by the specific recruitment of translation initiation factors, several ribosome components, protein chaperones, and the DNA-PK complex. As expected, MRE11 prioritized homologous recombination and DNA replication (unwinding and synthesis), mainly through interactions with the MDC1-MRN complex, Toposome, PCNA-RFC, and several chromatin remodeling complexes (SWI/SNF and BRG1). This functional specialization locates each sensor in direct connection with specific functions that require tight modulation during the repair process: MRE11 regulates DNA synthesis and the replication fork, SIRT6 links DNA repair with transcription and RNA processing, while Ku80 generates a link with ribosomes and translation.

Functional analysis of the interaction clusters revealed that this specialization arises mainly at early and intermediate stages, whereas restoration stages (IR8h) coalesce into similar homeostatic functions. This early divergence is characterized by the specialization of SIRT6 in transcriptional, splicing, and chromatin regulation, Ku80 in telomere maintenance and DNA synthesis, and MRE11 in DNA replication, unwinding, and homologous recombination; all three evolve towards homeostasis-maintenance mechanisms of protein turnover, ribosome biogenesis, p53 signaling, and RNA processing during restoration (**Figure 2B, Supplementary Figure 3A-C**). This supports a divergence-convergence model, where sensors initially drive divergent, specialized programs, potentially contributing to repair-pathway bias, but later they converge on a unified mechanism to restore cellular homeostasis.

Finally, mapping these interactors onto genome-wide DNA repair screens^30–36^ revealed that all sensors recruit factors from multiple repair pathways (NHEJ, HR), including NER, MMR, and BER (**Figure 2C**). This suggests that DSB sensors orchestrate a broad, multipathway response to cope with the complex metabolic stress of DNA damage.

Together, these results indicate that DSB sensors mobilize a conserved nuclear repair stress core while acquiring sensor-specific functional programs. This shared-but-specialized organization suggests that SIRT6, Ku80, and MRE11 coordinate repair not as isolated pathway components, but as dynamic organizers of broader nuclear environments required for repair progression and recovery.

### DSB-sensor interactomes converge on a functional core with pathway-specific diversification

To understand how the DSB-sensor interactome achieves both functional specialization and convergence, we compared the protein composition of SIRT6ID, Ku80ID, and MRE11ID interactomes. Across the complete time course, 249 proteins were shared by all three sensors, defining a conserved interactor core. In contrast, each sensor also contained unique interactors, with 82 proteins specific to SIRT6ID, 60 to Ku80ID, and 271 to MRE11ID (**Figure 3A**). Thus, DSB-sensor interactomes combine a shared molecular scaffold with large sensor-specific protein sets.

**Figure 3.**
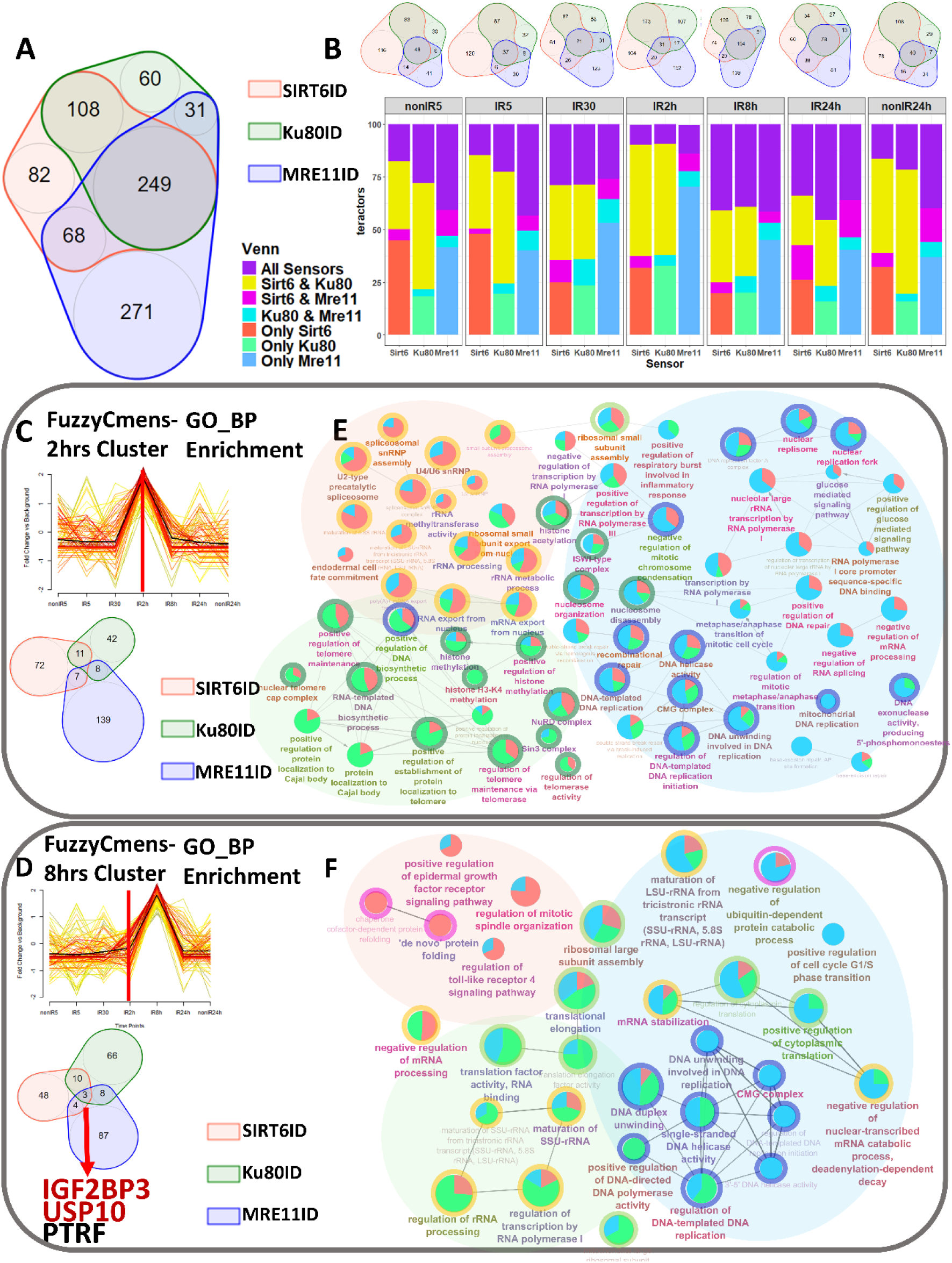
Sensor-specific interactors drive early functional divergence and late homeostatic convergence. A) Venn diagram of aggregated SIRT6ID, Ku80ID, and MRE11ID interactomes, showing the shared core and sensor-specific interactors. B) Temporal evolution of interactor overlap: stacked bars show shared and unique proportions per DDR stage, with time-specific Venn diagrams above (maximal divergence at IR2h, increased convergence at IR8h). C–D) Protein overlap between sensors within the C) IR2h and D) IR8h Fuzzy C-means clusters. E–F) ClueGO networks of processes enriched among E) IR2h and F) IR8h cluster proteins; nodes are enriched GO terms, pie charts show each sensor’s contribution.

The degree of overlap changed markedly during DDR progression. Time-resolved comparison showed that IR2h was characterized by high sensor specificity, whereas IR8h showed increased overlap between the three networks (**Figure 3B**). This temporal pattern mirrors the functional transitions described above: DSB sensors diverge during intermediate repair stages, when pathway-specific processing and chromatin remodeling are most active, and converge during late recovery, when homeostasis-associated functions become dominant.

We next asked whether the interaction clusters responsible for these transitions were composed of shared or sensor-specific proteins. The IR2h dynamic clusters showed minimal overlap between sensors, indicating that early functional divergence is largely driven by distinct protein sets (**Figure 3C**). These clusters were composed of proteins driving the specialization of SIRT6 in RNA processing, Ku80 in telomere maintenance and histone methylation, and MRE11 in DNA replication and nucleosome disassembly (**Figure 3E**). Therefore, the specialization observed at IR2h is not simply a change in abundance of the same proteins but rather reflects the recovery of sensor-specific interactors. A similar principle was observed at IR8h. Conversely, although the IR8h clusters are also composed mainly of specific interactors, their functions converged into RNA processing, ribosome biogenesis, and translation regulation (**Figure 3D-F**). This suggests that each sensor employs different molecular tools to achieve the same homeostatic goal. Only three proteins, IGF2BP3 (RNA binding), USP10 (deubiquitinase), and PTRF (Cavin-1), were shared across all sensors at this specific restoration stage, identifying them as candidate shared factors for homeostasis restoration. Double-peak clusters contained sets of proteins that sustain DNA replication (SIRT6/Ku80) and RNA processing (MRE11) throughout the entire DDR, suggesting that these clusters may contain sustained repair-associated programs (**Supplementary Figure 4**). Together, these results support a model in which DSB sensors use a common molecular core but recruit distinct interactors to execute specialized functions. This organization may provide robustness to the DDR by allowing similar nuclear processes to be maintained through alternative sensor-specific protein assemblies, making the repair process permissive to some degree or missing/mutated proteins.

### DSB-sensor interactomes are organized into structural modules linked by connector proteins

SPARK-ID experimentally defined the network nodes, whereas prey–prey edges were reconstructed from previously supported physical interactions in STRING; the resulting topology should therefore be interpreted within this physical-PPI scaffold. The functional diversification of DSB-sensor interactomes suggested that repair-associated nuclear processes may not be randomly distributed across the networks but instead organized into discrete structural compartments. To test this, we partitioned the networks into densely connected structural modules (**Figure 4A-B**)^37–39^. This analysis revealed that DSB-sensor interactomes are organized into multiple structural modules whose dominant functions converged into: DNA metabolism (red), containing repair, replication, and chromatin regulation; RNA processing (yellow), including splicing and processing factors; ribosome biogenesis (green) and nucleolar proteins; nuclear pore (blue) and nuclear lamina-associated proteins; and proteostasis-associated factors (grey) (**Figure 4B-C, Supplementary Figure 5A-D**). This structural organization was present across DSB-sensor interactomes but differed in size, composition, connectivity, and temporal dynamics.

**Figure 4.**
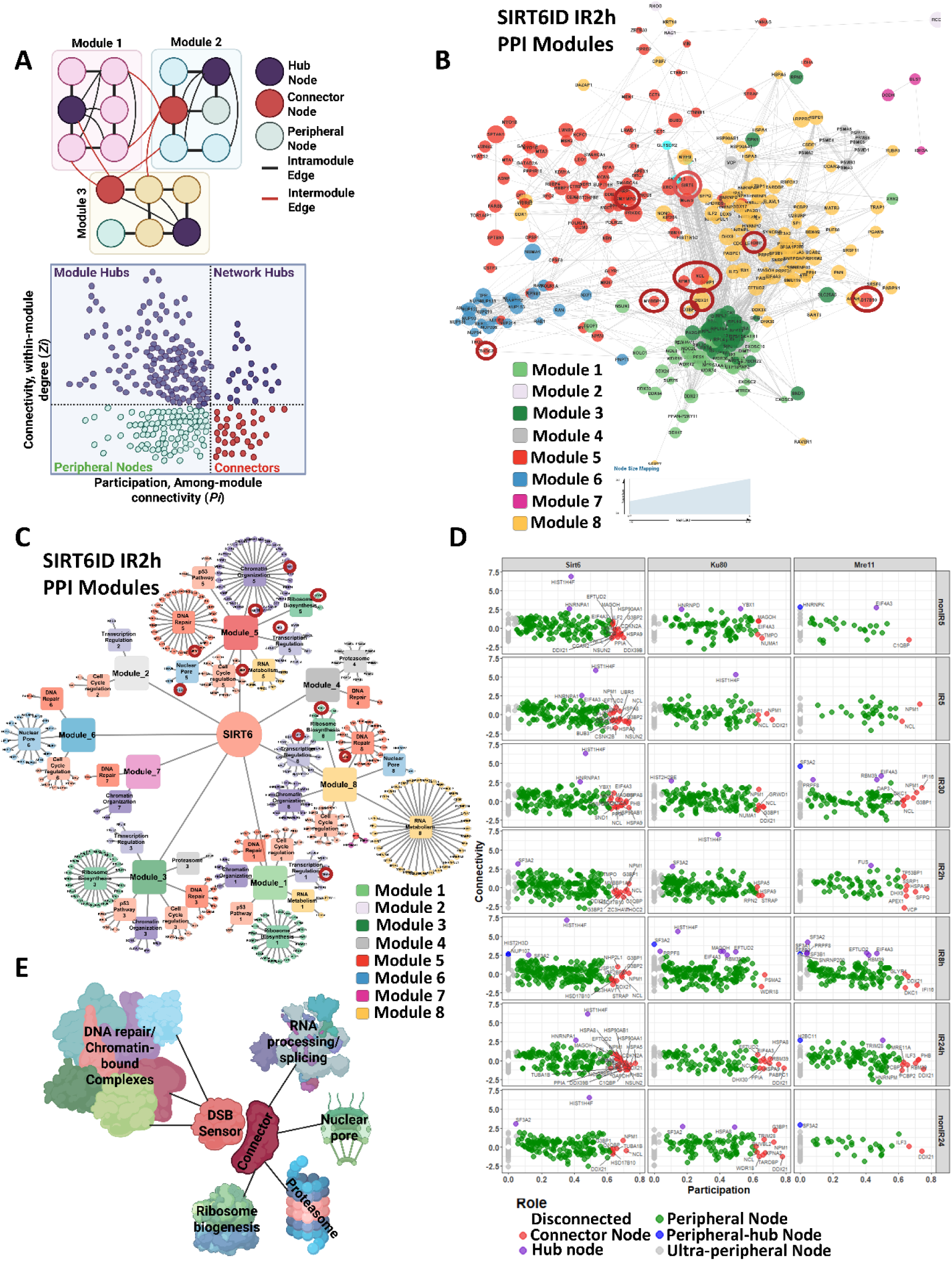
DSB-sensor interactomes are organized into structural modules linked by connector proteins. A) Modularity analysis schematic: PPI networks partitioned into modules by connection density, with nodes classified by within-module connectivity and participation coefficient (peripheral, ultra-peripheral, module hub, peripheral hub, connector). B) Modular architecture of the SIRT6ID IR2h network; nodes colored by module, connectors outlined in red. C) Functional map of the same modules, with nodes colored by biological function, showing physical segregation of DNA/chromatin, RNA metabolism, ribosome biogenesis, nuclear pore/envelope, and proteostasis modules. D) Topological classification across all three networks: participation coefficient versus within-module connectivity, with connectors (red) and module hubs (purple). E) Model: connectors (e.g., NCL, NPM1) bridge peripheral modules with the sensor core.

Importantly, these modules were not isolated; they were physically linked by a specific class of proteins: connector proteins. To classify the topological role of each interactor, we used two parameters: within-module connectivity (*Zi*), which identifies proteins highly connected inside their own module, and participation coefficient (*Pi*), which identifies proteins connected across different modules^38,40^ (**Figure 4A, D**). This approach classified proteins as peripheral and ultra-peripheral nodes, module-hubs, peripheral-hubs, or connector proteins. Module-hubs showed high intra-module connectivity, likely organizing local functional neighborhoods. In contrast, connector proteins showed high inter-module participation, occupied positions between module interfaces, likely acting as molecular hinges that link distinct biological processes (**Figure 4B-D, Supplementary Figure 5A-B**).

The abundance of connector and hub nodes changed during the DDR progression, indicating that network architecture is dynamically remodeled after DNA damage (**Supplementary Figures 5E, 6A-C**). Notably, connector expansion coincides with stages in which connector proteins mediate inter-module communication and provide flexibility to the DSB-sensor networks (**Figure 4E**). Therefore, SPARK-ID not only identifies repair-associated proteins but also reveals how these proteins are structurally organized into modular repair-stress networks.

Connector identity was also highly dynamic. During early and intermediate DDR stages, Nucleolin (NCL) and Nucleophosmin (NPM1) emerged as prominent connectors linking ribosome biogenesis and RNA metabolism modules with the DNA repair/chromatin core (**Figure 5A, Supplementary Figure 6A-C**). This suggests that nucleolar and RNA-associated factors are integrated into DSB-sensor networks during the repair response. As repair progressed toward recovery, helicases such as DDX21 and DDX39 acquired connector roles linking RNA metabolism with chromatin-associated processes, potentially contributing to transcriptional restart and chromatin restoration. In parallel, chaperones such as HSPA8 and HSPA9 connected repair-associated modules with proteostasis machinery, suggesting a role in the turnover or refolding of repair-stress proteins (Supplementary Figure 6A-C). Together, these results indicate that connector proteins are dynamic structural regulators of DSB-sensor interactomes.

**Figure 5.**
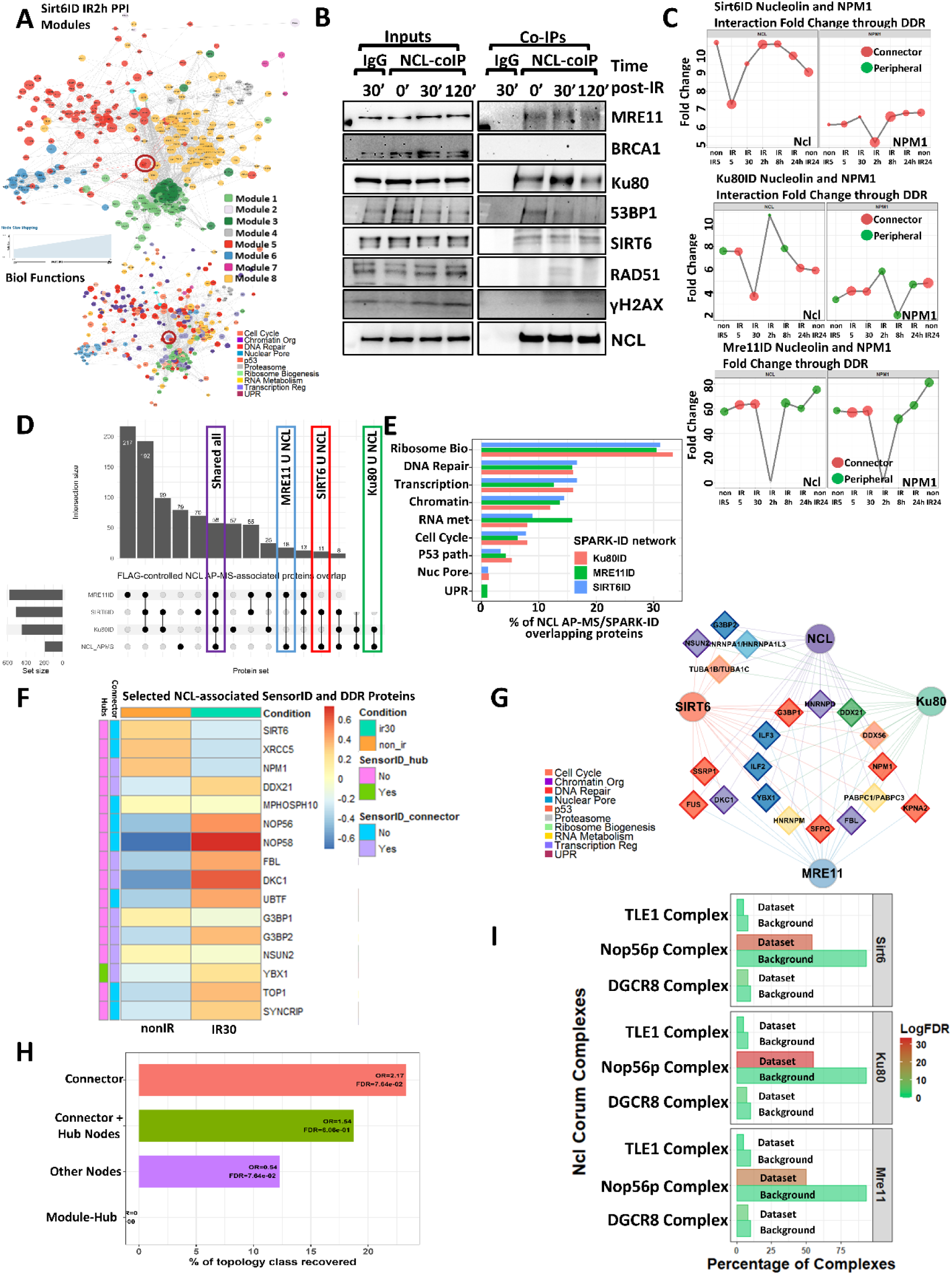
Nucleolin links DSB-sensor interactomes with RNA and nucleolar repair-stress modules. A) SIRT6ID IR2h network, colored by module (upper) and biological function (lower); NCL sits at the DNA-metabolism/ribosome-biogenesis/RNA-metabolism interface. B) NCL co-IP from irradiated SH-SY5Y cells, immunoblotted for MRE11, BRCA1, Ku80, 53BP1, SIRT6, RAD51, γH2AX, and NCL, with input and IgG controls; representative of four experiments. C) DDR-dependent abundance and topology of NCL and NPM1 across the three networks: y-axis, fold-change vs. negative control; point size, participation coefficient; color, topological role. D) UpSet analysis of overlap between FLAG-controlled NCL AP-MS proteins and each network (indistinguishable AP-MS groups counted as single features). E) Functional composition of the overlapping NCL AP-MS features per network. F) Heatmap of selected NCL AP-MS proteins shared with SPARK-ID/DDR pathways (WT nonIR and IR30); values are row-scaled, NCL-bait-normalized log2 intensities (four replicates), with annotations for SPARK-ID membership, connector/hub class, and FDR support. G) Network of NCL AP-MS features classified as SPARK-ID connectors or hubs; node color, dataset membership; shape, topological class; edge color, supporting dataset; NCL and the three sensors shown as bait nodes. H) Recovery of NCL AP-MS proteins per topological class; odds ratios and BH-adjusted FDRs from Fisher’s exact tests (unambiguous mappings, NCL bait excluded). I) CORUM NCL/NPM1 complexes represented in the networks (Nop56p, DGCR8, TLE1); bars compare recovered components with complex background, colored by enrichment significance.

### Nucleolin links DSB-sensor interactomes with RNA and nucleolar repair-stress modules

Among the connector proteins identified by SPARK-ID, NCL was particularly notable because it occupied the interface between DNA metabolism, RNA metabolism, and ribosome biogenesis modules (**Figure 5A**). Consistently, NCL co-immunoprecipitated with multiple DSB-repair proteins, including the three DSB-sensors SIRT6, Ku80, and MRE11, as well as 53BP1, RAD51, and γH2AX, in non-irradiated and irradiated SH-SY5Y cells (**Figure 5B**). Supporting the prediction that NCL is physically associated with DSB-repair protein environments.

SPARK-ID networks also revealed dynamic changes in NCL and NPM1 abundance and topological role during DDR progression, behaving as connector proteins during early stages in Ku80ID and MRE11ID networks, and a more sustained connector role in the SIRT6ID network (**Figure 5C**). These dynamics suggest that NCL and NPM1 form part of the flexible nucleolar/RNA-processing scaffold that is physically engaged by DSB sensors during repair progression.

To independently test whether NCL occupies a protein environment related to SPARK-ID networks, we analyzed NCL-associated proteins by IP-NCL affinity-purification followed by mass spectrometry (AP-MS). This analysis identified 58 SPARK-ID proteins that overlapped with the NCL-protein microenvironment, with a prominent overlap of the MRE11ID interactome (**Figure 5D, Supplementary Figure 7A**). Functionally, the shared NCL AP-MS/SPARK-ID proteins were mainly ribosome biogenesis, DNA repair, and transcription factors (**Figure 5E**). This functional composition mirrors the modules connected by NCL in the SPARK-ID networks.

The NCL-associated protein microenvironment was also responsive to DNA damage. Selected NCL-associated SPARK-ID and DDR proteins changed across non-irradiated and IR30 conditions, suggesting that the NCL-protein microenvironment is remodeled by genotoxic stress (**Figure 5F, Supplementary Figure 7B-C**). Notably, NCL-associated proteins preferentially associated with SPARK-ID connectors (**Figure 5G-H**). This indicates that the NCL AP-MS environment and the SPARK-ID network topology converge on a shared set of scaffolding and module-bridging proteins. Therefore, NCL is not merely present within the DSB-sensor networks; it is preferentially associated with the topological class of proteins predicted to mediate communication between repair, RNA metabolism, and nucleolar functions.

Finally, CORUM analysis identified NCL/NPM1-containing protein complexes represented within the SPARK-ID networks, including the Nop56p complex, involved in pre-rRNA maturation^41^; the DGCR8 complex, implicated in RNA processing^42^; and the TLE1 corepressor complex, associated with transcriptional repression^43^ (**Figure 5I**). These complexes mapped to the nucleolar and RNA-metabolism regions of the networks and showed irradiation-dependent remodeling across the three DSB-sensor interactomes (**Supplementary Figure 7D-F**). Together, these results support a model in which NCL and NPM1 participate in connector-rich nucleolar/RNA-processing assemblies that bridge DSB-sensor interactomes with broader nuclear repair-stress programs.

### Experimental removal of a connector node rewires DSB-sensor interactomes

To experimentally test the functional relevance of connector proteins identified by SPARK-ID, we depleted Nucleolin in SPARK-ID SH-SY5Y cells using an NCL-targeting shRNA (shNcl), with scrambled shRNA as control (shScr). We then applied the same SPARK-ID workflow and analytical strategy to reconstruct SIRT6ID, Ku80ID, and MRE11ID (ShScramble vs shNCL at IR2h for each sensor), a time point characterized by strong sensor-specific functional divergence and prominent NCL/NPM1 connector behavior in the original SPARK-ID networks (**Figure 6A**). Only replicates with confirmed NCL-depletion by immunoblotting and immunofluorescence were included in the analysis (**Supplementary Figure 8A-D**).

**Figure 6.**
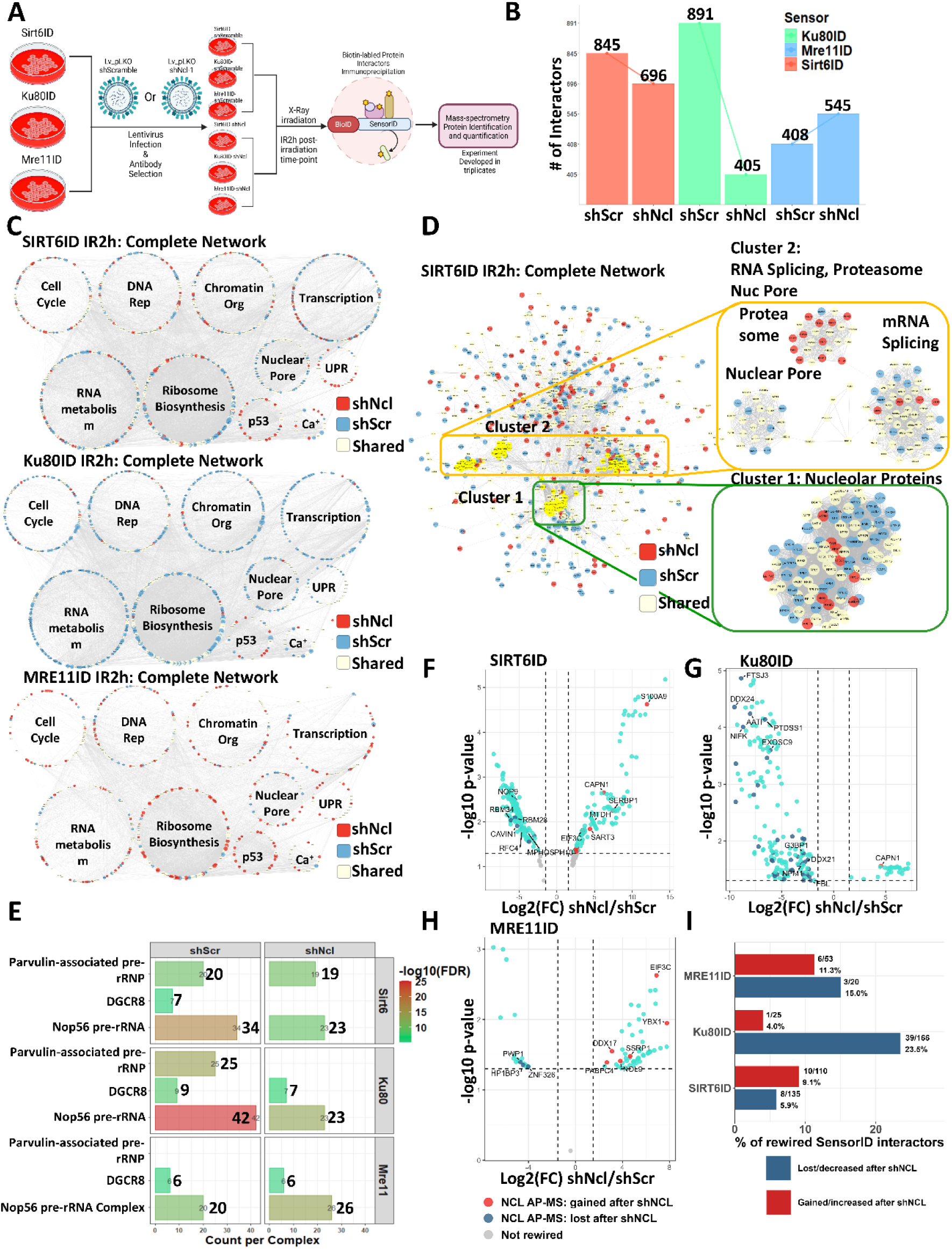
Experimental removal of NCL rewires DSB-sensor interactomes. A) Strategy: SPARK-ID SH-SY5Y cells transduced with scrambled or NCL-targeting shRNA, irradiated, analyzed at IR2h. B) Interactor counts in shScr and shNCL per network. C) Integrated shScr/shNCL networks, nodes grouped by nuclear function and colored as gained, lost, or shared after shNCL. D) MCODE subnetworks of SIRT6ID affected by NCL depletion (nucleolar, RNA-splicing, nuclear-pore, proteasome assemblies). E) CORUM enrichment of NCL-containing complexes across conditions (Parvulin-associated pre-rRNP, DGCR8, Nop56 pre-rRNA); bar length, detected components; color, significance. F–H) Direct shNCL-vs-shScr differential binding for F) SIRT6ID, G) Ku80ID, H) MRE11ID, restricted to nuclear SPARK-ID interactors; rewired at p < 0.05 and log2FC > 1.5 (positive = gained, negative = lost), with NCL AP-MS proteins highlighted. I) Recovery of NCL AP-MS proteins among rewired interactors; bars show the percentage of gained or lost interactors per network recovered by NCL AP-MS, labeled as NCL AP-MS proteins/total rewired (unambiguous mappings).

NCL-depletion markedly altered the DSB-sensor interactome composition. Ku80ID showed the strongest contraction, losing 486 proteins relative to shScr, while SIRT6ID lost 149 interactors. Surprisingly, the MRE11ID expanded after NCL depletion, gaining 137 interactors (**Figure 6B**). Thus, removal of a single connector protein did not produce a uniform collapse, but instead induced sensor-specific remodeling: contraction of the SIRT6ID and Ku80ID interactomes, and expansion of MRE11ID. To visualize this rewiring, we classified proteins as lost after shNcl, gained after shNcl or shared between shNcl and shScr. This revealed broad remodeling of functional neighborhoods associated with DNA repair, chromatin organization, RNA metabolism, ribosome biogenesis, nuclear pore, proteostasis, and stress signaling (**Figure 6C**). These results suggest that NCL does not function only as a passive bridge between modules. Rather, the sensor-specific rewiring is consistent with NCL acting as a connector-gatekeeper that helps define which proteins and complexes are incorporated into each DSB-sensor microenvironment.

### NCL regulates the access of RNA, nuclear-pore, and proteostasis complexes to DSB-sensor networks

We next asked whether NCL-dependent rewiring affected specific protein complexes within the DSB-sensor networks. MCODE analysis identified densely connected subnetworks corresponding to nucleolar/ribosome-biogenesis, spliceosome/RNA-processing, nuclear-pore, and proteasome-associated proteins, matching the structural modules bridged by NCL **(Figure 6D, Supplementary Figure 8E-F)**. These subnetworks were remodeled in a sensor-specific manner: in SIRT6ID and Ku80ID, NCL depletion reduced splicing, nucleolar, and nuclear-pore-associated interactors while increasing proteasome subunits, whereas MRE11ID instead gained nucleolar and splicing-associated components.

CORUM enrichment analysis reproduced this sensor-specific pattern at the level of annotated complexes, with NCL-containing pre-rRNA-processing complexes most strongly affected **(Figure 6E, Supplementary Figure 9A)**: SIRT6ID lost DGCR8 and Nop56p components; Ku80ID showed marked disruption of pre-rRNA-processing complexes; and MRE11ID gained a subset of Nop56p subunits. Consistently, the rewired proteins were enriched for RNA splicing, ribosome biogenesis, proteostasis, chromatin regulation, and DNA repair (**Supplementary Figure 9B-C)**.

To test whether these changes tracked with NCL’s own protein environment, we intersected the rewired interactors with the FLAG-controlled NCL AP-MS set **(Figure 6F-I)**. The overlap was strongest and most directional in Ku80ID, where 39 of 166 interactors lost after silencing were NCL-associated, versus only 1 of 25 gained, indicating that NCL depletion removes a defined subset of its own microenvironment from the Ku80 network. SIRT6ID and MRE11ID showed smaller, more bidirectional overlaps **(Figure 6I)**.

Together, these results indicate that NCL removal does not uniformly collapse the networks but instead produces sensor-specific contraction, expansion, or redistribution, and that the affected proteins are preferentially those NCL normally bridges. These findings are consistent with a model in which NCL acts as a connector-gatekeeper influencing access of RNA/nucleolar, proteostasis, and repair-associated complexes to DSB-sensor interactomes.

### NCL-silencing expands and disorganizes γH2AX-marked chromatin domains and alters CTCF-associated nuclear organization

The broad rewiring of DSB-sensor interactomes suggested that NCL depletion might alter not only interactome composition but also the nuclear organization of the DDR. Consistent with this, NCL-silencing changed DSB-sensor interactions with proteins implicated in chromatin-domain and repair-domain architecture, including cohesin/condensin subunits (PDS5A, NCAPG, SMC4, and regulatory subunits), CTCF, FACT, and Shieldin/53BP1-associated components⁴⁴⁻⁴⁶ **(Figure 7A)**. Because several of these factors constrain γH2AX-domain boundaries and loop extrusion, their loss raised the possibility that NCL depletion would relax the spatial containment of γH2AX-marked chromatin.

**Figure 7.**
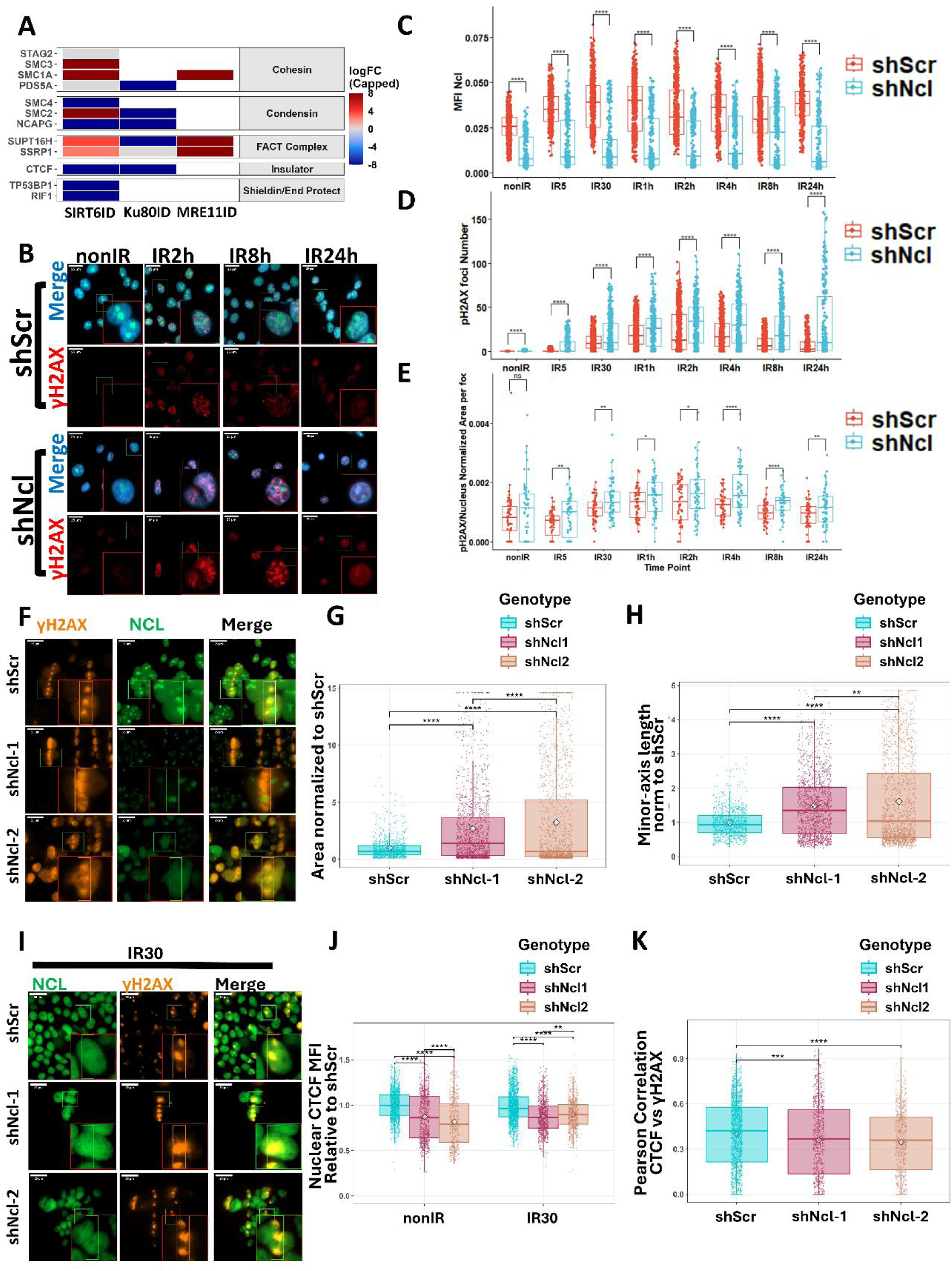
NCL depletion expands γH2AX-marked chromatin domains and alters CTCF-associated nuclear organization. A) Heatmap of NCL-dependent changes in sensor interactions with chromatin- and repair-domain factors (cohesin, condensin, Shieldin/53BP1-associated, FACT, CTCF); log2FC capped at ±8. B–E) γH2AX foci after NCL depletion in irradiated SH-SY5Y cells across DDR time points: B) representative γH2AX/NCL/DAPI images; C–E) nuclear NCL intensity, γH2AX foci number, and γH2AX foci area (multiple t-tests, Bonferroni; n = 4). F–H) Laser-induced γH2AX-domain geometry in HEK293 cells (shScr, shNCL-1, shNCL-2), fixed 30 min post-damage: F) representative images; G–H) domain area and minor-axis length perpendicular to the track (Kruskal-Wallis, Dunn’s; n = 4). I–K) CTCF and γH2AX after NCL depletion, processed as in F: I) representative images; J–K) nuclear CTCF intensity and CTCF/γH2AX Pearson colocalization (Kruskal-Wallis, Dunn’s; n = 4).

We therefore tested this directly. In irradiated SH-SY5Y cells, shNcl increased γH2AX-foci number and enlarged γH2AX-foci area, both in non-irradiated cells and at late DDR stages **(Figure 7B-E)**. Laser-induced DNA damage in HEK293 cells reproduced this phenotype in a geometrically constrained system: γH2AX domains were larger and wider after NCL-silencing, with more irregular borders and a higher frequency of hyper-expanded domains **(Figure 7F-H, Supplementary Figure 10)**. Thus, NCL is required to restrict the size and geometry of γH2AX-marked chromatin domains after DNA damage.

We next examined CTCF, a known determinant of γH2AX-domain boundaries. Under these conditions CTCF was not strongly enriched within γH2AX domains; however, NCL-silencing reduced nuclear CTCF levels and altered CTCF/γH2AX colocalization **(Figure 7I-K, Supplementary Figure 10)**, consistent with the loss of CTCF from the SIRT6ID and Ku80ID networks after shNcl **(Figure 7A)**. These changes indicate that NCL depletion perturbs CTCF-associated nuclear organization alongside γH2AX-domain expansion. Whether reduced CTCF contributes causally to that expansion, or reflects a parallel consequence of NCL loss, will require direct testing.

### NCL links DNA damage signaling with transcriptional repression and repair-pathway execution

NCL-silencing also changed DSB-sensor interactions with transcriptional regulatory complexes, including Mediator, NuRD, SWI/SNF, and RNA polymerase subunits **(Figure 8A)**. Because NCL bridges RNA/nucleolar modules with the DSB-sensor networks, this raised the possibility that NCL depletion would disrupt the transcriptional response to DNA damage. We tested this using BrU metabolic labeling. In control cells, irradiation induced nucleolar and nucleoplasmic transcriptional repression, most evident at IR30 and IR2h. NCL-silencing increased basal BrU incorporation and blunted this early repression of both nucleolar and nucleoplasmic RNA synthesis after irradiation **(Figure 8B-E, Supplementary Figure 11A-D)**. Thus, NCL is required for the normal coordination of DNA damage signaling with nuclear and nucleolar transcriptional homeostasis.

**Figure 8.**
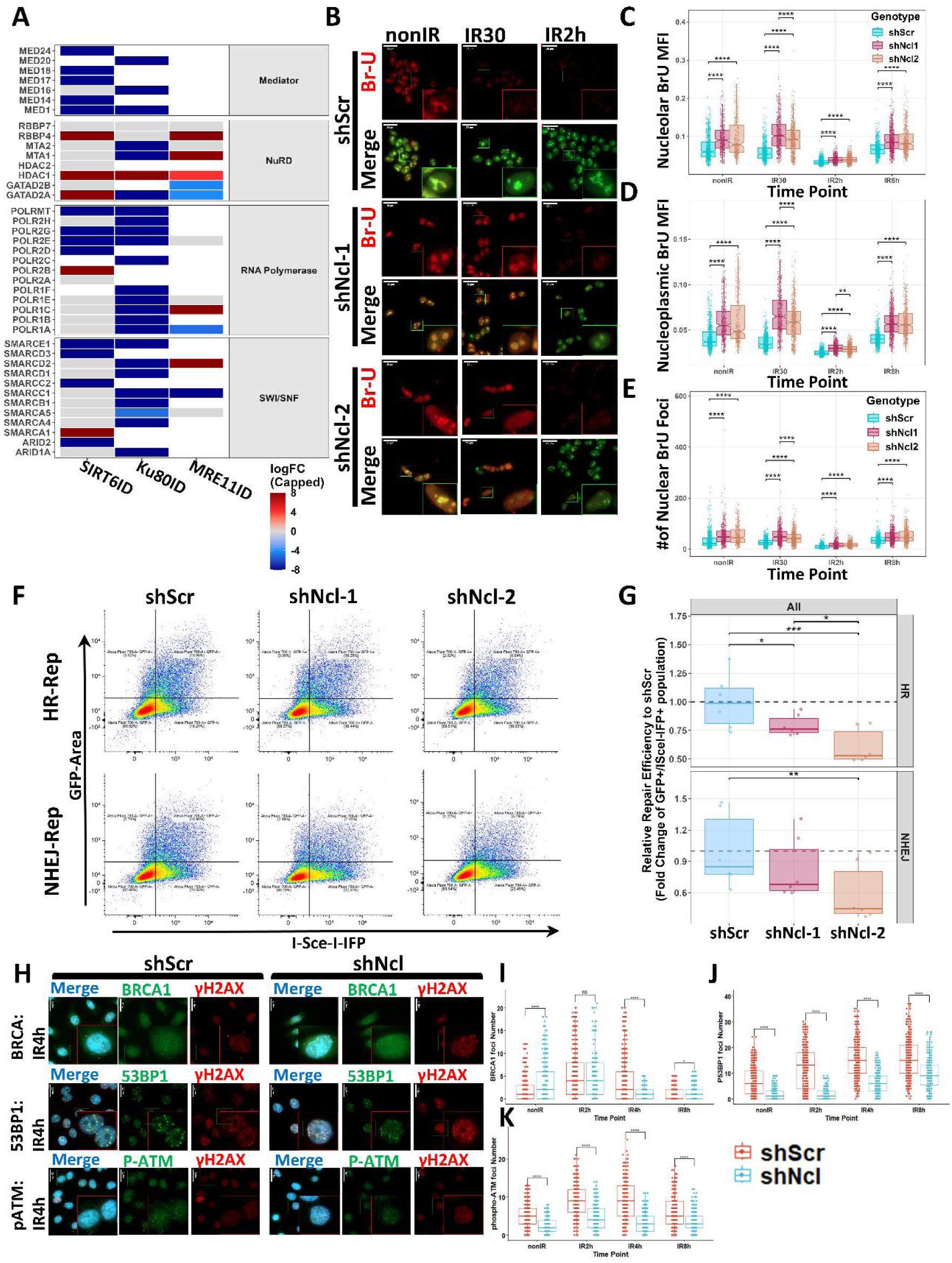
NCL depletion disrupts DNA damage-induced transcriptional repression and impairs HR/NHEJ repair output. A) Heatmap of NCL-dependent changes in sensor interactions with transcriptional regulators (chromatin remodelers, Mediator, RNA polymerase, transcription factors). B–E) BrU labeling of de novo RNA synthesis after NCL depletion in HEK293 cells (shScr, shNCL-1, shNCL-2) at nonIR, IR30, IR2h, IR8h: B) representative BrU/NCL/DAPI images; C–E) nucleolar BrU, nucleoplasmic BrU, and nuclear BrU foci (Kruskal-Wallis, Dunn’s, Bonferroni). F–G) HR and NHEJ reporter assays: F) representative FACS strategy; G) repair efficiency (GFP⁺/I-SceI-IFP⁺ among IFP⁺ cells, normalized to shScr; mixed-effects ANOVA, Tukey, FDR). H–K) Repair-factor foci in SIRT6ID SH-SY5Y cells (shScr/shNCL): H) representative IR4h images; I–K) BRCA1, 53BP1, and phospho-ATM foci number (multiple t-tests, Bonferroni).

We next asked whether NCL depletion affected repair-pathway output. HR and NHEJ reporter assays showed that NCL depletion impaired both pathways, with a stronger effect on HR **(Figure 8F-G)**. Consistently, NCL depletion reduced the number and size of BRCA1, 53BP1, and phospho-ATM foci after irradiation **(Figure 8H-K, Supplementary Figure 11E-J)**. Notably, this impaired recruitment of focal repair effectors occurred despite the expansion of γH2AX-marked domains described above; that is, larger damage-marked chromatin did not translate into more productive repair-factor assembly. NCL is therefore required not only to delimit γH2AX-domain expansion but also to support focal repair signaling and HR/NHEJ pathway execution.

Altogether, these loss-of-function experiments show that NCL depletion perturbs DSB-sensor interactome composition, γH2AX-domain organization, transcriptional homeostasis, and repair-pathway efficiency, consistent with a connector-gatekeeper function integrating these processes.

## DISCUSSION

Most interactome studies of DNA repair have focused on single factors or captured static snapshots. By mapping three mechanistically distinct DSB sensors across a common time course, SPARK-ID resolves how sensor-associated interactomes diverge during early and intermediate repair and reconverge toward late recovery, providing a temporally resolved landscape of the evolving DNA damage response that single-factor or single-timepoint approaches cannot.

The individual nuclear processes engaged during repair, chromatin remodeling, transcription, RNA metabolism, ribosome biogenesis, nuclear transport, and proteostasis, are individually well documented. What emerges from our data is their organization into discrete structural modules, and the identification of a limited set of connector proteins that physically bridge them. This modular-connector architecture offers a framework for understanding how the DNA damage response is coordinated systemically: how distinct sensors partition specialized functions while sharing a common core, and how the network is remodeled as the cell transitions from lesion-centered repair back toward homeostasis.

Within this framework, NCL emerged as a candidate connector-gatekeeper. NCL’s contribution to genome stability and DSB repair is established, including effects on ATM-dependent signaling and HR/NHEJ efficiency⁹³. Our results extend this role in three directions not previously linked to NCL: it occupies a topological position that bridges DNA repair, RNA metabolism, and nucleolar modules; NCL is required to restrain the size and geometry of γH2AX-marked chromatin domains; and it is needed for the normal damage-induced repression of nucleolar and nucleoplasmic transcription. Together, these places NCL as an organiser of the repair environment that acts well beyond its canonical nucleolar function.

### SPARK-ID reveals dynamic and modular DSB-sensor interactomes

Beyond identifying this modular organization, our data suggest a functional rationale for it. The recruitment and diversification of shared and sensor-specific modules followed an early disengagement from basal nuclear functions, an architecture that may increase network robustness by allowing each sensor to coordinate complementary functions while preserving a common response to DNA damage.

Systemic integration was concentrated through a small set of connector proteins, including NCL, NPM1, CSNK2B, and DDX21, that occupied interfaces between otherwise distinct macromolecular modules. Their dynamic topology supports a model in which connectors govern which protein environments associate with each sensor during repair progression, potentially linking the local lesion response to the broader nuclear state. Importantly, the changing representation of CORUM complex subunits across the DDR should not be read as evidence of complete complex assembly or disassembly; rather, it indicates that sensors engage selected components of known assemblies in a temporally regulated manner.

We deliberately mapped these interactomes in a context with largely intact repair machinery, capturing three DSB-sensor environments under conditions where repair pathways operate simultaneously and develop time-dependent crosstalk^44,45^. Loss-of-function strategies, commonly used to separate repair pathways mechanistically, would instead disengage the crosstalk nodes through which information passes between differently assembled protein environments^46^.

### SPARK-ID enables temporal mapping of DSB-sensor-associated protein interactomes

To our knowledge, the SIRT6ID, Ku80ID, and MRE11ID datasets represent one of the first systematic attempts to map temporal changes in three sensors simultaneously within a single system across the interactomes of three mechanistically distinct DSB sensors. By coupling BioID proximity labeling with an inducible degron, SPARK-ID extends previous static or transient interaction-capture approaches^21,47–50^. Most importantly, sampling from 5 min to 24 h after irradiation allowed us to track the progression from early damage recognition to repair-associated network diversification and, later, the restoration of nuclear homeostasis.

Our dataset significantly expands the known landscape of DSB repair, validating ∼43% of known SIRT6 interactions and providing the most comprehensive map of Ku80 and MRE11 to date ^21,22,47,49–51,51,52^. SPARK-ID used a 5-min post-IR labeling pulse followed by chase; later samples therefore track the chromatin-associated persistence, redistribution, and co-purified associations of the initially labeled cohort rather than direct de novo proximity at each collection time . Very rapid recruitment events occurring within seconds may therefore be underrepresented^53,54^. Faster proximity-labelling systems could, in principle, narrow this window, but each carries a liability incompatible with our design. APEX2-based labeling requires hydrogen peroxide, which itself induces oxidative DNA damage and is therefore unsuitable for a study of the DNA damage response.

In addition, the chromatin-enrichment strategy favors chromatin-bound protein environments and may reduce the recovery of soluble nucleoplasmic or loosely associated proteins. Mass-spectrometry biases also contributed to the loss of large, low-abundance, or poorly detected repair factors, including ATM, ATR, BRCA1/2, and RAD51, despite their detection by immunoblotting before MS. To reconstruct the physical organization of each experimentally defined protein set, SPARK-ID proteins were connected using experimentally supported physical protein-protein interactions from STRING^55^. Thus, the resulting networks represent temporally resolved, chromatin-associated DSB-sensor interactomes in which nodes were identified experimentally by SPARK-ID and edges correspond to supported PPIs among these proteins.

Despite these limitations, the temporal interactomes captured an early predominance of NHEJ-associated stages between IR5 and IR30^9,56,57^, followed by increasing engagement of HR-associated (IR2h)^58^, replication, chromatin-remodelling, and recovery programs at later stages^9,54^. This interval encompasses much of the established kinetics of the mammalian DSB response^59,60^. HR-associated functions were already detectable by IR30 in MRE11ID and SIRT6ID, whereas IR2h showed the strongest representation of DNA-synthesis and repair-related processes in these networks. Ku80ID remained comparatively enriched in chromatin-remodeling and telomere-maintenance functions. The early association of Ku80ID with telomere-maintenance machinery is notable because classical NHEJ is normally constrained at telomeres; however, replisome-associated alternative end-joining mechanisms at damaged telomeres could contribute to this pattern^61^.

MRE11ID also displayed marked temporal restructuring. At IR2h, MRE11 lost its association with C1QBP1, a reported inhibitor of MRE11 exonuclease activity^62^. This observation is consistent with, although does not directly demonstrate, a later stage of MRE11 activation or altered end-processing capacity. By IR8h, all three sensor networks showed increased representation of rRNA processing, RNA splicing, ribosome biogenesis, translation, and other homeostatic functions, suggesting a transition from lesion-centred repair toward broader restoration of nuclear organization. Notably, our transcriptional response experiments further confirmed the recovery of transcriptional activity 8 hrs after irradiation.

### DSB-sensor interactomes integrate repair with broader nuclear homeostasis

Our results reveal that DSB-sensors coordinate a multilayered response that extends beyond the damage site. Transcriptional regulation and RNA metabolism were particularly prominent in SIRTID^63,64^, consistent with the requirement to prevent conflicts between transcriptional machinery and DNA-repair complexes^65–67^. RNA-processing factors and RNA molecules can also contribute directly to both NHEJ and HR^68–70^. Accordingly, SIRT6ID showed extensive associations with RNA-binding proteins, including hnRNPA2B1, which participates in transcriptional regulation at DSBs^71^. Together, these findings place RNA metabolism within the physical architecture of the DSB response rather than a secondary consequence of damage signaling.

Nuclear-pore components and SUMO E3 ligases were also recruited to the sensor interactomes. Persistent DSBs can be relocated to nuclear pores through SUMO-dependent mechanisms that promote error-prone HR^72–74^, or alternative-NHEJ ^75^. In our networks, nuclear-pore and SUMO-associated interactions appeared as early as IR5. This does not demonstrate physical relocation of lesions at this time point but suggests that engagement of nuclear-pore-associated surveillance may begin earlier than previously appreciated.

Proteasome subunits were instead most prominent during later DDR stages. Proteasome-dependent degradation of polyubiquitinated repair factors contributes to repair progression and restoration of nuclear homeostasis ^76–81^. Our results extend this relationship by positioning proteasome-associated proteins within chromatin-bound DSB-sensor interactomes and identifying chaperone-like connectors that link proteostasis to repair modules. Whether these interactions represent local turnover at damaged chromatin or a broader nuclear stress response remains to be determined.

The extensive representation of nucleolar and ribosome-biogenesis proteins may be explained partly by direct rDNA damage, nucleolar-cap formation, and HR-mediated rDNA repair^82^, as well as by established nucleolar functions of SIRT6^83,84^, Ku80, and MRE11^85^. However, the nucleolus is also a stress-responsive nuclear compartment that participates more broadly in the DDR^86^, and multifunctional nucleolar proteins such as NCL and NPM1 can relocate to extranucleolar damage sites^85,87^. Their recruitment may therefore coordinate repair with the transient repression and subsequent restoration of ribosome biogenesis^88^. Consistent with this interpretation, our transcriptional measurements showed recovery of nucleolar and nucleoplasmic RNA synthesis by 8 h post-IR.

### Dynamic connector proteins coordinate modular repair-stress networks

Topological analysis provided a structural explanation for the integration of these diverse nuclear functions. The DSB-sensor interactomes were organized into interconnected macromolecular modules that broadly represented DNA and chromatin metabolism, RNA processing, ribosome biogenesis, nuclear transport, and proteostasis. Communication between these modules was concentrated through connector proteins, which formed physical routes between otherwise distinct functional neighborhoods^38^.

The identity and connectivity of these proteins changed during DDR progression. NCL and NPM1 linked ribosome-biogenesis and RNA-metabolism modules to the DNA-repair core during early and intermediate stages, whereas DDX21 assumed a stronger connector role during later recovery. Other connectors linked specific peripheral modules, including chaperone proteins associated with the proteasome and NSUN2 at the interphase with the nuclear-pore-associated proteins.

Modularity analysis has been widely used to identify organizing principles in complex biological networks^37^. Here, its application across multiple time points showed that modules can become more or less interconnected as DDR progresses. These changes should not be interpreted as direct assembly or disassembly of complete supramolecular complexes. Rather, they indicate temporally regulated changes in the physical relationships among selected components of known assemblies. This structural plasticity may allow DSB sensors to coordinate repair with transcription, RNA metabolism, nuclear transport, and recovery without requiring a single invariant repair complex.

### NCL shows properties consistent with a connector-gatekeeper role in DSB-sensor interactomes

We focused on NCL because it regulates ribosome biogenesis, associates with chromatin, and is rapidly recruited to DSBs^89–92^. Previous studies detected NCL recruitment within seconds after damage, whereas our temporal interactomes and co-immunoprecipitation experiments showed that its association with Ku80 and MRE11 remained detectable through IR30, whereas its association with SIRT6ID was more sustained. These findings extend previous observations by positioning NCL within RNA metabolism and nucleolar modules, which are physically connected to DSB-repair proteins.

NCL contributes to genome stability, and its depletion has been reported to impair both NHEJ and HR^93,94^. NCL has also been proposed to facilitate DNA-end resection through histone-chaperone activity and displacement of H2A-H2B dimers^92^. Its recruitment to damaged chromatin was reported to depend on PARP activity^53,54^. Our results suggest that these repair-associated functions occur within a broader NCL-linked protein environment that integrates repair with RNA processing, ribosome biogenesis, and chromatin organization.

FLAG-controlled NCL AP-MS provided an independent biochemical test of this model. Fifty-eight NCL AP-MS-associated proteins overlapped SPARK-ID networks and were dominated by ribosome biogenesis, DNA repair, transcriptional, chromatin, and RNA metabolism functions. This composition recapitulated the modules bridged by NCL in the network analysis. NCL-associated proteins also showed a trend toward preferential representation among SPARK-ID connectors, supporting convergence between the biochemical NCL-associated environment and the topological class predicted to mediate intermodule communication. The AP-MS environment was comparatively stable after irradiation, with selective changes rather than global remodeling, including a marked reduction in SIRT6 association at IR30.

NCL depletion provided direct evidence that this connector influences network composition. Its removal did not produce uniform disconnection: Ku80ID and SIRT6ID contracted, whereas MRE11ID expanded. Nucleolar, RNA-splicing, nuclear-pore, proteasome, chromatin, and repair-associated modules were remodeled in a sensor-dependent manner. The overlap with NCL AP-MS-associated proteins was particularly directional in Ku80ID, where 39 of 166 lost interactors belonged to the biochemical NCL-associated set, compared with only one of 25 gained interactors. This suggests that NCL stabilizes or facilitates access to a defined subset of the Ku80-associated environment. By contrast, the smaller bidirectional overlaps in SIRT6ID and MRE11ID are more consistent with remodeling than simple stabilization.

These findings support the role of NCL beyond being a passive scaffold. We propose that NCL may act as a connector-gatekeeper or module-boundary regulator, helping determine which proteins and assemblies gain access to each DSB-sensor interactome. This model is consistent with the previously reported physical association between SIRT6 and NCL^48^, and with our earlier observation that SIRT6 loss reduces nucleolar NCL levels^95,96^. The present results extend that relationship by showing that the consequences of NCL removal depend on the identity of the initiating DSB sensor, and different networks rely differently on NCL.

The cellular phenotypes further connect this network rewiring to repair organization. NCL-silencing expanded and distorted γH2AX-marked chromatin domains while reducing the assembly of BRCA1, 53BP1, and phospho-ATM (ser1981) foci and impairing both HR and NHEJ reporter repair. Thus, expansion of γH2AX-domains did not correspond to more productive repair-factor recruitment. Instead, NCL depletion appears to uncouple damage-associated chromatin from efficient repair-domain organization^97^.

NCL depletion also altered nuclear CTCF abundance and its spatial relationship with γH2AX. These changes are consistent with disruption of chromatin-boundary organization, since it has been shown that CTCF delimits γH2AX-dmains, and its silencing causes their abnormal expansion^98,99^. In parallel, NCL-silencing increased basal transcription and delayed the nucleolar and nucleoplasmic transcriptional repression normally induced by DNA damage^100^. Because suppression of transcription is required to limit conflicts between repair and transcriptional machinery^63,65,101^ this phenotype provides a functional link between the loss of RNA/transcriptional modules and defective repair.

Together, the network, biochemical, and functional results support a model in which NCL coordinates the organization of damage-associated chromatin with repair-factor recruitment and transcriptional control. Its loss disrupts the boundaries and composition of the repair environment, producing sensor-specific network contraction, expansion, or remodeling and ultimately impairing both HR and NHEJ.

### Limitations of the study

Several limitations should be considered. In addition to the limitations previously discussed, further limitations require consideration. SPARK-ID network construction also depends on previously documented physical PPIs. Although only experimentally supported STRING physical interactions meeting the selected confidence threshold were used, incompletely characterized proteins may have fewer documented edges and therefore appear less connected. Topological measurements should consequently be interpreted as properties of the currently supported physical network rather than a complete representation of every possible cellular interaction.

MCODE modules and CORUM overlap indicate coordinated representation of protein assemblies but do not establish that complete biochemical complexes were assembled or disassembled. Similarly, NCL AP-MS was performed under stringent high-salt conditions and compared with FLAG-empty controls. It therefore provides orthogonal evidence for NCL AP-MS-associated proteins rather than a definitive or exhaustive NCL interactome.

Finally, the experiments were performed in asynchronous SH-SY5Y populations, which are predominantly in G0/G1, and cell-cycle-specific interaction programs could not be resolved. Functional validation was also conducted across complementary SH-SY5Y and HEK293 systems, and some consequences of NCL depletion may reflect its broad effects on nucleolar function, transcription, and cellular homeostasis. Future studies using shorter labeling windows, cell-cycle-resolved analyses, and targeted perturbation of individual connector pathways will help distinguish direct organizational functions from secondary effects.

## Acknowledgements.

The study was funded by the European Research Council (ERC) under the European Union’s Horizon 2020 research and innovation program (grant agreement No 849029), the David and Inez Myers foundation, the Israeli Ministry of Science and Technology (MOST), The Israel Science foundation (No.: 422/23) the High-tech, Biotech and Negev fellowships of Kreitman School of Advanced Research of Ben-Gurion University. Thanks to Prof. Shai Pilosof and Prof. Barak Rotblat for their valuable insights. Thanks to Dr. Monica Einav for his previous work in plasmid construction and characterization. Thanks to Dr. Lior Onn for her valuable ideas and comments. Thanks to Dr Miguel Portillo for his ideas and comments during the development of this manuscript, and to Dr Uri Grupel for the development of the website. Thanks to the Technion Proteomics core facility for their invaluable help in sample processing, mass spectrometry analysis, and protein mapping and quantification.

## Author contributions

AGV: Conceptualization, hypothesis formulation, experimental work development, data analysis and manuscript writing. DT: Conceptualization, direction of the project, hypothesis formulation, experimental work development, data analysis, funding management, grant management, and manuscript writing. DABE: Data analysis, hypothesis formulation, and manuscript writing. SKK: management activities. ML: network concepts, application, and data analysis.

## Conflict of interest statements

The authors that participate in the execution and writing of this manuscript declare not to have any conflict of interest regarding its publication.

### Supplementary items titles and legends

**Supplementary Table 1. SPARK-ID Networks.** The first sheet contains node-level metrics, including fold change values across all analyzed time points, for the three DSB sensor proteins (SIRT6ID, Ku80ID, and MRE11ID). The second sheet contains the edge lists for the SIRT6ID, Ku80ID, MRE11ID, and the merged network SPARK-ID. Each interaction (edge) is represented with two edge weight metrics: Weight 1: The level of experimental evidence supporting the interaction. Weight 2: A composite score incorporating physical interaction strength and expression correlation between the protein pairs. The third sheet contains the LIMMA calculated Log2FC, p-values and FDR-values for all the statistically significant proteins.

**Supplementary Table 2. SPARK-ID Networks after shScr and shNcl Transduction.** The first sheet contains node-level metrics, including fold change values for all time points, for the three DSB sensor proteins under two conditions: control (shScr) and Nucleolin knockdown (shNcl), for the SIRT6ID, Ku80ID, and MRE11ID networks. The second sheet provides protein-protein interaction (PPI) edge lists for the SPARK-ID networks under the shScr and shNcl conditions. Each interaction includes two edge weight measurements: Weight 1: The amount of experimental evidence supporting the interaction. Weight 2: A composite metric reflecting physical interaction strength and expression correlation. The third sheet contains the LIMMA-calculated Log2FC, p-values, and FDR-values for all the statistically significant proteins.

## STAR Methods

### Experimental model

#### Cell culture

The SH-SY5Y (ATCC, Cat# CRL-2266, RRID: CVCL_0019) human neuroblastoma and HEK293 (ATCC, Cat# CRL-1573, RRID: CVCL_0045) cell lines were cultured in DMEM (Thermo, Cat# 41965, MA, USA) supplemented with 10% FBS, 2 mM L-glutamine, 100 U/mL penicillin, and 100 µg/mL streptomycin, under standard conditions (37 °C, 5% CO₂, humidified atmosphere). SPARK-ID protein induction was performed by adding Shield1 (Takara, Cat# 632188, Japan) at 50 nM for SIRT6ID and 500 nM for Ku80ID and MRE11ID. Biotinylation was performed by adding 50 µM biotin (Cayman Chemical, Cat# 22582, MI, USA) to the culture medium.

#### SPARK-ID SH-SY5Y cell lines generation

For the generation of SPARK-ID dynamic interactomes, we used three cell lines, SIRT6ID-SH-SY5Y, Ku80ID-SH-SY5Y, and MRE11ID-SH-SY5Y, derived from SH-SY5Y cells by lentiviral transduction of the SIRT6ID, Ku80ID, and MRE11ID chimeric transgenes (see Methods details for plasmid construction and cell-line generation). The SPARK-ID cell lines were cultured in DMEM (Thermo, Cat# 41965, MA, USA) supplemented with 10% FBS, 2 mM L-glutamine, 100 U/mL penicillin, and 100 µg/mL streptomycin, under standard conditions (37 °C, 5% CO₂, humidified atmosphere). One week before each streptavidin-directed immunoprecipitation, standard medium was replaced with biotin-depleted medium, and the cells were maintained under biotin-free conditions until exogenous biotin was added for labeling. Biotin-depleted medium was prepared by adding 2 mL of avidin solution (BioLock, IBA Life Sciences, Cat# 2-0205-050) to 500 mL of the supplemented DMEM described above, incubating for 25 min at 300 rpm, and re-sterilizing through a 0.2 µm filter.

For the SPARK-ID interactomes after Nucleolin silencing, the three cell lines were transduced with a second lentivirus encoding either an NCL-targeting shRNA (shNcl-2) or a non-targeting shRNA (shScramble) (see Methods details for cell-line generation). Transduced lines were maintained in the DMEM and biotin-depleted media described above, supplemented with 3 mg/mL Geneticin (G418, Formedium, England, Cat# G418S). For each streptavidin-directed co-immunoprecipitation, cells were grown in biotin-depleted medium for at least 1 week beforehand, and Nucleolin silencing was confirmed by Western blotting and immunofluorescence prior to mass spectrometry analysis.

### Methods details

#### Nucleoplasm-chromatin and chromatin fractions

For the HCl-based chromatin fractionation, cells were harvested at ∼80% confluence, detached by scraping, washed once with PBS, and centrifuged at 1,000 × g for 5 min at 4 °C. The pellet was resuspended in cytoplasmic lysis buffer (10 mM HEPES pH 7.4, 10 mM KCl, 0.05% NP-40, 0.5 mM PMSF, 10 nM TSA, phosphatase-inhibitor cocktail) at 50 µL per 10⁶ cells, incubated for 20 min on ice, and centrifuged at 19,000 × g for 10 min at 4 °C. The supernatant was collected as the cytoplasmic fraction. The pellet was washed three times with the same buffer (50 µL per 10⁶ cells), incubating 5 min on ice and centrifuging 5 min at 19,000 × g, 4 °C, between washes. The washed pellet was resuspended in 2 N HCl (30 µL per 10⁶ cells), incubated on ice for 30 min, and centrifuged at 19,000 × g for 10 min at 4 °C. The supernatant was collected as the chromatin-bound fraction and neutralized with 1 M Tris pH 8 (30 µL per 10⁶ cells).

The nucleoplasm–chromatin fractionation was performed as described¹⁰⁷. Briefly, cells were detached by scraping, washed once with PBS, and centrifuged at 1,000 × g for 5 min at 4 °C. The pellet was resuspended in sucrose buffer (5 µL per 10⁶ cells; 0.32 M sucrose, 3 mM CaCl₂, 2 mM MgOAc, 0.1 mM EDTA, 10 mM DTT, 0.5 mM PMSF, 10 nM TSA, phosphatase-inhibitor cocktail), and an equal volume of sucrose buffer containing 0.5% NP-40 was added. The homogenate was centrifuged for 10 min at 1,100 × g, 4 °C. The supernatant was collected as the cytoplasmic fraction, and the pellet was washed twice with sucrose buffer (5 µL per 10⁶ cells), centrifuging at 400 × g for 10 min at 4 °C after each wash. The pellet was resuspended in soluble-nuclear lysis buffer (5 µL per 10⁶ cells; 50 mM HEPES pH 7.4, 50 mM KCl, 300 mM NaCl, 0.1 mM EDTA, 10% glycerol, 1 mM DTT, 0.1 mM PMSF, 10 nM TSA, phosphatase-inhibitor cocktail) and incubated with rotation at 4 °C for 1 h. Samples were centrifuged at 10,000 × g for 3 min at 4 °C, and the supernatant was collected as the nucleoplasmic fraction. The remaining pellet was resuspended in chromatin lysis buffer (5 µL per 10⁶ cells; 50 mM HEPES pH 7.4, 3 mM MgCl₂, 300 mM NaCl, 1 mM DTT, 5 U/µL Benzonase, 0.5 mM PMSF, 10 nM TSA, phosphatase-inhibitor cocktail), incubated with rotation at 4 °C for 1 h, and centrifuged at 10,000 × g for 3 min at 4 °C. The supernatant was collected as the chromatin fraction.

#### Western Blotting

Protein samples were mixed with 4× Laemmli buffer to a final concentration of 1 µg/µL, loaded, and resolved on 12% SDS-PAGE gels. Proteins were transferred to nitrocellulose membranes (Pall Life Sciences, Cat# 66485, NY, USA), which were blocked with 5% milk in T-TBS for 1 h with orbital agitation. Membranes were incubated overnight with primary antibodies: anti-biotin (Abcam, Cat# ab53494, 1:1000), streptavidin-HRP (Thermo, Cat# N100, 1:3000), anti-myc-tag (Cell Signaling, Cat# 2276S, 1:1000), anti-SIRT6 (Cell Signaling, Cat# 12486, 1:1000), anti-Ku80 (Cell Signaling, Cat# 2180, 1:1000), anti-MRE11 (Abcam, Cat# ab214, 1:1000), anti-SNF2H (Novus Biologicals, Cat# NB100-55310, 1:1000), anti-YY1 (Abcam, Cat# ab109237, 1:1000), anti-Emerin (Cell Signaling, Cat# 30853S, 1:1000), anti-γH2AX (Abcam, Cat# ab2893, 1:1000), anti-53BP1 (Santa Cruz, Cat# SC-22760, 1:1000), anti-Ku70 (Santa Cruz, Cat# SC-56129, 1:1000), anti-NBS1 (Abcam, Cat# ab32074, 1:1000), anti-Rad51 (Active Motif, Cat# 39194, 1:1000), anti-histone H3 (Abcam, Cat# ab1791, 1:1000), anti-NCL (Abcam, Cat# ab22758, 1:1000), anti-Fibrillarin (Abcam, Cat# ab4566, 1:1000), anti-ActB (Cell Signaling, Cat# 4970, 1:1000), anti-C1QBP1 (Abcam, Cat# ab270033, 1:1000), and anti-BRCA1 (Novus Biologicals, Cat# NB100-404, 1:500). Membranes were washed four times with T-TBS, incubated for 1 h with secondary antibodies (HRP goat anti-mouse IgG, Abcam, Cat# ab97046, 1:10,000; HRP goat anti-rabbit IgG, Abcam, Cat# ab6721, 1:10,000), washed four times with T-TBS, and developed with a chemiluminescence kit (Advansta, Cat# K-12042-D20, CA, USA).

#### Immunofluorescence Experiments

SH-SY5Y cells (ATCC, Cat# CRL-2266) were plated on 8-well slides (Ibidi, Cat# 80826, Germany), grown to ∼80% confluence, and irradiated with 4 Gy X-rays. At the indicated times post-IR, cells were washed twice with PBS and fixed with 4% PFA in PBS (Electron Microscopy Sciences, Cat# 15710, PA, USA) for 10 min at room temperature. Cells were washed twice with PBS and twice with washing buffer (1× PBS, 0.25% BSA, 0.1% Tween-20), incubated with antigen-retrieval solution (0.1% sodium citrate, 0.1% Triton X-100), and washed three times with washing buffer. Slides were blocked in washing buffer containing 10% normal goat serum for 1 h at room temperature, then incubated overnight at 4 °C with primary antibodies diluted in blocking solution: anti-biotin (Abcam, Cat# ab53494, 1:1000), anti-myc-tag (Cell Signaling, Cat# 2276S, 1:1000), anti-Nucleolin (Abcam, Cat# ab22758, 1:1500), anti-Fibrillarin (Abcam, Cat# ab4566, 1:1000), anti-γH2AX rabbit (Abcam, Cat# ab2893, 1:1500), anti-γH2AX mouse (Millipore, Cat# 05-636, 1:1500), anti-BRCA1 (Novus Biologicals, Cat# NB100-404, 1:500), anti-53BP1 (Novus Biologicals, Cat# NB100-305, 1:500), anti-phospho-ATM (S1981) (Cell Signaling, Cat# 5883, 1:500), and anti-BrU (Sigma-Aldrich, Cat# B8438-100UL, 1:1000). For laser-induced DNA damage experiments, cells were fixed 30 min after laser irradiation and stained with anti-γH2AX (Millipore, Cat# 05-636, 1:1500), anti-NCL (Abcam, Cat# ab22758, 1:1500), and anti-CTCF (Cell Signaling, Cat# 3418, 1:1000). Slides were then washed three times with washing buffer (10 min each) and incubated for 1 h at room temperature with secondary antibodies (donkey anti-rabbit AF488, Jackson ImmunoResearch, Cat# 711-545-152; goat anti-mouse AF647, Jackson ImmunoResearch, Cat# 715-605-150; both 1:250). Slides were washed three times with washing buffer (10 min each), stained with 5 µg/mL Hoechst 33342 (Thermo, Cat# H1399, MA, USA), washed four times with PBS, and imaged using the 63× objective of an Axio Observer fluorescence microscope (Carl Zeiss Microscopy, Germany).

Images were processed in ImageJ¹⁰⁸. ROI identification, foci counting and size measurements, and fluorescence-intensity quantification were performed in CellProfiler¹⁰⁹.

#### Laser-induced DNA damage

LIDD experiments were performed as described^15^. Briefly, HEK293 cells transduced with shScr or shNcl were plated on glass-bottom 8-well slides and, 48 h after plating, sensitized with Hoechst 33342 (5 µg/mL; Thermo, Cat# H1399, MA, USA) for 10 min at 37 °C in fully supplemented DMEM without phenol red. Cells were washed twice with warm PBS and returned to phenol red-free DMEM. Laser damage was induced on an Axio Observer microscope (Carl Zeiss Microscopy, Germany) using a 40× objective and a UGA-42 Caliburn 355 nm pulsed laser (1 kHz, 42 µJ/pulse; Rapp OptoElectronic, Germany) at 3% power output and 1.5 s illumination per ROI, drawing parallel lines along the vertical axis of the field at ∼20 µm spacing. Cells were fixed with 4% PFA 30 min after irradiation and stained as described above for immunofluorescence. Fluorescence intensity and γH2AX-domain size and shape were quantified in CellProfiler v4.2.8^102^.

#### BrU analysis of de novo transcription

For de novo transcription analysis, HEK293 cells transduced with shScr, shNcl-1, or shNcl-2 were irradiated with 4 Gy X-rays and fixed with 2% PFA at 30 min, 2 h, and 8 h post-irradiation; a non-irradiated control was included for each condition. De novo transcription was metabolically labeled by adding 2 mM 5-bromouridine (BrU; Sigma-Aldrich, Cat# <ADD>, [ADD] purity) 20 min before fixation and incubating for 20 min at 37 °C. After fixation, cells were stained with anti-BrU (Sigma-Aldrich, Cat# B8438-100UL, 1:1000) and anti-NCL to identify nucleoli, following the immunofluorescence protocol described above.

#### NHEJ and HR repair efficiency analysis

HEK293 cells transduced with shScr, shNcl-1, or shNcl-2 were co-transfected with equal amounts of the I-SceI–IFP endonuclease plasmid (pRRL-sEF1α-HA.NLS.Sce(opt).T2A.IFP; Addgene #31484) and either the DR-GFP homologous-recombination reporter (pDR-GFP; Addgene #26475) or the EJ5-GFP non-homologous-end-joining reporter (pimEJ5-GFP; Addgene #44026). After 72 h, cells were harvested, fixed with 4% PFA, and analyzed on a BD FACSAria III (Becton, Dickinson and Company). GFP was detected with the 488 nm laser (502 nm long-pass, 530/60 nm band-pass) and IFP with the 640 nm laser (690 nm long-pass, 730/45 nm band-pass). Debris and doublets were excluded (FSC-A vs. SSC-A, then FSC-A vs. FSC-H and SSC-A vs. SSC-H) before gating IFP-positive (transfected) cells. Repair efficiency was calculated as 100 × (GFP⁺IFP⁺ / IFP⁺). FCS files were analyzed in Floreada.io (https://floreada.io/).

#### SPARK-ID plasmid construction

To generate the SPARK-ID plasmids, the SIRT6, Ku80, and MRE11 coding sequences were amplified from the previously constructed plasmids SIRT6-FLAG, pEGFP-C1-FLAG-Ku80, and FLAG-MRE11, respectively. SIRT6-FLAG was a gift from Eric Verdin¹¹⁰ (Addgene #13817; RRID:Addgene_13817), pEGFP-C1-FLAG-Ku80 was a gift from Steve Jackson¹¹¹ (Addgene #46958; RRID:Addgene_46958), and FLAG-MRE11 was as previously described¹¹²,¹¹³. Purified PCR products were inserted by Gibson cloning¹¹⁴ into the pcDNA3.1-mycBioID backbone, which contains the myc-tagged BioID domain (a gift from Kyle Roux²⁹; Addgene #35700; RRID:Addgene_35700). The resulting myc-BioID-SIRT6, myc-BioID-Ku80, and myc-BioID-MRE11 cassettes were PCR-amplified and subcloned into the Lenti-X-pTuner backbone, which carries the FKBP-based destabilization (DD) domain (pLVX-PTuner; Takara, Cat# 632173, Japan). These Lenti-X-DD-mycBioID-sensor constructs were used for lentiviral packaging and are referred to hereafter as Lenti-X-SPARK-ID.

#### Nucleolin silencing experiments

Two lentiviral shRNA constructs targeting Nucleolin were obtained from the Sigma-Aldrich MISSION shRNA library: shNcl-1 (TRCN0000062284, “CGGTGAAATTGATGGAAATAA”) and shNcl-2 (TRCN0000062283, “CCTTGGAAATCCGTCTAGTTA”) (Sigma-Aldrich, Germany), together with a non-targeting scrambled control (shScramble) from the same library. Both NCL-targeting shRNAs were tested in SH-SY5Y cells; shNcl-2 achieved efficient NCL knockdown and is the hairpin designated “shNcl” in the Results. shRNA plasmids were packaged into lentiviral particles and used to transduce SPARK-ID SH-SY5Y cells. Transduced cells were selected in DMEM supplemented with 10 µg/mL puromycin (InvivoGen, Cat# ant-pr-1) and 3 mg/mL Geneticin (G418; Formedium, England, Cat# G418S). NCL depletion became detectable after more than two weeks of selection but was progressively lost by ∼3–4 weeks; shNcl and shScr transductions were therefore repeated for each independent experiment. For every streptavidin-directed immunoprecipitation, NCL silencing was confirmed by Western blotting of chromatin extracts and by immunofluorescence of the same samples prior to mass-spectrometry analysis. For NCL silencing in HEK293 cells, both shNcl-1 and shNcl-2 were used.

#### Lentivirus packaging and cell transfection and transduction

Lenti-X-SPARK-ID plasmids were packaged into lentiviral particles in HEK293 cells by co-transfection with lentiviral packaging plasmids using PolyJet (SignaGen, Cat# SL100688, MD, USA)¹¹⁵. Particles were harvested and used to transduce SH-SY5Y cells, which were then selected with 10 µg/mL puromycin (InvivoGen, Cat# ant-pr-1). Stably transduced cells were used for biotin-labeling experiments. Transient transfections were performed according to the PolyJet manufacturer’s protocol.

#### X-ray irradiation and Biotin-labeling experiments

To reduce background biotinylation, biotin was depleted from the culture medium by adding BioLock biotin-blocking solution (IBA Lifesciences, Cat# 2-0205-050, Göttingen, Germany), agitating for 20 min, and filtering through a 0.2 µm unit. SPARK-ID SH-SY5Y cells were maintained in this biotin-depleted medium for at least one week before labeling. Five hours before irradiation, SPARK-ID protein expression was induced by adding Shield1 at 50 nM (SIRT6ID) or 500 nM (Ku80ID and MRE11ID). Cells were then irradiated with 4 Gy X-rays in a Faxitron MultiRad 160 cabinet (Faxitron, AZ, USA), and 50 µM biotin was applied as a 5-min pulse immediately after irradiation. Cells were washed four times with PBS, returned to biotin-depleted medium, and collected after 5 min, 30 min, 2 h, 8 h, or 24 h chase intervals. Non-irradiated controls (nonIR) were treated identically but mock-irradiated, and collected 5 min or 24 h afterwards. Negative controls were irradiated at 5 min and 24 h post-IR but received neither Shield1 nor biotin, allowing basal (Shield1/biotin-independent) biotinylation to be subtracted.

#### Streptavidin-directed Immunoprecipitations and co-immunoprecipitations

Chromatin proteins were quantified by Bradford assay, and 2 mg per condition was used for streptavidin-directed immunoprecipitation. Samples were adjusted to 1 mL with 150 mM KCl lysis buffer (150 mM KCl, 25 mM Tris pH 7.5, 5% glycerol, 0.1% Triton X-100, 1 mM DTT, 0.2 mM EDTA, 0.5 mM PMSF, 1× phosphatase-inhibitor cocktail, 10 nM TSA), and 10% (100 µL) was reserved as input. Dynabeads M-280 Streptavidin (100 µL; Thermo Fisher, Cat# 11206D, MA, USA) were added and samples were incubated with rotation at 4 °C for 2 h. Beads were then washed sequentially, all at 4 °C with rotation and bead recovery on a magnetic stand: twice (5 min each) with 150 mM KCl lysis buffer, once with 300 mM KCl lysis buffer, twice with 400 mM KCl lysis buffer, once with 300 mM KCl lysis buffer, and twice with 150 mM KCl lysis buffer (the 300 and 400 mM buffers were identical to the 150 mM buffer except for KCl concentration). Captured proteins were eluted by boiling the beads in 50 µL of 1× Laemmli buffer (62.5 mM Tris-HCl pH 6.8, 2% SDS, 10% glycerol, 5% 2-mercaptoethanol, 0.002% bromophenol blue) for 20 min; the eluate was recovered, the beads were boiled again in a further 50 µL of 1× Laemmli buffer for 10 min, and the two eluates were combined.

For FLAG-directed immunoprecipitations, SH-SY5Y SIRT6-KO cells¹¹⁶ were transfected with SIRT6-FLAG using PolyJet and, 24 h later, irradiated (4 Gy X-rays) or mock-irradiated, then collected at the same DDR time points used for the biotin co-IPs. Cells (∼15 × 10⁶) were homogenized in 1 mL of 150 mM KCl lysis buffer, incubated on ice for 30 min, and cleared by centrifugation at 17,000 × g for 20 min at 4 °C. Protein was quantified by Bradford, 1 mg was used per co-IP, and 100 µL was reserved as input. Anti-FLAG M2 magnetic beads (50 µL; Sigma, Cat# M8823, MO, USA) were added and incubated for 1 h at 4 °C with rotation. Beads were washed twice with 150 mM KCl lysis buffer, twice with 300 mM KCl lysis buffer, and once with 150 mM KCl lysis buffer, and bound proteins were eluted by rotating for 1 h at 4 °C in 150 mM KCl lysis buffer containing FLAG peptide.

For NCL immunoprecipitations, the SIRT6-FLAG protocol was followed with the modifications below. NCL AP-MS was performed in WT and SIRT6-KO SH-SY5Y cells under nonIR and IR30 conditions, with four biological replicates per condition; empty-vector anti-FLAG immunoprecipitations from the same backgrounds served as background controls. After lysis and whole-lysate normalization, 1.5 mg of total protein was incubated overnight at 4 °C with rotation (20 rpm) with anti-NCL antibody (1:200; Abcam, Cat# ab22758, RRID:AB_776878) and 50 µL of protein G magnetic beads (Invitrogen, Cat# 10004D). For NCL AP-MS, beads were washed four times with high-salt lysis buffer (0.5 M KCl, 50 mM Tris-HCl pH 7.5, 1% NP-40, 0.5 mM DTT, 0.5 mM PMSF, 1× protease- and phosphatase-inhibitor cocktails) and once with SDAC buffer (50 mM Tris-HCl pH 9.0, 4 mM MgCl₂, 50 mM NaCl, 0.5 mM DTT, 0.5 mM PMSF, 1× protease- and phosphatase-inhibitor cocktails). Proteins were eluted by boiling in 1× Laemmli buffer.

#### Mass spectrometry sample processing and analysis

For mass spectrometry analysis, first, we loaded 50 µL of each co-IP sample (one entire SPARK-ID experiment per gel) in a 12% acrylamide Tris-glycine gel. We did an electrophoretic separation of the samples, and the protein gel was stained with Coomassie blue to visualize every lane. Every lane was isolated from the gel with the help of a scalpel. The proteins in the gel lanes were reduced with 3mM DTT (60C for 30min), modified with 10mM iodoacetamide in 100mM ammonium bicarbonate (in the dark, room temperature for 30min), and digested in 10% acetonitrile and 10mM ammonium bicarbonate with modified trypsin (Promega) at a 1:10 enzyme-to-substrate ratio, overnight at 37C. Second digestion with Trypsin was done for 4 hours at 37C. The tryptic peptides were desalted using C18 tips (Homemade stage tips) for SIRT6ID and Ku80ID samples, and an Oasis HLB 96-well elution plate (Waters) for MRE11ID samples. The peptides were dried and re-suspended in 0.1% Formic acid in 2% acetonitrile.

For SIRT6ID and Ku80ID samples, the resulting peptides were analyzed by LC-MS/MS using a Q Exactive HF mass spectrometer (Thermo) fitted with a capillary HPLC (easy nLC 1200, Thermo-Fisher). The peptides were loaded in solvent A (0.1% formic acid in water) on a homemade capillary column (30 cm, 75-micron ID) packed with Reprosil C18-Aqua (Dr. Maisch GmbH, Germany). The peptide mixture was resolved with a 6 to 30% linear gradient of solvent B (80% acetonitrile with 0.1% formic acid for 60 minutes) followed by a gradient of 15 minutes of 30 to 95% and 15 minutes at 95% Solvent B at flow rates of 0.15 μl/min. Mass spectrometry was performed in positive mode (m/z 300–1800, resolution 60,000 for MS1 and 15,000 for MS2) using repetitively full MS scans followed by high-collision dissociation (HCD, at 27 normalized collision energy) of the 18 most dominant ions (>1 charge) selected from the first MS scan. The AGC settings were 3×10^6^ for the full MS and 1×10^5^ for the MS/MS scans. The intensity threshold for triggering MS/MS analysis was 8×10^4^. A dynamic exclusion list was enabled with an exclusion duration of 20s.

For MRE11ID samples, the resulting peptides were analyzed by LC-MS/MS using a Q Exactive HFX mass spectrometer (Thermo) fitted with a capillary HPLC (Ultimate 3000, Thermo Scientific). The peptides were loaded in solvent A (0.1% formic acid in water) on a homemade capillary column (30 cm, 75-micron ID) packed with Reprosil C18-Aqua (Dr. Maisch GmbH, Germany). The peptide mixture was resolved with a 5 to 28% linear gradient of solvent B (99.9% acetonitrile with 0.1% formic acid) for 120 minutes, followed by a gradient of 15 minutes from 28 to 95% and 10 minutes at 95% acetonitrile with 0.1% formic acid in water at flow rates of 0.15 μl/min. Mass spectrometry was performed in positive mode (m/z 350–1200, resolution 120,000 for MS1 and 15,000 for MS2) using repetitively full MS scans followed by high-collision dissociation (HCD, at 27 normalized collision energy) of the 20 most dominant ions (>1 charge) selected from the first MS scan. The AGC settings were 3×10^6^ for the full MS and 1×105 for the MS/MS scans. The intensity threshold for triggering MS/MS analysis was 1.3×10^5^. A dynamic exclusion list was enabled with an exclusion duration of 20s.

For the Mass Spectrometry analysis of the SPARK-ID-interactomes after shScr and shNcl treatments, the differences in sample and mass spectrometry analysis were as follows. The sample processing was done in the same way as with previous interactomes. After tryptic proteolysis of the proteins in the interactome, the tryptic peptides were desalted using Oasis HLB 96-well uElution plates (Waters) dried and re-suspended in 0.1% formic acid in 2% acetonitrile. The resulting peptides were analyzed by LC-MS/MS using a Q Exactive HFX mass spectrometer (Thermo) fitted with a capillary HPLC (Ultimate 3000, Thermo Scientific). The peptides were loaded in solvent A (0.1% formic acid in water) on a homemade capillary column (30 cm, 75-micron ID) packed with Reprosil C18-Aqua (Dr. Maisch GmbH, Germany). The peptide mixture was resolved with a 5 to 28% linear gradient of solvent B (99.9% acetonitrile with 0.1% formic acid) for 120 minutes, followed by a gradient of 15 minutes of 28 to 95% and 15 minutes at 95% solvent B at flow rates of 0.15 μl/min. Mass spectrometry was performed in positive mode (m/z 350–1200, resolution 120,000 for MS1 and 15,000 for MS2) using repetitively full MS scans followed by high-collision dissociation (HCD, at 27 normalized collision energy) of the 20 most dominant ions (>1 charge) selected from the first MS scan. The AGC settings were 3×10^6^ for the full MS and 1×105 for the MS/MS scans. The intensity threshold for triggering MS/MS analysis was 2.7×105. A dynamic exclusion list was enabled with an exclusion duration of 20 seconds.

#### Mass spectrometry raw data processing

Mass spectrometry data were processed in MaxQuant, version 1.5.2.8 (SIRT6ID and Ku80ID), 2.1.1.0 (MRE11ID)¹¹⁷, and 2.1.1.1 (shNcl experiments), using the Andromeda search engine against the human UniProt proteome, with a precursor mass tolerance of 4.5 ppm (6 ppm for the shNcl experiments) and a fragment-ion tolerance of 20 ppm. Methionine oxidation, protein N-terminal acetylation, and biotinylation were set as variable modifications, and cysteine carbamidomethylation as a fixed modification. Minimum peptide length was seven amino acids, with up to two missed cleavages. Proteins were quantified by label-free quantification (LFQ). Peptide- and protein-level false discovery rates (FDRs) were controlled at 1% using the target-decoy approach, and reverse-database hits and common contaminants were removed. Perseus v1.6.7.0¹¹⁴ was used for preliminary inspection and quality control, and MaxQuant output tables were exported for protein-level preprocessing and differential-abundance analysis in R (see Quantification and statistical analysis).

#### NCL AP-MS sample analysis

NCL AP-MS samples were processed by SDS-PAGE, in-gel reduction, alkylation, tryptic digestion, desalting, and peptide resuspension as described above. Peptides were analyzed on a Q Exactive HFX mass spectrometer coupled to an Ultimate 3000 nanoLC system. Peptides were separated using a 60-min gradient from 5% to 28% solvent B, followed by 15 min from 28% to 95% and 15 min at 95% solvent B, at a flow rate of 0.15 µL/min. MS acquisition was performed in positive-ion mode over *m/z* 300–1,500, at resolutions of 60,000 for MS1 and 15,000 for MS2. The 18 most abundant multiply charged ions were selected for HCD fragmentation at a normalized collision energy of 27. AGC targets were 3 × 10^6^ for MS1 and 1 × 10^5^ for MS2, with a maximum MS2 injection time of 60 ms, an MS/MS trigger threshold of 1.7 × 10^4^, and a dynamic-exclusion duration of 20 s.

Raw files were processed using MaxQuant v2.6.6.0 with the Andromeda search engine against the human UniProt proteome. Precursor and fragment mass tolerances were 4.5 and 20 ppm, respectively. Methionine oxidation and protein N-terminal acetylation were included as variable modifications, and cysteine carbamidomethylation as a fixed modification. A minimum peptide length of seven amino acids and a maximum of two missed cleavages were allowed. Label-free quantification was performed in MaxQuant, with peptide- and protein-level FDRs controlled at 1% using the target-decoy strategy. Reverse identifications and common contaminants were removed before downstream analysis.

### Quantification and statistical analysis

#### Proteomic preprocessing and moderated differential-abundance analysis

Protein identifications corresponding to reverse-database matches, common proteomic contaminants, and proteins supported by fewer than two peptides were removed before statistical analysis. Undetected measurements in the protein-intensity input matrices were represented by a fixed baseline intensity of 100,000; consequently, no missing-value imputation was performed during downstream analysis in R. The protein-level abundance matrices were processed using the quantify.proteins() function from the proteomics workflow described by Kammers et al.^103,104^, generating median-polished, log2-transformed, and normalized protein-abundance values. Differential abundance was then evaluated using the empirical-Bayes moderated statistics implemented in the same workflow. For each comparison, the model returned a log2 fold-change (log2FC), ordinary and moderated t-statistics, moderated p-value (p_mod), and corresponding multiple-testing-adjusted values. The empirical-Bayes variance-moderation procedure was used without modification.

#### Subcellular-localization curation

Protein groups were manually curated before and after differential-abundance analysis using UniProt subcellular-location annotations, Gene Ontology Cellular Component annotations, the Human Protein Atlas, and manual GeneCards queries. Proteins were retained only when all resources supported nuclear localization. Additional non-nuclear localization annotations did not result in exclusion when nuclear localization was concordantly supported.The strict requirement for concordant nuclear localization across three annotation resources may have excluded genuine nuclear proteins with incomplete or discordant database annotation.

#### Time-resolved SPARK-ID interactor definition

SIRT6ID, Ku80ID, and MRE11ID were analyzed independently. Each non-irradiated or post-irradiation condition contained three biological replicates and was compared with six negative-control samples using a two-group design consisting of an intercept and condition coefficient. The negative-control group comprised three controls collected with the IR5 experiment and three controls collected with the IR24 experiment. A manually curated nuclear or chromatin-associated protein was classified as a SPARK-ID interactor at a given time point when it showed positive enrichment over the negative controls with log2FC > 1.5 and p_mod < 0.05. A log2FC of 1.5 corresponds to approximately 2.8-fold enrichment. FDR-adjusted values were retained in the exported results as sensitivity metrics but were not used as the primary network-construction criterion.

This discovery-oriented criterion was selected to preserve sensitivity for network-level reconstruction in a sparse, time-resolved proximity-proteomics dataset, while requiring both a substantial effect size and moderated statistical evidence. Because nominal moderated p-values do not provide proteome-wide false-discovery control, individual protein calls were interpreted together with fold enrichment, peptide support, recurrence across experiments, nuclear-localization curation, and orthogonal validation where available.

#### SPARK-ID interactor definition after NCL silencing

Protein groups were manually curated for nuclear or chromatin-associated localization before statistical analysis using the Human Protein Atlas, GeneCards, and Gene Ontology Cellular Component annotations. The processed abundance matrices were analyzed using the same Kammers empirical-Bayes workflow applied to the original SPARK-ID time course. For each DSB sensor, shScr and shNCL samples were compared independently with the negative-control samples. Each experimental condition contained three biological replicates. Proteins showing positive enrichment over negative controls with log2FC > 1.5 and p_mod < 0.05 were classified as interactors in the corresponding shScr or shNCL network.

The same discovery-oriented criterion and interpretive constraints described above were applied to the rewiring analysis.

#### Direct shNCL-versus-shScr differential-binding analysis

To prevent proteins lacking evidence of SPARK-ID association from entering the direct perturbation analysis, the comparison universe was restricted to the union of nuclear proteins classified as interactors in either the shScr or shNCL condition relative to their negative controls. Within this curated interactor universe, direct shNCL-versus-shScr contrasts were evaluated using the same moderated limma framework. Proteins were classified as rewired when the direct contrast satisfied p_mod < 0.05 and log2FC > 1.5. Positive log2 fold-changes represented interactions gained or increased after NCL depletion, whereas negative values represented interactions lost or decreased. FDR-adjusted values were retained as sensitivity metrics but were not used as the primary rewiring criterion.

#### Physical protein–protein interaction network reconstruction

For each DSB sensor and experimental condition, the final curated SPARK-ID interactor list was used as the network node set. A physical-PPI prior was required because proximity labeling identifies proteins within the bait-associated environment but does not directly measure prey– prey interactions. Network reconstruction therefore evaluated how the experimentally identified node sets were organized within the currently supported physical-interaction literature rather than inferring new physical edges from the SPARK-ID abundance data. Physical protein–protein interactions among these proteins were retrieved from the STRING physical-interaction network32. For interactions between non-bait proteins, edges were retained when the STRING physical-interaction confidence score was ≥0.4. An exception was applied to edges directly involving the corresponding DSB-sensor bait,SIRT6, Ku80, or MRE11,for which all physical interactions represented in STRING were retained irrespective of score. This exception was introduced because several established and experimentally validated sensor interactions had unexpectedly low STRING confidence values and would otherwise have been excluded from the reconstructed networks. The original STRING experimental-evidence score was retained as the analytical edge weight, including for bait-associated edges below the 0.4 threshold. Thus, low-scoring bait interactions contributed to network connectivity but remained proportionally down-weighted in weighted analyses. The STRING combined score and complementary evidence channels, including co-expression, were retained as descriptive annotations but were not used to determine edge inclusion, network visualization, modularity, or topological classification.

The resulting networks were represented as undirected weighted graphs in which nodes corresponded to proteins experimentally identified by SPARK-ID and edges represented previously supported physical PPIs among those proteins. Proteins lacking a qualifying physical edge were retained in the complete SPARK-ID interactor tables but were excluded from graph-dependent analyses.

#### Network visualization and structural measurements

Condition-specific PPI networks were visualized in Cytoscape v3.10.2^105^. Node attributes, including SPARK-ID log2 fold-change, experimental condition, sensor membership, functional annotation, and topological classification, were mapped onto the network graphs. Basic network characteristics and node-level structural measurements were calculated using the igraph R package v2.0.3121. Only the experimentally supported physical PPI layer and its corresponding experimental-evidence weight were used for these calculations.

We used as the computational environment the R base package libraries (V4.4.0, https://www.R-project.org/).

#### Temporal network comparison

Time-point-specific networks for each DSB sensor were aligned and visualized as multiplex networks using the MuxViz framework122. Node identities and graph positions were maintained across the corresponding sensor-specific time course to facilitate visualization of temporal gains, losses, and changes in network organization. Similarity in node composition between time points was evaluated using Pearson correlation of the corresponding node-presence profiles.

#### Modularity and node classification analysis

The condition-specific SPARK-ID physical PPI edge lists were analyzed as undirected weighted networks using the rnetcarto R package v0.2.6^37–39^. Edge weights corresponded to the STRING experimental physical-interaction evidence scores described above. Network partitioning was performed using a simulated-annealing algorithm as proposed by Guimerà and Amaral 2005125,126, that searched for the node partition maximizing network modularity, thereby assigning each connected protein to a structural module. We calculated network modularity using a node partitioning into modules optimisation approach grounded on simulated annealing, in order to maximise the modularity function:

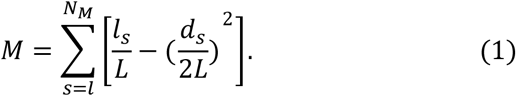

where NM is the number of modules, L is the number of links in the network, ls is the number of links between nodes in module s, and ds is the sum of the degrees (number of links to or from that node) of the nodes in module s. The partition that maximised M, determined the nodes’ module membership.

To establish the roles of proteins within networks, we calculated measures of within-module degree and participation coefficients, which quantify the degree of connectivity of each node within its own module and the rest of the network respectively (Guimerà and Amaral 2005) 42,125,126. Within-module degree is defined thus:

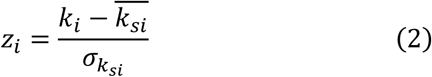

where ki is the number of links of node I to other nodes in its module si, *k_si_* is the average of k over all the nodes in si, and *σ_ksi_* is the standard deviation of k in si. The participation coefficient quantifies how broadly a node distributes its interactions across modules:

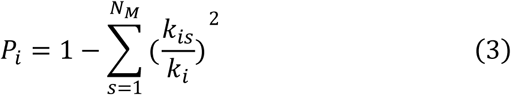

where kis is the number of links of node I to nodes in module s, and ki is the total degree if node i. Large values of Pi (close to 1) indicate that node I has its links uniformly distributed across all modules in the network, whereas Pi close to 0 indicates that all its links fall within the module it belongs to. The relationship between these two quantities defines the role of the nodes (i.e. proteins) within their respective networks (Figure 4A). We calculated modularity M, the corresponding modular partition of nodes (modules), zi’s and Pi’s using the rnetcarto library in R (V0.2.6, https://cran.r-project.org/web/packages/rnetcarto/index.html).

Topological roles were assigned from the *z_i_*–*P_i_* cartography generated by rnetcarto. The package classifies connected nodes as Ultra peripheral, Peripheral, Connector, Kinless, Peripheral Hub, Connector Hub, or Kinless Hub. In the manuscript figures and downstream analyses, these package-derived role labels were retained; briefly, using these parameters we classified the nodes into peripheral nodes (Zi<2.5, and Pi<0.6), module hubs (Zi>2.5), network hubs (Zi>2.5, and Pi>0.6), and connector nodes (Zi<2.5, and Pi>0.6)42,125; Proteins without a qualifying physical edge, and therefore without calculable module or cartographic measurements, were classified as disconnected or unclassified. Module numbers were generated algorithmically and had no intrinsic biological meaning. Biological module descriptions were assigned manually after examining the dominant functional annotations and protein composition of each module. The role classification refers to the position and connectivity of each node and allows us to rank the nodes based on the configuration of their interactions.

Structural modules were identified computationally without using functional annotations. After module detection, modules were manually described according to the predominant protein functions represented within each partition, including DNA/chromatin metabolism, transcription/RNA metabolism, ribosome biogenesis, nuclear transport, and proteostasis. These descriptive labels were used for biological interpretation and visualization but did not influence module detection, node connectivity, participation coefficients, or role assignment.

#### Fuzzy-C means clustering analysis

Temporal interaction profiles were analyzed independently for SIRT6ID, Ku80ID, and MRE11ID using the Mfuzz Bioconductor package (V4.4)123. For each sensor, the input matrix contained aggregated SPARK-ID interactors as rows and seven ordered experimental conditions,nonIR5, IR5, IR30, IR2h, IR8, IR24, and nonIR24,as columns. Matrix entries consisted of preassembled interaction fold-change values relative to the corresponding negative controls. Values recorded as zero in the input matrices were replaced by 1, representing no change, before clustering.

Protein trajectories containing more than 25% missing values were excluded using filter.NA(), and any remaining missing values were filled by K-nearest-neighbor imputation using fill.NA(mode = “knn”). Because the analyzed matrices contained no residual missing measurements, this imputation step served as a computational safeguard. Profiles without temporal variance were removed using filter.std(min.std = 0). Each remaining protein trajectory was standardized using Mfuzz::standardise(), resulting in a mean of zero and a standard deviation of one across conditions.

The fuzzifier parameter was estimated independently for each interactome using mestimate(). Candidate cluster solutions were evaluated using cselection() with repeated clustering runs, together with examination of empty clusters and separation of temporal profiles. The final analyses used five clusters for SIRT6ID with *m* = 1.638145, four clusters for Ku80ID with *m* = 1.648535, and six clusters for MRE11ID with *m* = 1.627608.

Mfuzz returned a primary cluster assignment and a membership value for each protein in every cluster. Proteins were assigned to the cluster reported by cl1$cluster, corresponding to their strongest fuzzy membership, without imposing an additional minimum-membership threshold. Complete membership matrices, cluster centroids, and cluster sizes were exported. Primary cluster assignments were subsequently incorporated manually into the expanded SPARK-ID node tables. Cluster numbering was sensor-specific and did not imply equivalence between similarly numbered clusters from different interactomes; comparable temporal patterns were identified by examining their centroid trajectories.

Line plots accompanying the fuzzy C-means analysis were generated from the standardized sensor-specific interaction trajectories. Values represent within-protein standardized changes across the seven experimental conditions and therefore describe the relative temporal profile of each protein rather than absolute differences in abundance between proteins. Proteins were grouped according to their primary FCMC assignment, and fuzzy membership values were displayed to indicate assignment strength.

#### Gene Ontology Biological Process enrichment analysis

Gene Ontology Biological Process enrichment was evaluated using ClueGO v2.5.10 within Cytoscape v3.10.2120. Curated SPARK-ID protein lists were submitted as discrete groups according to DSB sensor, experimental condition, or fuzzy C-means cluster. The complete set of human genes available in the ClueGO annotation database was used as the reference background.

Over-representation was evaluated using a hypergeometric test, followed by Benjamini– Hochberg correction for multiple testing. GO Biological Process terms spanning ontology levels 6–12 were included. Terms with FDR < 0.01 were retained for the global and time-resolved interactome analyses, whereas FDR < 0.05 was used for analyses of selected fuzzy C-means clusters and the IR2h- and IR8h-associated protein sets.

Functionally related GO terms were organized using a ClueGO kappa-score threshold of 0.4, which connects terms sharing annotated proteins. Redundant terms were fused according to their percentage similarity using the ClueGO term-fusion option. Kappa-based network construction and term fusion were used to reduce annotation redundancy and facilitate visualization but did not alter the underlying hypergeometric enrichment statistics. Enriched terms were subsequently grouped and colored manually according to broader biological themes, including DNA repair and synthesis, chromatin and transcriptional regulation, RNA metabolism, ribosome biogenesis, nucleocytoplasmic transport, proteostasis, telomere maintenance, and cell-cycle regulation.

Because enrichment was evaluated against the complete human-gene background rather than the experimentally detected proteome, pathway estimates may partly reflect differences in protein detectability or annotation density.

#### Manual broad functional annotation

Each SPARK-ID interactor was manually assigned to one predominant broad functional category by reviewing its Gene Ontology Biological Process annotations and consolidating related terms into a common biological theme. The ten categories were Cell-cycle regulation, Chromatin regulation or organization, DNA repair and metabolism, Nuclear pore, Proteasome, RNA metabolism, Ribosome biogenesis, Transcriptional regulation, Unfolded-protein response, and p53 pathway.

A single predominant category was assigned to each protein to provide a consistent node-level annotation across the SIRT6ID, Ku80ID, and MRE11ID networks. When a protein participated in several processes, its category was selected according to the function considered most representative in the nuclear and DNA-damage-response context. The assignments were used for network coloring, visualization, functional-composition summaries, and subsequent integrative analyses. They did not determine interactor inclusion, STRING edge selection, network modularity, or GO-term enrichment significance.

#### DNA-repair pathway annotation

To identify SPARK-ID interactors with previously reported roles in genome maintenance, protein lists from published genome-wide and high-throughput DNA-repair screens were manually curated and harmonized by gene symbol^30–36^. These reference datasets included screens identifying genes required for resistance to genotoxic agents, maintenance of genome stability, or activity of specific DNA-repair pathways. The curated gene lists were intersected with the SIRT6ID, Ku80ID, and MRE11ID interactors.

Repair-associated proteins were classified into nine categories: general double-strand-break repair (DSB), homologous recombination (HR), non-homologous end joining (NHEJ), single-strand-break repair (SSB), mismatch repair (MMR), nucleotide-excision repair (NER), base-excision repair (BER), replication-fork stalling or protection (ForkStall), and R-loop regulation (RLOOPS). Proteins not represented in the curated repair-screen lists were designated as having no screen-derived DNA-repair annotation.

For each DSB-sensor interactome and repair category, pathway coverage was calculated as the number of SPARK-ID interactors within an specific time point assigned to that category divided by the total number of SPARK-ID unique proteins in the corresponding curated reference list :

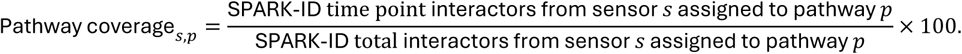

The DNA-repair annotation was independent of the broad functional category: for example, a protein could carry a broad annotation of Chromatin Organization while also being classified as an HR factor based on the repair screens.

#### CORUM protein-complex over-representation analysis

Aggregated SIRT6ID, Ku80ID, and MRE11ID interactor lists were submitted independently to the Enrichr web application and analyzed against the CORUM (version 3.0) protein-complex gene-set library^110^. Enrichr evaluates overlap between the submitted gene list and each database gene set using Fisher’s exact test and additionally reports rank-based and combined enrichment scores. Statistical significance in this study was evaluated using the Enrichr-reported multiple-testing-adjusted p-values rather than the combined score. Complexes with an adjusted p-value below 0.05 were considered significantly over-represented.

To examine the macromolecular assemblies associated with the shared connector proteins NCL and NPM1, the complete significant CORUM result tables were searched for complexes annotated as containing either protein. Selected complexes, including the Nop56p, DGCR8, and TLE1 complexes, were then mapped back onto the SIRT6ID, Ku80ID, and MRE11ID interactor sets and condition-specific physical PPI networks. The number and identity of detected subunits were evaluated across sensors and DDR conditions to describe the temporal representation of each complex within the SPARK-ID networks. These analyses used the enrichment statistics calculated in the initial global CORUM analysis and did not constitute an additional hypothesis-testing step.

For each displayed complex, the number of CORUM-annotated subunits detected in the corresponding SPARK-ID interactome was compared with the total number of annotated subunits in that complex. The overlap count and total annotated complex size were displayed together, while bar color represented the −log_10_-transformed adjusted p-value. Significantly represented complexes were subsequently grouped manually according to their predominant biological functions for visualization. The global supplementary analysis showed CORUM complexes associated primarily with chromatin regulation, RNA metabolism, ribosome biogenesis, proteostasis, and nuclear transport.

Detection or enrichment of multiple proteins belonging to a CORUM complex was interpreted as representation of complex components within the SPARK-ID interactome, not as evidence that the complete intact biochemical complex was assembled or directly associated with the sensor.

#### Reconstruction of SPARK-ID networks after NCL silencing

SIRT6ID, Ku80ID, and MRE11ID SH-SY5Y cells transduced with either scrambled control shRNA (shScr) or NCL-targeting shRNA (shNCL) were irradiated with 4 Gy and analyzed at 2 h post-irradiation. Each sensor and treatment condition contained three biological replicates. NCL depletion was confirmed by immunoblotting and immunofluorescence before the corresponding streptavidin-purified samples were analyzed by mass spectrometry.

Protein abundances were processed using the same Kammers empirical-Bayes workflow and nuclear-localization curation applied to the original SPARK-ID time course. For each DSB sensor, the shScr and shNCL samples were compared independently with the negative-control samples. A protein was classified as an interactor in the corresponding condition when it showed positive enrichment over the negative controls with log2FC > 1.5 and p_mod < 0.05.

Condition-specific SIRT6ID, Ku80ID, and MRE11ID physical PPI networks were reconstructed using the same STRING physical-interaction procedure applied to the original SPARK-ID networks. Interactions between non-bait proteins were retained at a STRING physical-interaction confidence score of ≥0.4, whereas all physical STRING edges directly involving the corresponding sensor bait were retained irrespective of score. Networks were represented as undirected weighted graphs, with the STRING experimental physical-interaction score (Weight 1) used as the analytical edge weight. Duplicate edges and self-loops were removed. Proteins lacking a qualifying physical edge were retained in the complete node tables but were excluded from graph-dependent calculations.

A complete union network containing proteins represented in either the shScr or shNCL condition was constructed to establish a common node layout and facilitate direct visualization of treatment-dependent network composition. The same node positions were subsequently used for the corresponding shScr and shNCL networks. The network script explicitly constructed this full union network before generating the individual treatment graphs. The individual SIRT6ID, Ku80ID, and MRE11ID shScr and shNCL networks were then generated as separate undirected graphs using Weight 1, with isolated nodes removed from the graph objects used for structural analysis.

#### Presence-based comparison of shScr and shNCL interactomes

For descriptive network comparisons, proteins were classified according to whether they met the interactor criteria relative to negative controls in the shScr condition, the shNCL condition, or both. Proteins detected only in shScr were designated shScr-specific or lost after NCL depletion; proteins detected only in shNCL were designated shNCL-specific or gained after NCL depletion; and proteins satisfying the interactor criteria in both conditions were designated shared. These categories describe threshold-dependent interactor-set membership and were used for integrated network visualization and interactor-count comparisons.

#### Direct shNCL-versus-shScr differential-binding analysis

Direct effects of NCL depletion were estimated independently for SIRT6ID, Ku80ID, and MRE11ID by comparing three shNCL and three shScr biological replicates using the same empirical-Bayes moderated-statistics workflow. Positive log2 fold-changes indicated increased recovery after NCL depletion, whereas negative log2 fold-changes indicated reduced recovery.

For downstream biological interpretation, the direct contrast was restricted to the union of manually curated nuclear proteins independently classified as interactors relative to negative controls in either the shScr or shNCL condition. Within this negative-control-supported interactor universe, proteins were classified as significantly rewired when the direct contrast satisfied p_mod < 0.05 and log2FC > 1.5. FDR-adjusted values were retained as sensitivity metrics but were not used as the primary rewiring criterion.

#### Detection of densely connected subnetworks using MCODE

For each DSB sensor, the union physical PPI network containing proteins represented in either the shScr or shNCL condition was imported into Cytoscape and analyzed with MCODE127 to identify densely connected subnetworks. Proteins were subsequently colored according to whether they were detected only in shScr, only in shNCL, or in both conditions. High-scoring subnetworks associated with nucleolar organization, RNA splicing, nuclear-pore function, and proteasome activity were selected for visualization based on their protein composition. MCODE outputs were interpreted as densely connected PPI subnetworks and not as proof of intact biochemical complex assembly or disassembly. Complete network settings and MCODE results are provided in the accompanying Cytoscape session file.

Physical PPI networks from the shScr and shNCL experiments were imported into Cytoscape and analyzed using the MCODE application to identify densely connected regions of the network. MCODE scores nodes according to the local density of their neighborhoods and expands candidate subnetworks from high-scoring seed nodes. The resulting groups were interpreted as densely connected PPI subnetworks rather than as evidence of intact biochemical complex formation.

#### NCL AP-MS definition and integration with SPARK-ID networks

NCL affinity-purification mass spectrometry was performed in WT SH-SY5Y cells under non-irradiated and IR30 conditions using four biological replicates per condition. NCL immunoprecipitations were compared with empty-vector anti-FLAG controls from the same cellular background. Proteins were classified as NCL AP-MS-associated when they were enriched over FLAG-empty controls in either condition with log2FC > 1.5 and moderated p_mod < 0.05, detected in at least three of four replicates in the qualifying condition, and supported by at least two razor/unique peptides. FDR values were retained as sensitivity annotations. Because the controls were generated using a different immunoprecipitating antibody, the resulting set was considered an orthogonal NCL-associated dataset rather than a definitive NCL interactome.

Candidate definition was performed at the gene level. When multiple MaxQuant protein groups represented the same gene, the maximum peptide support was retained. Indistinguishable protein groups were subsequently represented as single AP-MS features for overlap counting, heatmaps, and network visualization, while a feature-to-gene map was maintained for integration with SPARK-ID annotations. Immunoglobulin-derived features were removed, and NCL was excluded from prey-set overlaps and formal enrichment tests because it was the AP-MS bait.

For quantitative visualization, WT NCL-IP intensities were log2-transformed and normalized by subtracting the corresponding NCL bait intensity from each prey measurement. Samples were median-centered for heatmap visualization, and protein profiles were row-standardized across WT nonIR and WT IR30 replicates. NCL AP-MS-associated features were mapped to the aggregated SIRT6ID, Ku80ID, and MRE11ID interactomes using harmonized gene symbols. Feature-level overlap was visualized by UpSet analysis, and the functional composition of overlapping features was summarized using the broad SPARK-ID annotation categories.

To evaluate whether NCL-associated proteins preferentially occupied specific SPARK-ID network positions, proteins were classified as connectors or hubs when they received the corresponding rnetcarto role in at least one sensor network or time point. Enrichment of NCL AP-MS recovery within Connector, Module Hub, Connector and Hub, and Other SPARK-ID classes was evaluated using two-sided Fisher’s exact tests, followed by Benjamini–Hochberg correction. The primary analysis included only AP-MS features mapping unambiguously to one gene and excluded NCL; a sensitivity analysis included all mapped genes from ambiguous features.

NCL AP-MS-associated connector and hub features overlapping SPARK-ID were exported to Cytoscape together with separate NCL, SIRT6, Ku80, and MRE11 bait nodes. Edges represented support by the corresponding AP-MS or SPARK-ID dataset and were not interpreted as STRING physical interactions or direct binding.

Irradiation-dependent changes were assessed exploratorily by comparing WT IR30 with WT nonIR using NCL-normalized intensities. Limma models included condition and biological replicate and were fitted to proteins detected in at least two of four replicates in both conditions with at least two razor/unique peptides. Robust empirical-Bayes moderation with intensity-trend estimation was applied, and canonical FLAG-controlled candidate status was added only after model fitting.

#### Heat map plots

The heat maps were done using the ComplexHeatmap R package (V4.4)124).

#### Venn Sets Diagrams

The Venn set diagrams were done using the presence of the proteins in any given time point or fuzzy-C means cluster to generate the sets of shared and unique proteins. This analysis was computed using the nVenn R package (V0.2.2)41. The Venn cluster per time point was used as an additional sharedness annotation for the list of interactors in further analysis.

#### Statistical analysis of biochemical and cell-based assays

Western-blot, immunofluorescence, and flow-cytometry data were analyzed using the rstatix R package or GraphPad Prism. Exact statistical tests, multiple-comparison corrections, sample sizes, and definitions of biological replication are reported in the corresponding figure legends. Unless otherwise stated, tests were two-sided and p < 0.05 was considered significant.

## Additional resources

### Code availability

The R scripts used for the analysis of the SPARK-ID interactomes are available for public query and usage in the Github repository: https://github.com/agarciavenzor/SPARK-ID_Data_Analysis. The R scripts were divided into sections that follows the same order as the results sections of this manuscript.

### Results availability

The interactome results described in this paper are broadly summarised on the webpage https://sparkid.bgu.ac.il/. The website contains all the results from the mass spec analysis and the network analysis, and it is available for public query. The information can be accessed by searching for specific interactors by name or Uniprot ID, or by searching by functional annotation group. Each section displayed contains a brief description to help the user to understand the meaning of each measurement. Furthermore, the website allows downloading the information as tables, and the networks as pictures.

Finally, the Mass Spectrometry raw data is available for public access in the PRIDE repository, with the data identifier: PXD066947.

## Supplementary Figures titles and legends

**Supplementary Figure 1.**
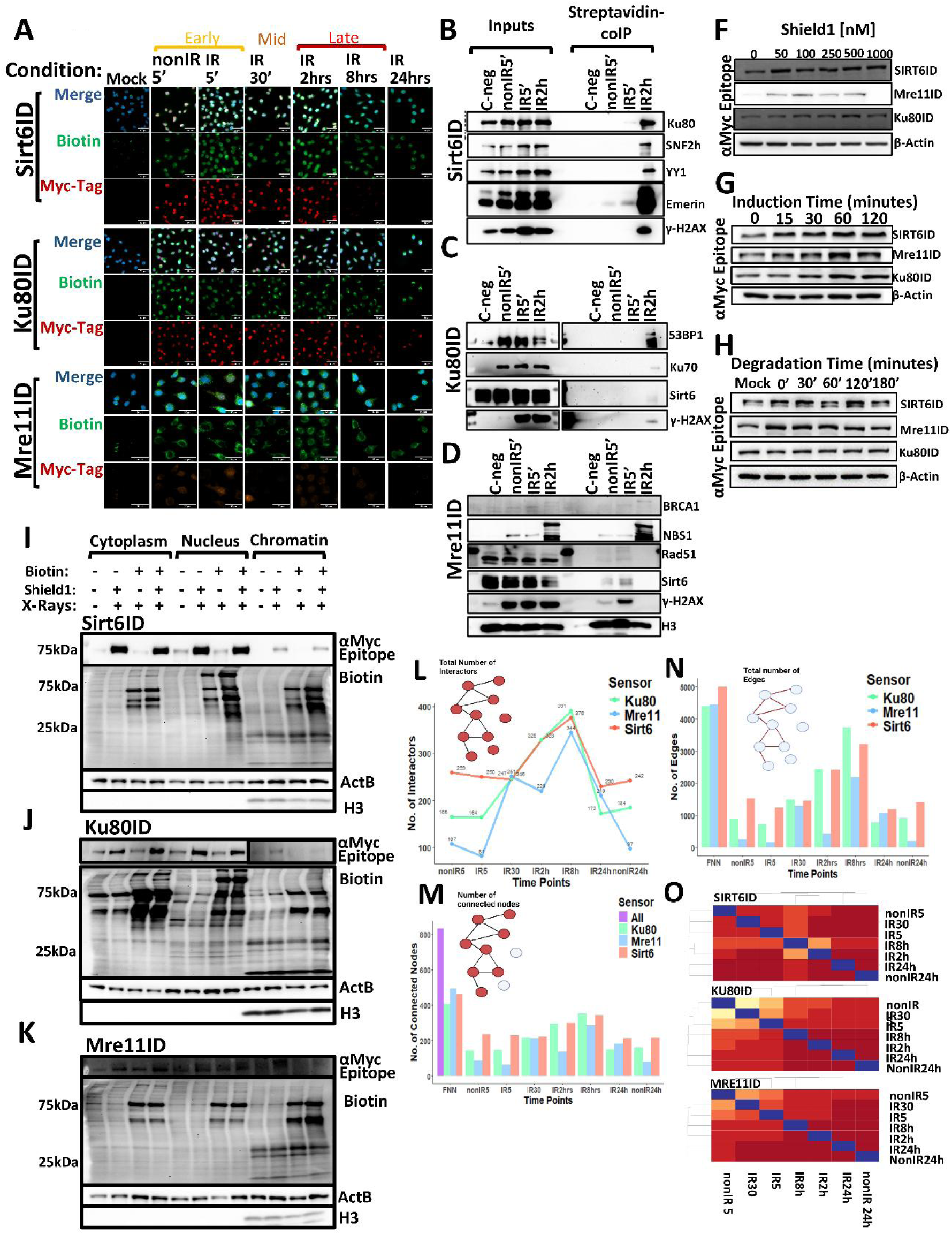
Validation and Quality Control of SPARK-ID systems. **A)** Immunofluorescence showing nuclear biotinylation and rapid SPARK-ID degradation upon Shield1 removal, done in SH-SY5Y cells. **B–D)** Co-IP validation confirming SPARK-ID interaction with known partners (e.g., Ku70, NBS1, γH2AX). **F–H)** Optimization of Shield1 dosage and induction/degradation kinetics. **I–K)** Subcellular fractionation confirming SPARK-ID recruitment to chromatin, and biotinylation of nuclear and chromatin-bound proteins. **L–O)** Number of interactors and interactions in the SPARK-ID networks: **L)** Total interactors identified per timepoint (limma, p<0.05, FC>1.5); **M)** Total number of connected nodes; **N)** Total number of interactions (edges). **O)** Pearson correlations of network node composition across timepoints, showing maximal divergence at 2 h and 8 h.

**Supplementary Figure 2.**
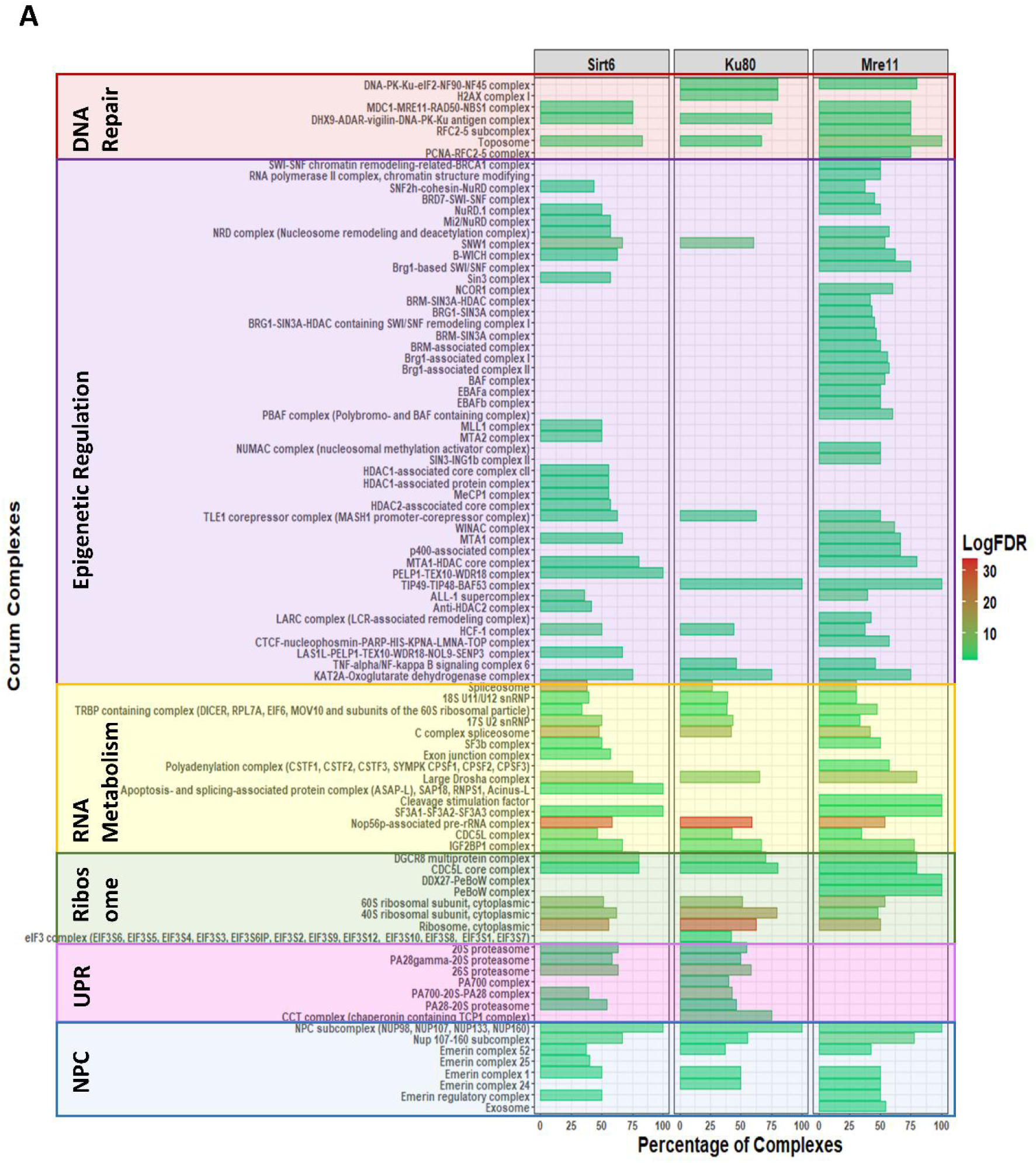
Enrichment of specific protein complexes. **A)** Enrichment of CORUM-annotated protein complexes within the SPARK-ID networks. Bar plots display the number of identified components (Dataset) versus total complex size (Background) for SIRT6ID, Ku80ID, and MRE11ID. Bars are colored by significance (-log10 FDR); all shown are significant (FDR < 0.05).

**Supplementary Figure 3.**
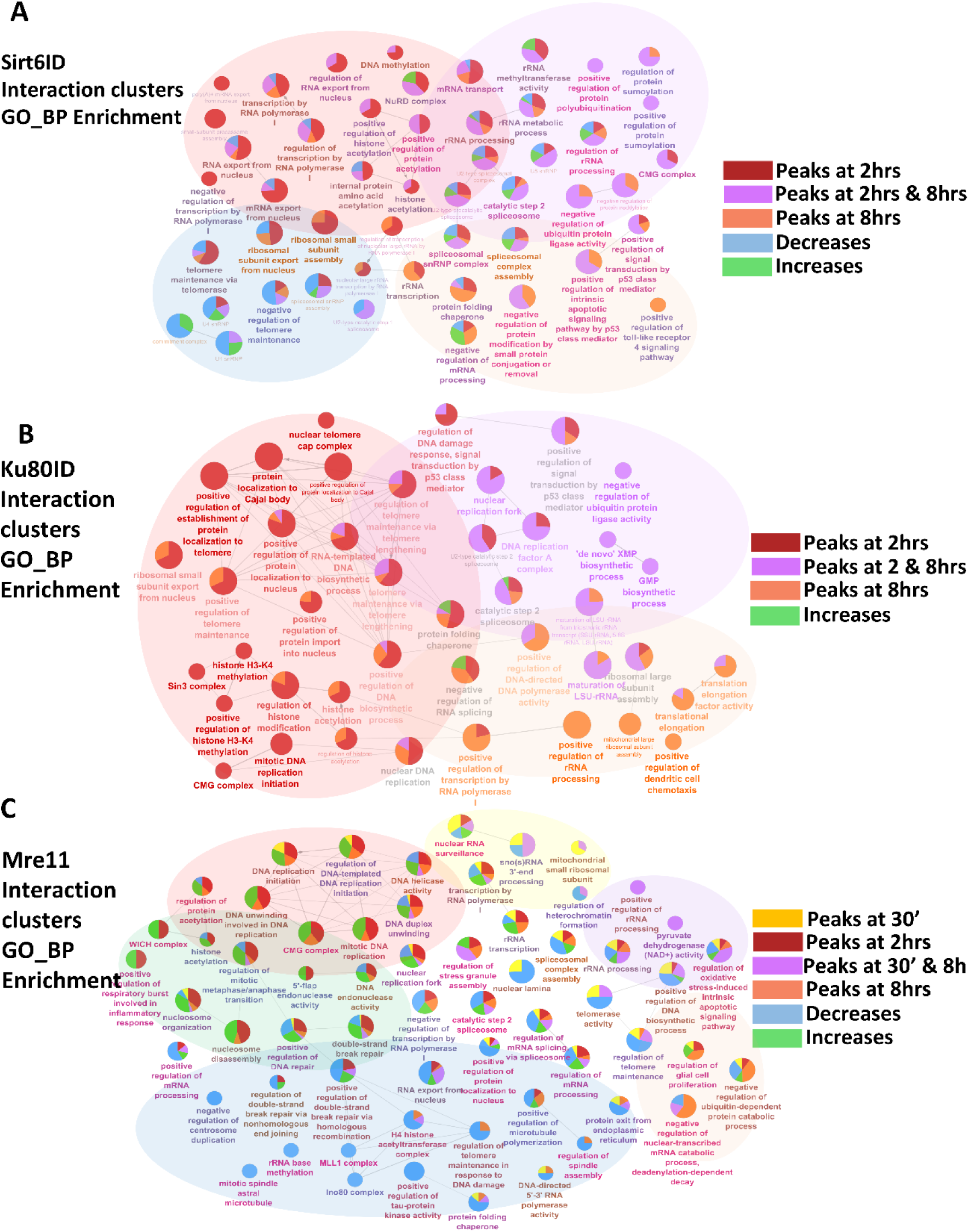
Temporal functional transitions of DSB sensors. ClueGO network maps visualizing the functional evolution of **(A)** SIRT6ID, **(B)** Ku80ID, and **(C)** MRE11ID based on Fuzzy C-Means clusters. Nodes represent enriched GO terms (FDR < 0.05); node size indicates significance. Pie charts within nodes indicate the contribution of specific temporal clusters to that function, highlighting the shift from early divergence (repair/replication) to late convergence (homeostasis/ribosome biogenesis).

**Supplementary Figure 4.**
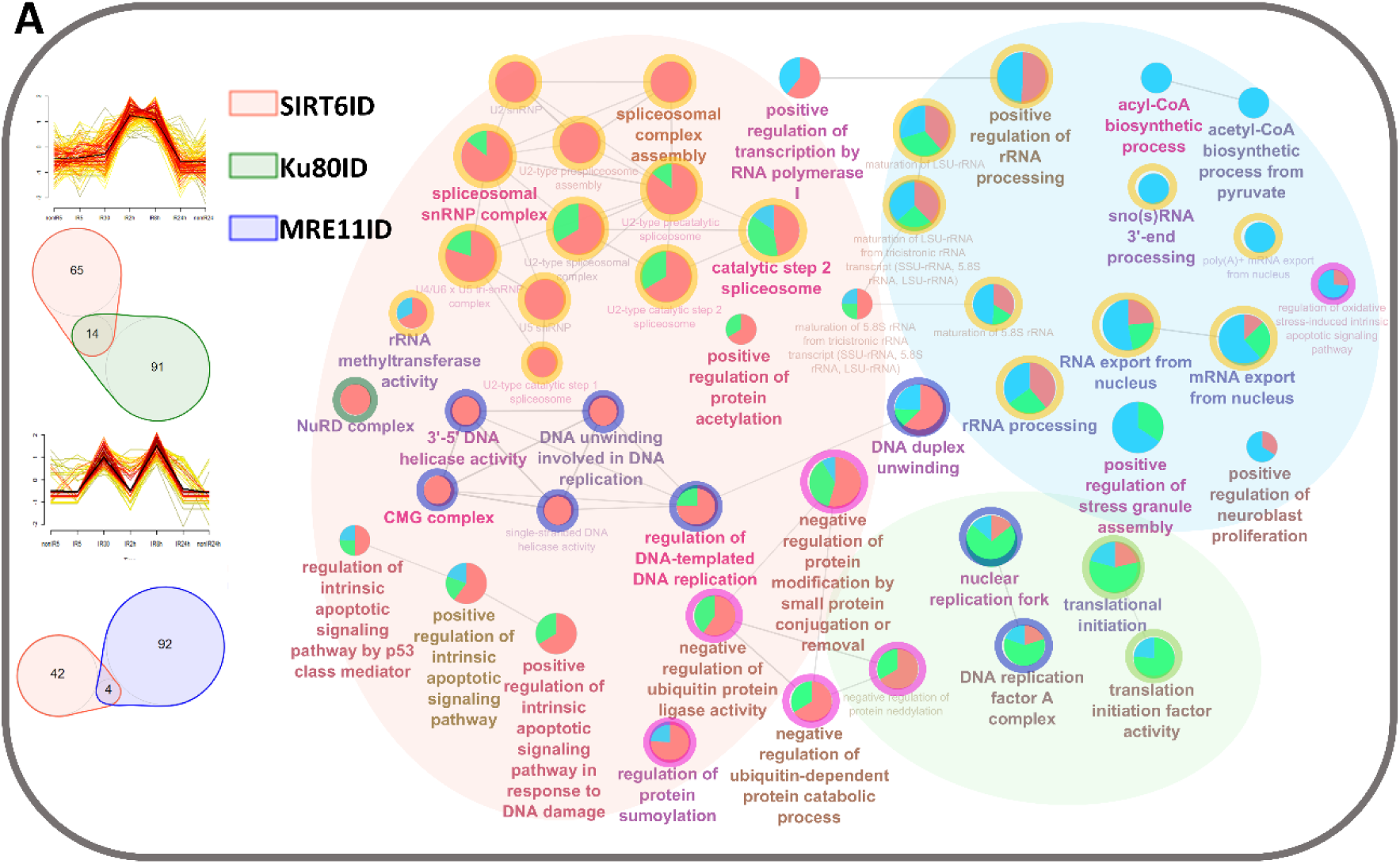
Sustained interaction clusters support continuous DNA and RNA metabolism during DDR progression. Gene Ontology Biological Process enrichment of double-peak Fuzzy C-means clusters. Nodes represent enriched GO terms, and node size indicates significance. Colors indicate manual functional grouping: RNA metabolism, DNA synthesis/unwinding, and telomere maintenance. SIRT6ID and Ku80ID sustained DNA replication-associated functions, whereas MRE11ID sustained RNA-processing functions during DDR progression.

**Supplementary Figure 5.**
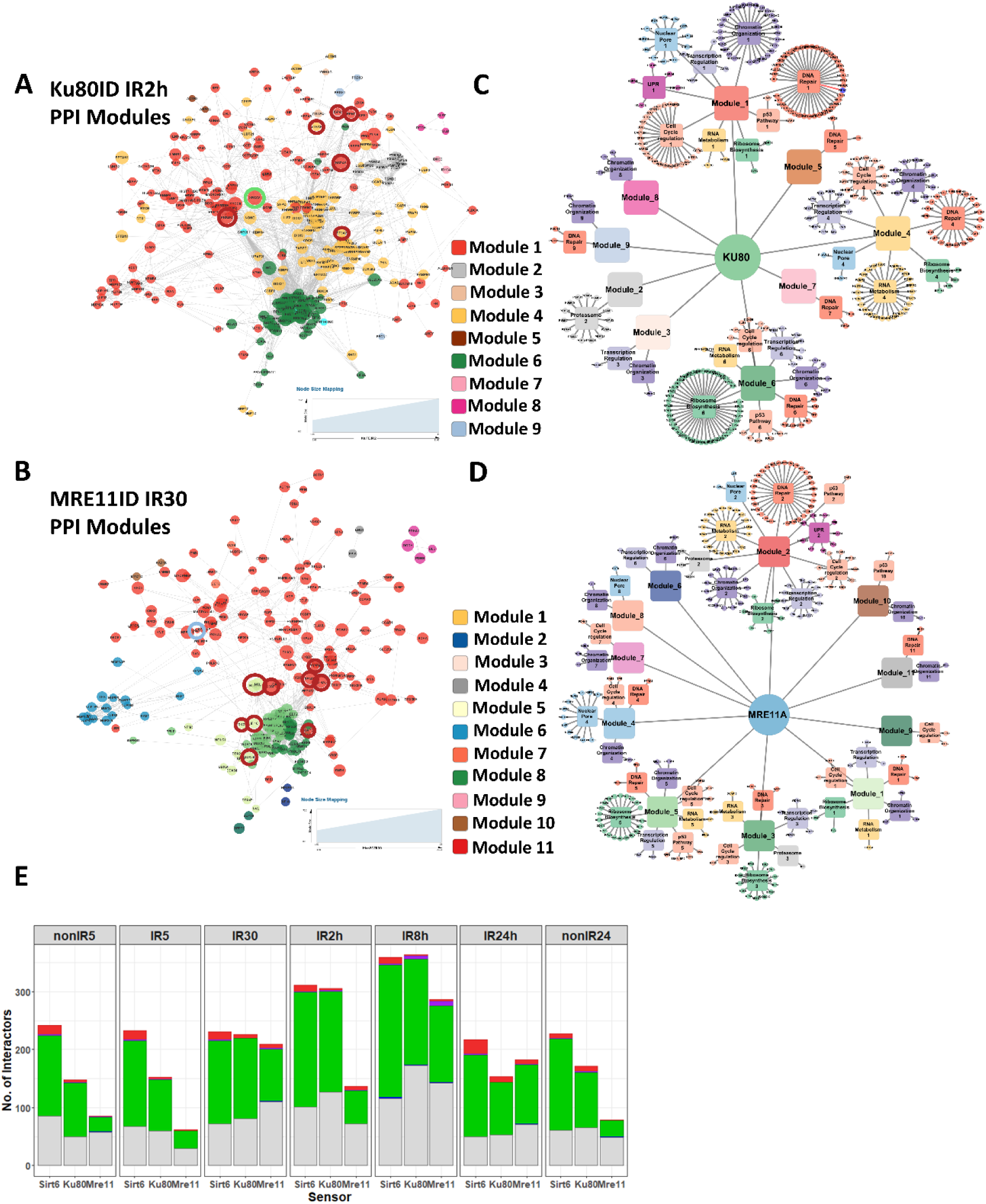
Modular architecture and topological remodeling of Ku80ID and MRE11ID networks. A-B) Modular network maps for **A)** Ku80ID at 2 h post-irradiation and **B)** MRE11ID at 30 min post-irradiation. Nodes are colored by structural module, and connector proteins are highlighted with red outlines. **C-D)** Detailed functional composition of **C)** Ku80ID and **D)** MRE11ID structural modules. Dominant module functions and corresponding proteins are indicated. **E)** Temporal dynamics of topological node classes across DSB-sensor networks. Bar plots quantify connector, module-hub, peripheral-hub, peripheral, ultra-peripheral, and disconnected nodes across DDR stages.

**Supplementary Figure 6.**
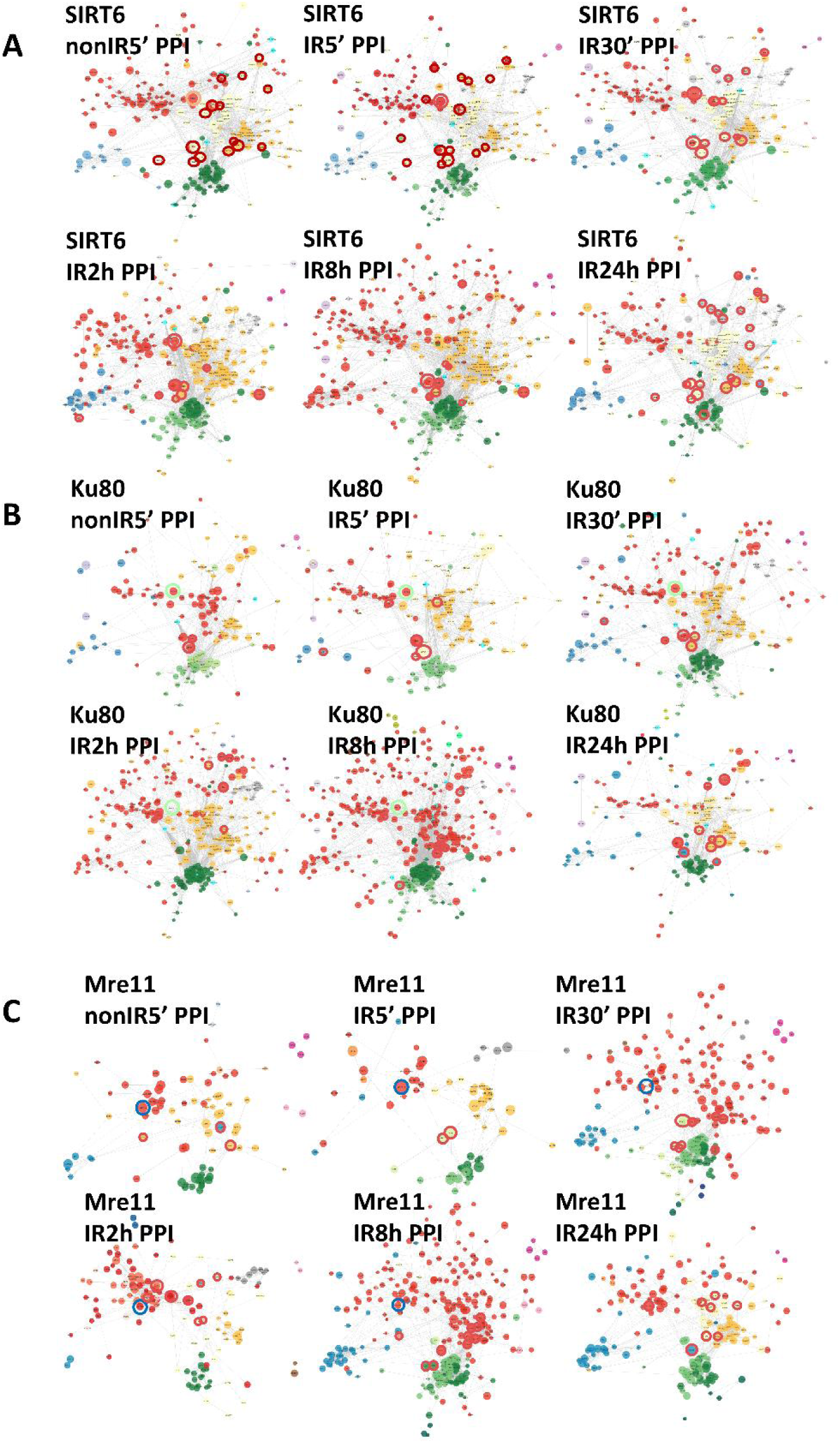
Temporal evolution of DSB-sensor network topology during DDR progression. A-C) Modular architecture of **A)** SIRT6ID, **B)** Ku80ID, and **C)** MRE11ID networks across DDR time points. Node positions are maintained within each sensor-specific time course to visualize network remodeling. Nodes are colored according to structural module, node size corresponds to interaction fold-change, and connector proteins are highlighted with red halos. The networks show temporal changes in module composition, connectivity, and connector positioning during repair progression.

**Supplementary Figure 7.**
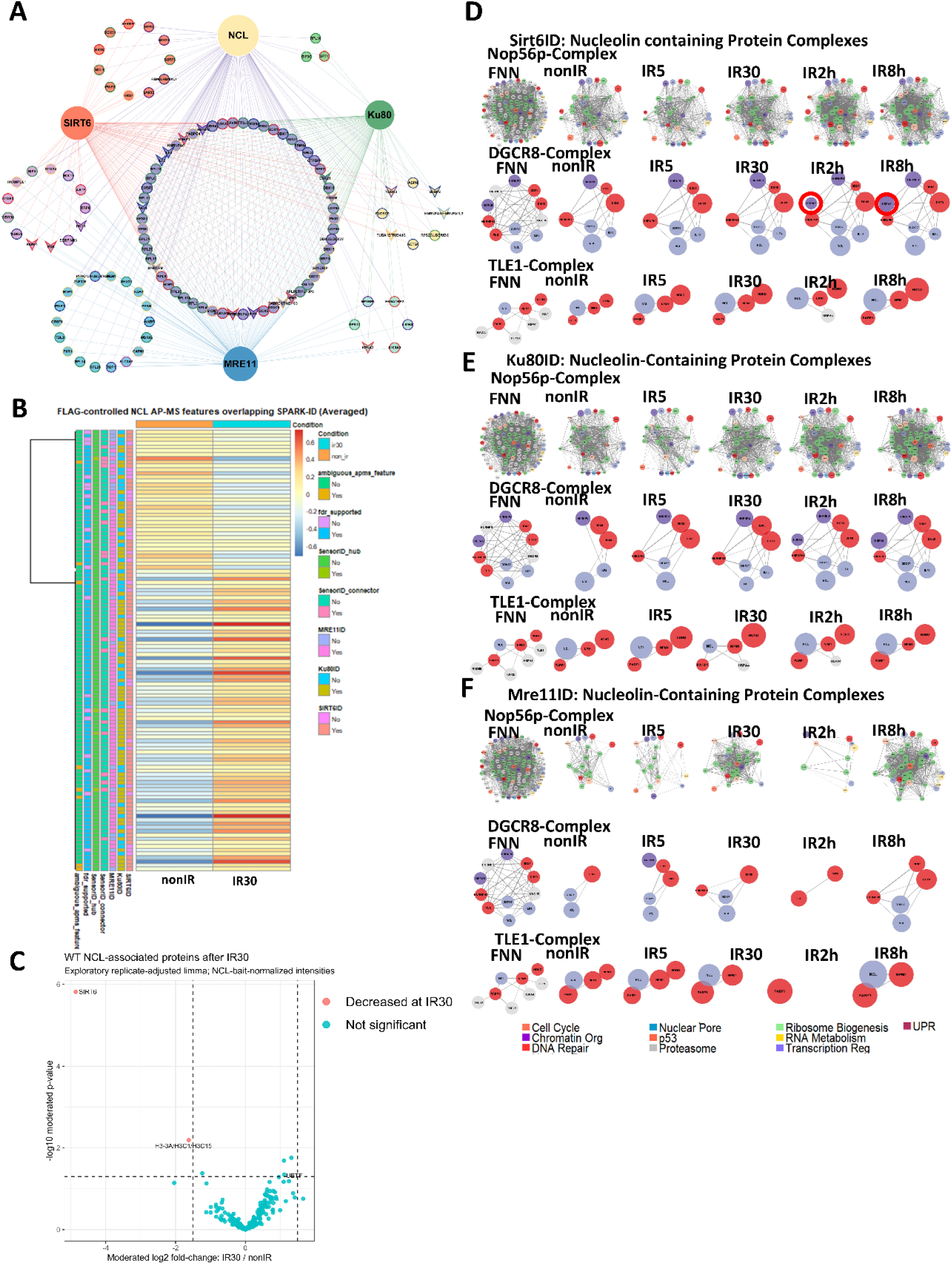
FLAG-controlled NCL AP-MS-associated proteins overlap SPARK-ID networks and RNA/nucleolar complexes. **A)** Network showing FLAG-controlled NCL AP-MS-associated protein features shared with the SIRT6ID, Ku80ID, and MRE11ID networks. Node color indicates dataset membership, node shape indicates topological class, and edge color indicates the supporting NCL AP-MS or SPARK-ID dataset. **B)** Heatmap of all FLAG-controlled NCL AP-MS-associated features overlapping SPARK-ID networks under WT nonIR and WT IR30 conditions. Values represent row-scaled, NCL-bait-normalized log2 MS intensities averaged across four biological replicates. Side annotations indicate SPARK-ID membership, topology, ambiguous protein-group mapping, and FDR support. **C)** Exploratory volcano plot of condition-dependent changes in the NCL-associated protein environment at IR30 relative to nonIR. Moderated limma statistics were calculated from NCL-bait-normalized intensities using a model that included biological replicate. Dashed lines indicate p_mod = 0.05 and |log2FC| = 1.5. Indistinguishable protein groups were represented as single AP-MS features. **D-F)** SPARK-ID subnetworks showing DDR-dependent representation of CORUM-annotated NCL/NPM1-containing complexes, including the Nop56p, DGCR8, and TLE1 complexes, in **D)** SIRT6ID, **E)** Ku80ID, and **F)** MRE11ID networks.

**Supplementary Figure 8.**
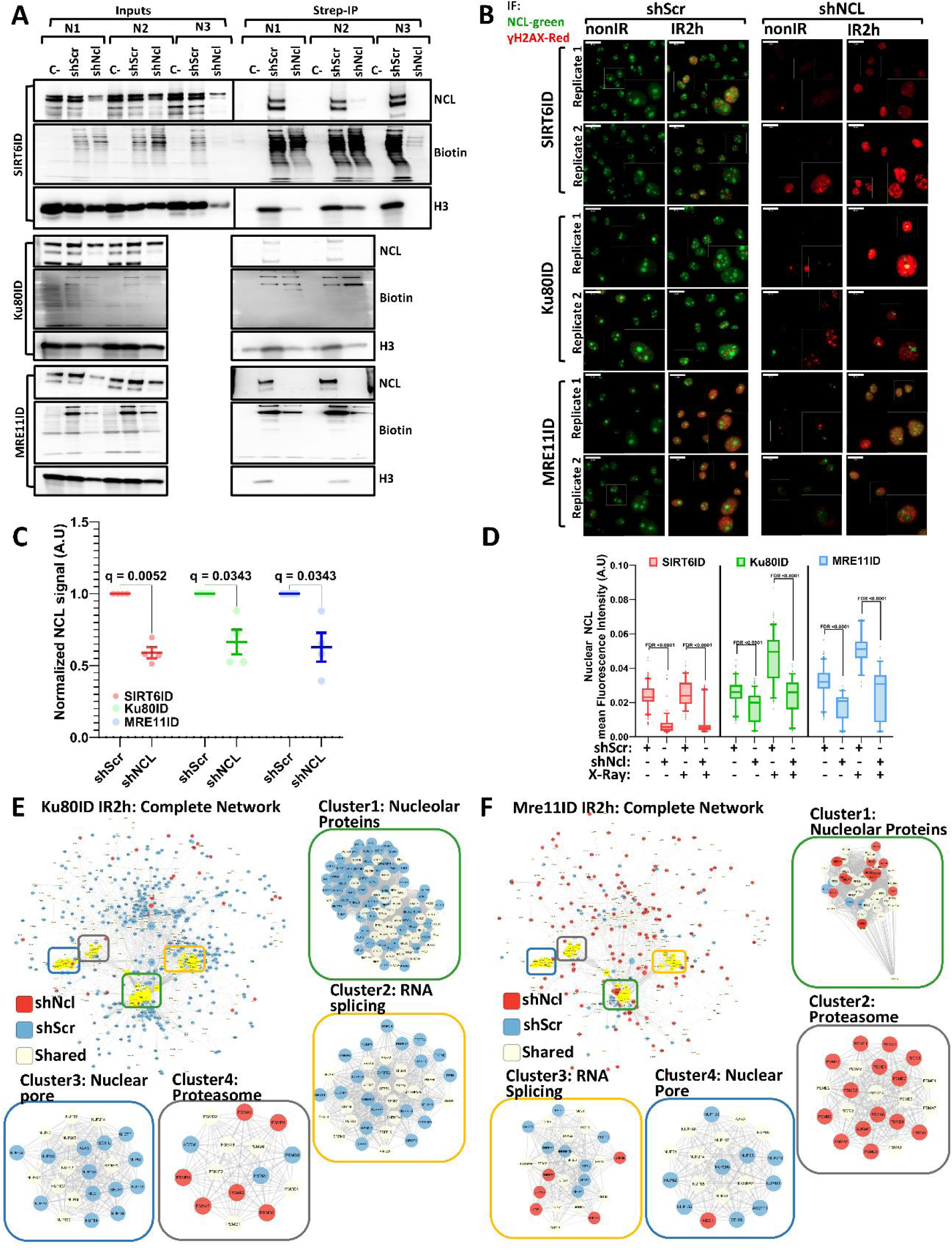
Validation of NCL depletion and extended shNCL network rewiring. **A)** Streptavidin pull-down and input immunoblots from SIRT6ID, Ku80ID, and MRE11ID SH-SY5Y cells transduced with shScr or shNCL and probed for NCL, biotinylated proteins, and histone H3. **B)** Representative immunofluorescence images showing NCL and γH2AX in shScr-and shNCL-transduced SPARK-ID cells under non-irradiated and IR2h conditions. **C-D)** Quantification of NCL depletion by immunoblot densitometry and nuclear NCL mean fluorescence intensity. **E-F)** Extended MCODE analysis of **E)** Ku80ID and **F)** MRE11ID networks after NCL depletion, highlighting nucleolar, RNA-splicing, nuclear-pore, and proteasome-associated subnetworks.

**Supplementary Figure 9.**
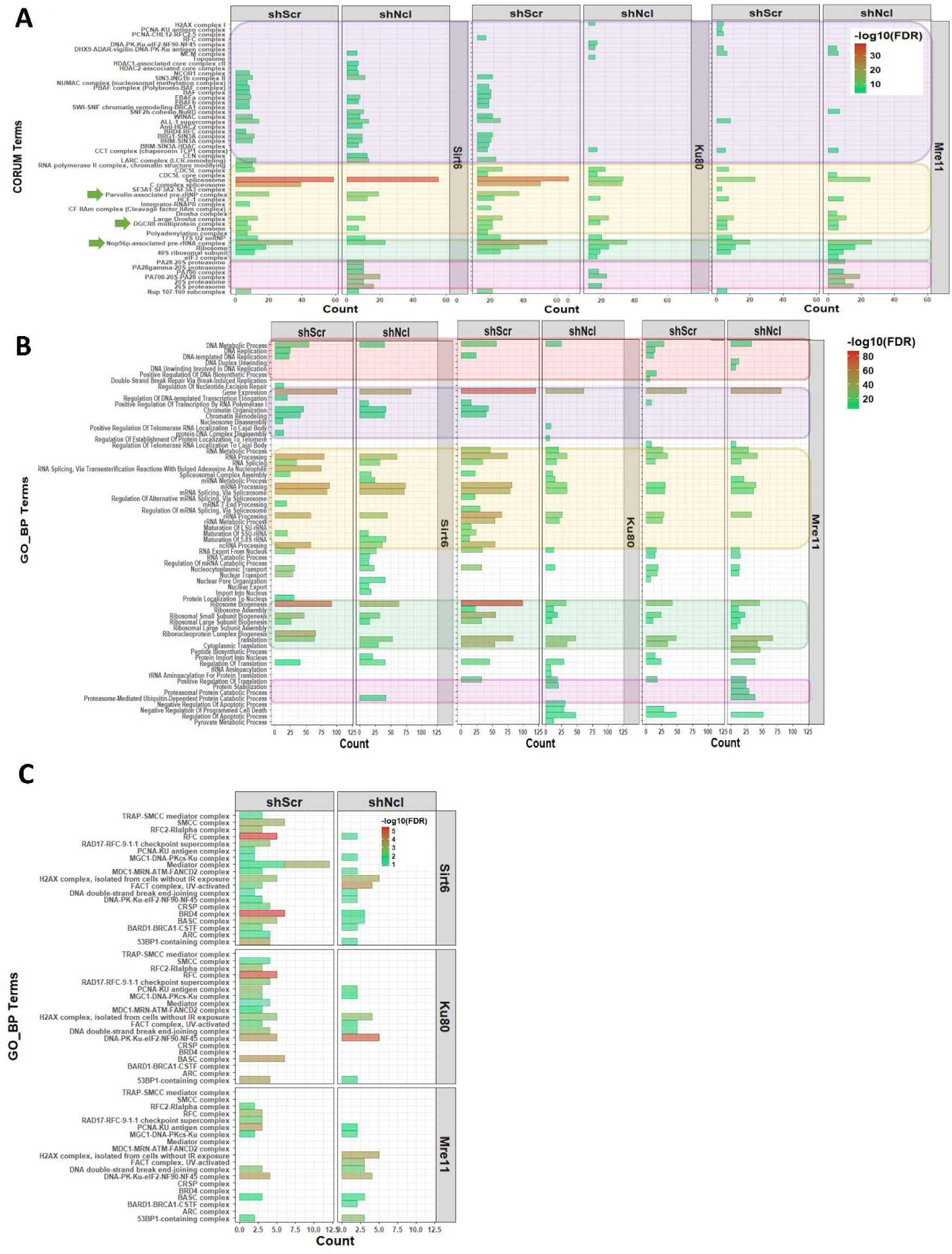
NCL depletion remodels RNA, nucleolar, proteostasis, chromatin, and DNA repair programs. **A)** CORUM enrichment of chromatin, RNA-metabolism, ribosome-biogenesis, and proteostasis complexes in shScr and shNCL SPARK-ID networks. **B)** GO biological-process enrichment of SIRT6ID, Ku80ID, and MRE11ID networks after shScr or shNCL treatment. **C)** CORUM enrichment of DNA repair-associated complexes across shScr and shNCL SPARK-ID networks.

**Supplementary Figure 10.**
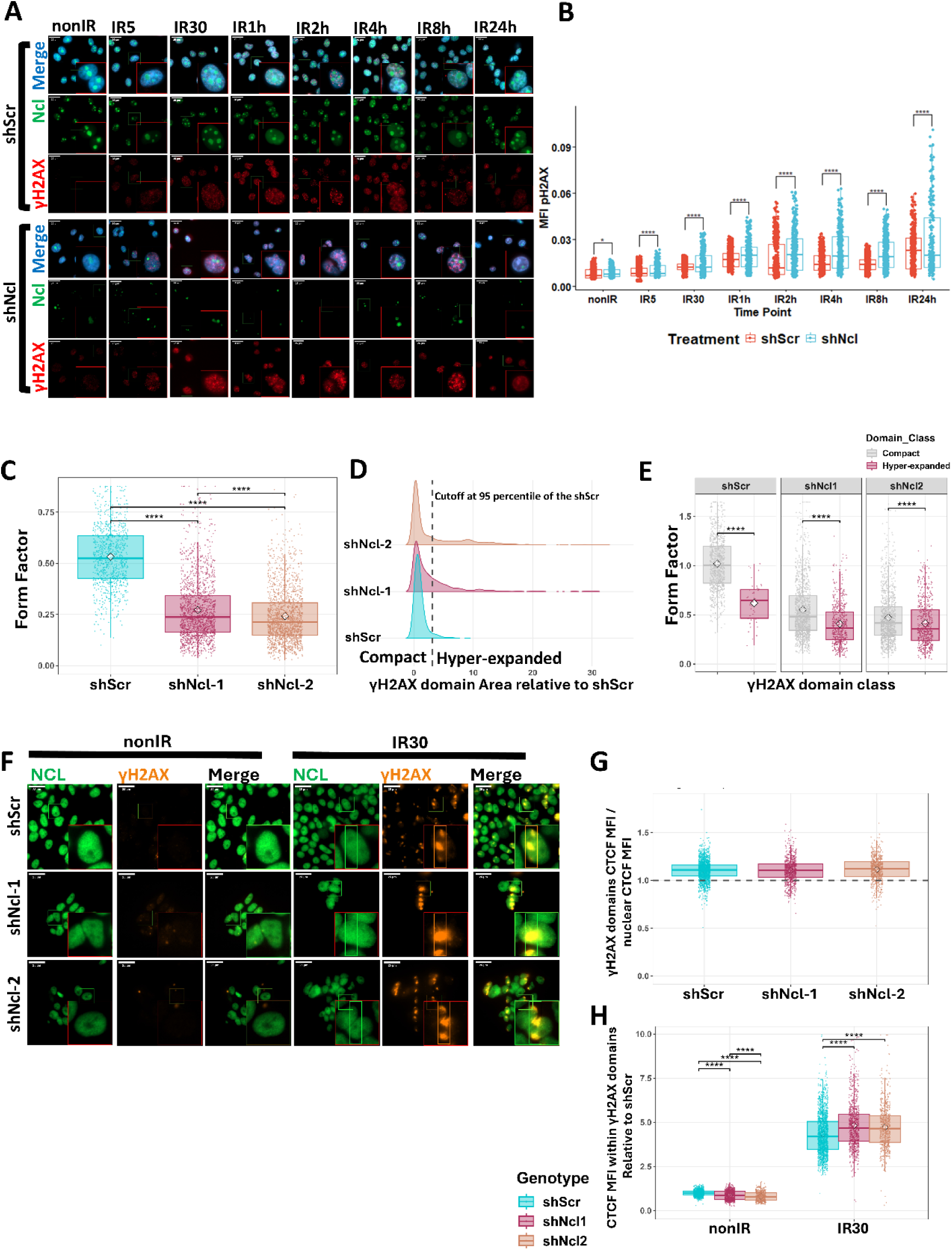
NCL depletion alters γH2AX-domain resolution and CTCF distribution after DNA damage. A-B) Extended γH2AX time-course analysis in SH-SY5Y cells transduced with shScr or shNCL, irradiated with 4 Gy, and fixed across DDR stages. **A)** Representative immunofluorescence images showing γH2AX, NCL, and DAPI. **B)** Quantification of nuclear γH2AX mean fluorescence intensity. **C-E)** Quantification of γH2AX-domain morphology after laser-induced DNA damage. **C)** Form factor was calculated as (4*π* × *area*)/*perimeter*^2^, with lower values indicating more irregular domains. **D)** The 95th percentile of the shScr γH2AX-domain area distribution was used to classify domains as compact or hyper-expanded. **E)** Hyper-expanded γH2AX domains displayed reduced form factor, indicating increased border irregularity. Statistical significance was assessed using Kruskal-Wallis test with Dunn’s post-hoc test; n = 4 biological replicates. **F-H)** CTCF distribution relative to γH2AX domains after laser-induced DNA damage. **F)** Representative images of non-irradiated and laser-damaged cells stained for CTCF and γH2AX. **G-H)** Quantification of CTCF intensity within γH2AX domains and relative enrichment over nuclear CTCF signal. Statistical significance was assessed using Kruskal-Wallis test with Dunn’s post-hoc test; n = 4 biological replicates.

**Supplementary Figure 11.**
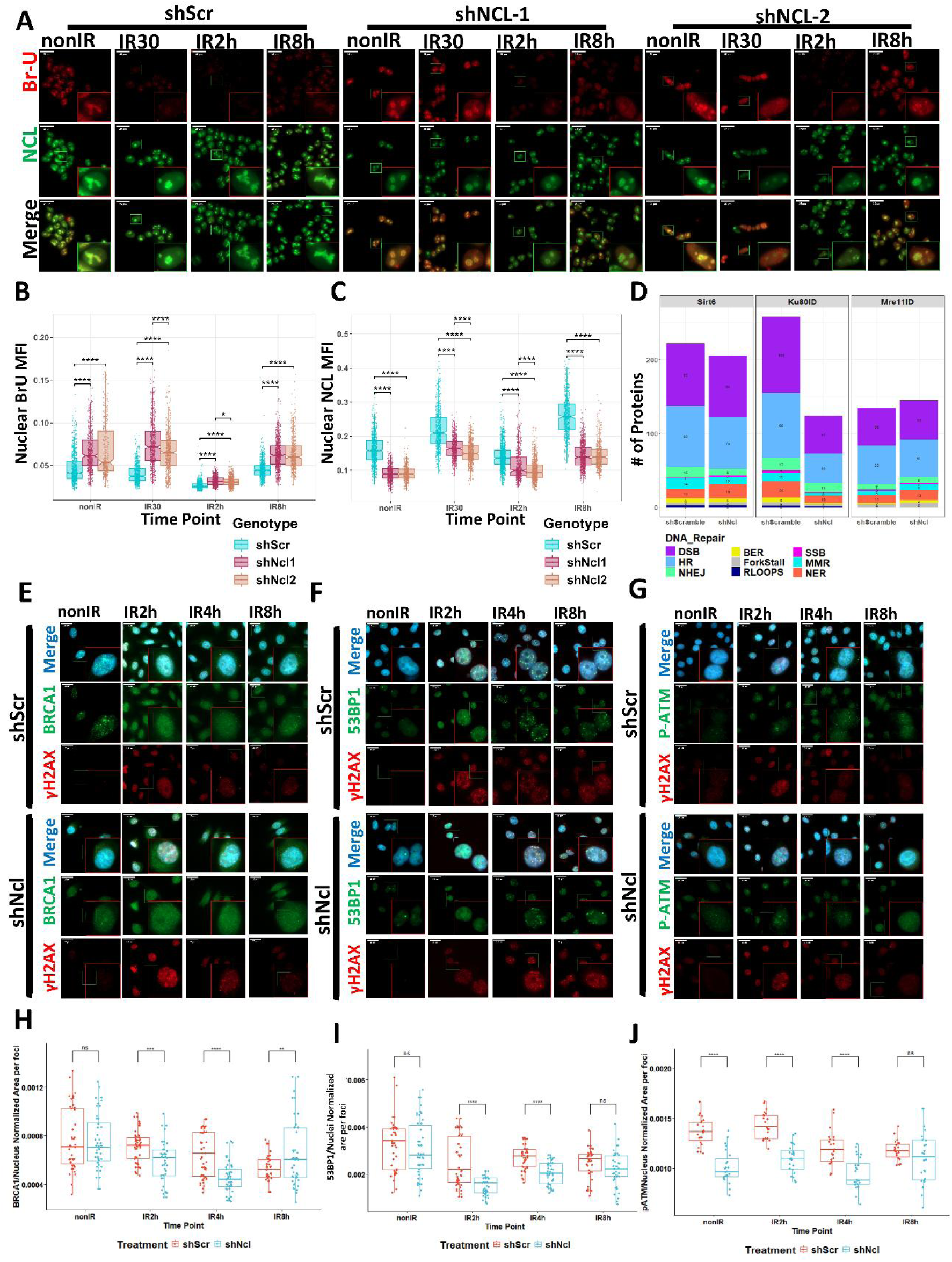
NCL depletion alters transcriptional recovery and repair-factor domain formation after DNA damage. A-C) Extended BrU metabolic-labeling analysis in HEK293 cells transduced with shScr, shNCL-1, or shNCL-2 and fixed after irradiation. **A)** Representative images showing BrU, NCL, and DAPI. **B-C)** Quantification of nuclear BrU mean fluorescence intensity and nuclear NCL mean fluorescence intensity. Statistical significance was assessed using Kruskal-Wallis test with Dunn’s post-hoc test and Bonferroni correction. **D)** Changes in the number of DNA repair-associated SPARK-ID interactors after NCL depletion, grouped by repair pathway annotation. **E-J)** Extended repair-factor foci analysis in SIRT6ID SH-SY5Y cells after NCL depletion. **E-G)** Representative immunofluorescence images for BRCA1, 53BP1, and phospho-ATM with γH2AX and DAPI across DDR time points. **H-J)** Quantification of BRCA1, 53BP1, and phospho-ATM foci area. Statistical significance was assessed using multiple t-tests with Bonferroni correction.

## References

1. Jeggo, P.A., and Löbrich, M. (2007). DNA double-strand breaks: Their cellular and clinical impact? Oncogene 2c, 7717–7719. 10.1038/SJ.ONC.1210868.

2. Sources of DNA Double-Strand Breaks and Models of Recombinational DNA Repair https://cshperspectives.cshlp.org/content/6/9/a016428.

3. Schumacher, B., Pothof, J., Vijg, J., and Hoeijmakers, J.H.J. (2021). The central role of DNA damage in the ageing process. Nature 2021 592:7856 592, 695–703. 10.1038/s41586-021-03307-7.

4. Tian, X., Firsanov, D., Zhang, Z., Cheng, Y., Luo, L., Tombline, G., Tan, R., Simon, M., Henderson, S., Steffan, J., et al. (2019). SIRT6 Is Responsible for More Efficient DNA Double-Strand Break Repair in Long-Lived Species. Cell 177, 622–638.e22. 10.1016/J.CELL.2019.03.043.

5. Maynard, S., Fang, E.F., Scheibye-Knudsen, M., Croteau, D.L., and Bohr, V.A. (2015). DNA Damage, DNA Repair, Aging, and Neurodegeneration. Cold Spring Harb. Perspect. Med. 5, a025130. 10.1101/CSHPERSPECT.A025130.

6. Barabási, A.L., Gulbahce, N., and Loscalzo, J. (2011). Network medicine: A network-based approach to human disease. Nat. Rev. Genet. 12, 56–68. 10.1038/NRG2918;SUBJMETA=144,154,208,2489,360,436,631,92;KWRD=DISEASE+GENETICS,NETWORKS+AND+SYSTEMS+BIOLOGY,PHARMACOLOGY.

7. Stoney, R., Robertson, D.L., Nenadic, G., and Schwartz, J.M. (2018). Mapping biological process relationships and disease perturbations within a pathway network. NPJ Syst. Biol. Appl. 4, 1–11. 10.1038/S41540-018-0055-2;SUBJMETA=114,2695,2704,553,631;KWRD=COMPUTATIONAL+BIOLOGY+AND+BIOINFORMATICS,COMPUTER+MODELLING,MODULARITY.

8. Kieffer, S.R., and Lowndes, N.F. (2022). Immediate-Early, Early, and Late Responses to DNA Double Stranded Breaks. Front. Genet. 13, 793884. 10.3389/FGENE.2022.793884/XML/NLM.

9. Kieffer, S.R., and Lowndes, N.F. (2022). Immediate-Early, Early, and Late Responses to DNA Double Stranded Breaks. Front. Genet. 13, 793884. 10.3389/FGENE.2022.793884/BIBTEX.

10. Petrini, J.H.J., and Stracker, T.H. (2003). The cellular response to DNA double-strand breaks: defining the sensors and mediators. Trends Cell Biol. 13, 458–462. 10.1016/S0962-8924(03)00170-3.

11. Onn, L., Portillo, M., Ilic, S., Cleitman, G., Stein, D., Kaluski, S., Shirat, I., Slobodnik, Z., Einav, M., Erdel, F., et al. (2020). SIRT6 is a DNA double-strand break sensor. Elife S. 10.7554/ELIFE.51636.

12. Aleksandrov, R., Hristova, R., Stoynov, S., and Gospodinov, A. (2020). The Chromatin Response to Double-Strand DNA Breaks and Their Repair. Cells 2020, Vol. 9, Page 1853 *9*, 1853. 10.3390/CELLS9081853.

13. Tan, J., Sun, X., Zhao, H., Guan, H., Gao, S., and Zhou, P.K. (2023). Double-strand DNA break repair: molecular mechanisms and therapeutic targets. MedComm (Beijing). 4, e388. 10.1002/MCO2.388.

14. Geng, A., Tang, H., Huang, J., Qian, Z., Qin, N., Yao, Y., Xu, Z., Chen, H., Lan, L., Xie, H., et al. (2020). The deacetylase SIRT6 promotes the repair of UV-induced DNA damage by targeting DDB2. Nucleic Acids Res. 48, 9181–9194. 10.1093/NAR/GKAA661.

15. Toiber, D., Erdel, F., Bouazoune, K., Silberman, D.M., Zhong, L., Mulligan, P., Sebastian, C., Cosentino, C., Martinez-Pastor, B., Giacosa, S., et al. (2013). SIRT6 Recruits SNF2H to DNA Break Sites, Preventing Genomic Instability through Chromatin Remodeling. Mol. Cell 51, 454–468. 10.1016/J.MOLCEL.2013.06.018.

16. Mostoslavsky, R., Chua, K.F., Lombard, D.B., Pang, W.W., Fischer, M.R., Gellon, L., Liu, P., Mostoslavsky, G., Franco, S., Murphy, M.M., et al. (2006). Genomic Instability and Aging-like Phenotype in the Absence of Mammalian SIRT6. Cell 124, 315–329. 10.1016/J.CELL.2005.11.044.

17. Wang, M., Wu, W., Wu, W., Rosidi, B., Zhang, L., Wang, H., and Iliakis, G. (2006). PARP-1 and Ku compete for repair of DNA double strand breaks by distinct NHEJ pathways. Nucleic Acids Res. 34, 6170–6182. 10.1093/NAR/GKL840.

18. Caron, M.C., Sharma, A.K., O’Sullivan, J., Myler, L.R., Ferreira, M.T., Rodrigue, A., Coulombe, Y., Ethier, C., Gagné, J.P., Langelier, M.F., et al. (2019). Poly(ADP-ribose) polymerase-1 antagonizes DNA resection at double-strand breaks. Nature Communications 2019 10:1 *10*, 1–16. 10.1038/s41467-019-10741-9.

19. Ray Chaudhuri, A., and Nussenzweig, A. (2017). The multifaceted roles of PARP1 in DNA repair and chromatin remodelling. Nature Reviews Molecular Cell Biology 2017 18:10 18, 610–621. 10.1038/nrm.2017.53.

20. Stinson, B.M., and Loparo, J.J. (2021). Repair of DNA Double-Strand Breaks by the Nonhomologous End Joining Pathway. Annu. Rev. Biochem. 90, 137–164. 10.1146/ANNUREV-BIOCHEM-080320-110356.

21. Abbasi, S., and Schild-Poulter, C. (2019). Mapping the Ku Interactome Using Proximity-Dependent Biotin Identification in Human Cells. J. Proteome Res. 18, 1064–1077. 10.1021/ACS.JPROTEOME.8B00771/SUPPL_FILE/PR8B00771_SI_005.XLSX.

22. Bayat, L., Abbasi, S., Balasuriya, N., and Schild-Poulter, C. (2024). Critical residues in the Ku70 von Willebrand A domain mediate Ku interaction with the LigIV-XRCC4 complex in non-homologous end-joining. Biochimica et Biophysica Acta (BBA) - Molecular Cell Research 1871, 119815. 10.1016/J.BBAMCR.2024.119815.

23. Trinkle-Mulcahy, L. (2019). Recent advances in proximity-based labeling methods for interactome mapping. F1000Res. 8. 10.12688/F1000RESEARCH.16903.1/DOI.

24. Mosler, T., Baymaz, H.I., Gräf, J.F., Mikicic, I., Blattner, G., Bartlett, E., Ostermaier, M., Piccinno, R., Yang, J., Voigt, A., et al. (2022). PARP1 proximity proteomics reveals interaction partners at stressed replication forks. Nucleic Acids Res. 50, 11600–11618. 10.1093/NAR/GKAC948.

25. Roux, K.J., Kim, D.I., Burke, B., and May, D.G. (2018). BioID: A Screen for Protein-Protein Interactions. Curr. Protoc. Protein Sci. 91, 19.23.1-19.23.15. 10.1002/CPPS.51.

26. Sears, R.M., May, D.G., and Roux, K.J. (2019). BioID as a Tool for Protein-Proximity Labeling in Living Cells. Methods in Molecular Biology 2012, 299–313. 10.1007/978-1-4939-9546-2_15.

27. Banaszynski, L.A., Sellmyer, M.A., Contag, C.H., Wandless, T.J., and Thorne, S.H. (2008). Chemical control of protein stability and function in living mice. Nature Medicine 2008 14:10 *14*, 1123–1127. 10.1038/nm.1754.

28. Roux, K.J., Kim, D.I., Raida, M., and Burke, B. (2012). A promiscuous biotin ligase fusion protein identifies proximal and interacting proteins in mammalian cells. Journal of Cell Biology 1Sc, 801–810. 10.1083/JCB.201112098.

29. Tisi, R., Vertemara, J., Zampella, G., and Longhese, M.P. (2020). Functional and structural insights into the MRX/MRN complex, a key player in recognition and repair of DNA double-strand breaks. Comput. Struct. Biotechnol. J. 18, 1137–1152. 10.1016/J.CSBJ.2020.05.013.

30. Olivieri, M., Cho, T., Álvarez-Quilón, A., Li, K., Schellenberg, M.J., Zimmermann, M., Hustedt, N., Rossi, S.E., Adam, S., Melo, H., et al. (2020). A Genetic Map of the Response to DNA Damage in Human Cells. Cell 182, 481–496.e21. 10.1016/J.CELL.2020.05.040.

31. Smogorzewska, A., Desetty, R., Saito, T.T., Schlabach, M., Lach, F.P., Sowa, M.E., Clark, A.B., Kunkel, T.A., Harper, J.W., Colaiácovo, M.P., et al. (2010). A genetic screen identifies FAN1, a Fanconi anemia-associated nuclease necessary for DNA interstrand crosslink repair. Mol. Cell 39, 36–47. 10.1016/J.MOLCEL.2010.06.023.

32. Adamson, B., Smogorzewska, A., Sigoillot, F.D., King, R.W., and Elledge, S.J. (2012). A genome-wide homologous recombination screen identifies the RNA-binding protein RBMX as a component of the DNA-damage response. Nature Cell Biology 2012 14:3 *14*, 318–328. 10.1038/ncb2426.

33. Brunette, G.J., Jamalruddin, M.A., Baldock, R.A., Clark, N.L., and Bernstein, K.A. (2019). Evolution-based screening enables genome-wide prioritization and discovery of DNA repair genes. Proc. Natl. Acad. Sci. U. S. A. 116, 19593–19599. 10.1073/PNAS.1906559116/SUPPL_FILE/PNAS.1906559116.SD02.XLSX.

34. López-Saavedra, A., Gómez-Cabello, D., Domínguez-Sánchez, M.S., Mejías-Navarro, F., Fernández-Ávila, M.J., Dinant, C., Martínez-Macías, M.I., Bartek, J., and Huertas, P. (2016). A genome-wide screening uncovers the role of CCAR2 as an antagonist of DNA end resection. Nature Communications 2016 7:1 *7*, 1–14. 10.1038/ncomms12364.

35. Milacic, M., Beavers, D., Conley, P., Gong, C., Gillespie, M., Griss, J., Haw, R., Jassal, B., Matthews, L., May, B., et al. (2024). The Reactome Pathway Knowledgebase 2024. Nucleic Acids Res. 52, D672–D678. 10.1093/NAR/GKAD1025.

36. Consortium, T.G.O., Aleksander, S.A., Balhoff, J., Carbon, S., Cherry, J.M., Drabkin, H.J., Ebert, D., Feuermann, M., Gaudet, P., Harris, N.L., et al. (2023). The Gene Ontology knowledgebase in 2023. Genetics 224. 10.1093/GENETICS/IYAD031.

37. Guimerà, R., and Amaral, L.A.N. (2005). Functional cartography of complex metabolic networks. Nature 433, 895–900. 10.1038/NATURE03288;KWRD=SCIENCE.

38. Guimerà, R., and Amaral, L.A.N. (2005). Functional cartography of complex metabolic networks. Nature 2005 433:7028 433, 895–900. 10.1038/nature03288.

39. Guimerà, R., and Amaral, L.A.N. (2005). Cartography of complex networks: modules and universal roles. Journal of Statistical Mechanics: Theory and Experiment 2005, P02001. 10.1088/1742-5468/2005/02/P02001.

40. Olesen, J.M., Bascompte, J., Dupont, Y.L., and Jordano, P. (2007). The modularity of pollination networks. Proc. Natl. Acad. Sci. U. S. A. 104, 19891–19896. 10.1073/PNAS.0706375104/SUPPL_FILE/INDEX.HTML.

41. Hayano, T., Yanagida, M., Yamauchi, Y., Shinkawa, T., Isobe, T., and Takahashi, N. (2003). Proteomic Analysis of Human Nop56p-associated Pre-ribosomal Ribonucleoprotein Complexes: POSSIBLE LINK BETWEEN Nop56p AND THE NUCLEOLAR PROTEIN TREACLE RESPONSIBLE FOR TREACHER COLLINS SYNDROME. Journal of Biological Chemistry 278, 34309–34319. 10.1074/JBC.M304304200.

42. Shiohama, A., Sasaki, T., Noda, S., Minoshima, S., and Shimizu, N. (2007). Nucleolar localization of DGCR8 and identification of eleven DGCR8-associated proteins. Exp. Cell Res. 313, 4196–4207. 10.1016/J.YEXCR.2007.07.020.

43. Ju, B.G., Solum, D., Song, E.J., Lee, K.J., Rose, D.W., Glass, C.K., and Rosenfeld, M.G. (2004). Activating the PARP-1 Sensor Component of the Groucho/ TLE1 Corepressor Complex Mediates a CaMKinase IIδ-Dependent Neurogenic Gene Activation Pathway. Cell 119, 815–829. 10.1016/J.CELL.2004.11.017.

44. Caron, P., Aymard, F., Iacovoni, J.S., Briois, S., Canitrot, Y., Bugler, B., Massip, L., Losada, A., and Legube, G. (2012). Cohesin Protects Genes against γH2AX Induced by DNA Double-Strand Breaks. PLoS Genet. 8, e1002460. 10.1371/JOURNAL.PGEN.1002460.

45. Arnould, C., Rocher, V., Finoux, A.L., Clouaire, T., Li, K., Zhou, F., Caron, P., Mangeot, P.E., Ricci, E.P., Mourad, R., et al. (2021). Loop extrusion as a mechanism for formation of DNA damage repair foci. Nature 590, 660–665. 10.1038/s41586-021-03193-z.

46. Aymard, F., Aguirrebengoa, M., Guillou, E., Javierre, B.M., Bugler, B., Arnould, C., Rocher, V., Iacovoni, J.S., Biernacka, A., Skrzypczak, M., et al. (2017). Genome-wide mapping of long-range contacts unveils clustering of DNA double-strand breaks at damaged active genes. Nat. Struct. Mol. Biol. 24, 353–361. 10.1038/nsmb.3387.

47. Tan, J., Sun, X., Zhao, H., Guan, H., Gao, S., and Zhou, P.K. (2023). Double-strand DNA break repair: molecular mechanisms and therapeutic targets. MedComm (Beijing). 4. 10.1002/MCO2.388.

48. Shrivastav, M., De Haro, L.P., and Nickoloff, J.A. (2008). Regulation of DNA double-strand break repair pathway choice. Cell Res. 18, 134–147. 10.1038/CR.2007.111.

49. McPherson, K.S., and Korzhnev, D.M. (2021). Targeting protein-protein interactions in the DNA damage response pathways for cancer chemotherapy. RSC Chem. Biol. 2, 1167–1195. 10.1039/D1CB00101A.

50. Abbasi, S., Bayat, L., and Schild-Poulter, C. (2023). Analysis of Ku70 S155 Phospho-Specific BioID2 Interactome Identifies Ku Association with TRIP12 in Response to DNA Damage. Int. J. Mol. Sci. 24, 7041. 10.3390/IJMS24087041/S1.

51. Lee, N., Kim, D.K., Kim, E.S., Park, S.J., Kwon, J.H., Shin, J., Park, S.M., Moon, Y.H., Wang, H.J., Gho, Y.S., et al. (2014). Comparative interactomes of SIRT6 and SIRT7: Implication of functional links to aging. Proteomics 14, 1610–1622. 10.1002/PMIC.201400001.

52. Miteva, Y. V., and Cristea, I.M. (2014). A Proteomic Perspective of Sirtuin 6 (SIRT6) Phosphorylation and Interactions and Their Dependence on Its Catalytic Activity. Molecular & Cellular Proteomics 13, 168–183. 10.1074/MCP.M113.032847.

53. Simeoni, F., Tasselli, L., Tanaka, S., Villanova, L., Hayashi, M., Kubota, K., Isono, F., Garcia, B.A., Michishita-Kioi, E., and Chua, K.F. (2013). Proteomic analysis of the SIRT6 interactome: novel links to genome maintenance and cellular stress signaling. Scientific Reports 2013 3:1 3, 1–6. 10.1038/srep03085.

54. Oughtred, R., Rust, J., Chang, C., Breitkreutz, B.J., Stark, C., Willems, A., Boucher, L., Leung, G., Kolas, N., Zhang, F., et al. (2021). The BioGRID database: A comprehensive biomedical resource of curated protein, genetic, and chemical interactions. Protein Science 30, 187–200. 10.1002/PRO.3978.

55. Lee, N., Kim, D.K., Kim, E.S., Park, S.J., Kwon, J.H., Shin, J., Park, S.M., Moon, Y.H., Wang, H.J., Gho, Y.S., et al. (2014). Comparative interactomes of SIRT6 and SIRT7: Implication of functional links to aging. Proteomics 14, 1610–1622. 10.1002/PMIC.201400001.

56. Aleksandrov, R., Dotchev, A., Poser, I., Krastev, D., Georgiev, G., Panova, G., Babukov, Y., Danovski, G., Dyankova, T., Hubatsch, L., et al. (2018). Protein Dynamics in Complex DNA Lesions. Mol. Cell cS, 1046-1061.e5. 10.1016/J.MOLCEL.2018.02.016/ATTACHMENT/95D5C65C-D518-492E-B3CA-C614C222C05D/MMC11.PDF.

57. 57. Kochan, J.A., Desclos, E.C.B., Bosch, R., Meister, L., Vriend, L.E.M., Van Attikum, H., and Krawczyk, P.M. (2017). Meta-analysis of DNA double-strand break response kinetics. Nucleic Acids Res. 45, 12625–12637. 10.1093/NAR/GKX1128.

58. Szklarczyk, D., Kirsch, R., Koutrouli, M., Nastou, K., Mehryary, F., Hachilif, R., Gable, A.L., Fang, T., Doncheva, N.T., Pyysalo, S., et al. (2023). The STRING database in 2023: protein–protein association networks and functional enrichment analyses for any sequenced genome of interest. Nucleic Acids Res. 51, D638–D646. 10.1093/NAR/GKAC1000.

59. Mao, Z., Bozzella, M., Seluanov, A., and Gorbunova, V. (2008). Comparison of nonhomologous end joining and homologous recombination in human cells. DNA Repair (Amst). 7, 1765–1771. 10.1016/J.DNAREP.2008.06.018.

60. Myler, L.R., Gallardo, I.F., Soniat, M.M., Deshpande, R.A., Gonzalez, X.B., Kim, Y., Paull, T.T., and Finkelstein, I.J. (2017). Single-Molecule Imaging Reveals How Mre11-Rad50-Nbs1 Initiates DNA Break Repair. Mol. Cell 67, 891–898.e4. 10.1016/J.MOLCEL.2017.08.002.

61. Mao, Z., Bozzella, M., Seluanov, A., and Gorbunova, V. (2008). DNA repair by nonhomologous end joining and homologous recombination during cell cycle in human cells. Cell Cycle 7, 2902–2906. 10.4161/CC.7.18.6679.

62. Shibata, A., Conrad, S., Birraux, J., Geuting, V., Barton, O., Ismail, A., Kakarougkas, A., Meek, K., Taucher-Scholz, G., Löbrich, M., et al. (2011). Factors determining DNA double-strand break repair pathway choice in G2 phase. EMBO Journal 30, 1079–1092. 10.1038/EMBOJ.2011.27/SUPPL_FILE/EMBJ201127.REVIEWER_COMMENTS.PD F.

63. Katsuki, Y., Jeggo, P.A., Uchihara, Y., Minoru Takata, ·, and Shibata, · Atsushi (2020). DNA double-strand break end resection: a critical relay point for determining the pathway of repair and signaling. Genome Instability & Disease 2020 1:4 *1*, 155–171. 10.1007/S42764-020-00017-8.

64. Rai, R., Gu, P., Broton, C., Kumar-Sinha, C., Chen, Y., and Chang, S. (2019). The Replisome Mediates A-NHEJ Repair of Telomeres Lacking POT1-TPP1 Independently of MRN Function. Cell Rep. 29, 3708–3725.e5. 10.1016/J.CELREP.2019.11.012.

65. Bai, Y., Wang, W., Li, S., Zhan, J., Li, H., Zhao, M., Zhou, X.A., Li, S., Li, X., Huo, Y., et al. (2019). C1QBP Promotes Homologous Recombination by Stabilizing MRE11 and Controlling the Assembly and Activation of MRE11/RAD50/NBS1 Complex. Mol. Cell 75, 1299–1314.e6. 10.1016/J.MOLCEL.2019.06.023.

66. Lu, H., Yang, M., and Zhou, Q. (2023). Reprogramming transcription after DNA damage: recognition, response, repair, and restart. Trends Cell Biol. 33, 682–694. 10.1016/J.TCB.2022.11.010.

67. Chang, A.R., Ferrer, C.M., and Mostoslavsky, R. (2020). SIRT6, a mammalian deacylase with multitasking abilities. Physiol. Rev. 100, 145–169. 10.1152/PHYSREV.00030.2018/ASSET/IMAGES/LARGE/Z9J0012029210006.JPE G.

68. Machour, F.E., and Ayoub, N. (2020). Transcriptional Regulation at DSBs: Mechanisms and Consequences. Trends in Genetics 36, 981–997. 10.1016/J.TIG.2020.01.001/ASSET/7C66F2DA-EA41-4971-97B7-A7790A16AD3E/MAIN.ASSETS/GR3.JPG.

69. Ui, A., Chiba, N., and Yasui, A. (2020). Relationship among DNA double-strand break (DSB), DSB repair, and transcription prevents genome instability and cancer. Cancer Sci. 111, 1443– 1451. 10.1111/CAS.14404.

70. Bader, A.S., Hawley, B.R., Wilczynska, A., and Bushell, M. (2020). The roles of RNA in DNA double-strand break repair. British Journal of Cancer 2020 122:5 122, 613–623. 10.1038/s41416-019-0624-1.

71. Chakraborty, A., Tapryal, N., Venkova, T., Horikoshi, N., Pandita, R.K., Sarker, A.H., Sarkar, P.S., Pandita, T.K., and Hazra, T.K. (2016). Classical non-homologous end-joining pathway utilizes nascent RNA for error-free double-strand break repair of transcribed genes. Nature Communications 2016 7:1 *7*, 1–12. 10.1038/ncomms13049.

72. A. Welty, S., Teng, Y., Liang, Z., Zhao, W., Sanders, L.H., Greenamyre, J.T., Rubio, M.E., Thathiah, , Kodali, R., Wetzel, R., et al. (2018). RAD52 is required for RNA-templated recombination repair in post-mitotic neurons. Journal of Biological Chemistry 293, 1353–1362. 10.1074/jbc.M117.808402.

73. Jeon, Y., Lu, Y., Ferrari, M.M., Channagiri, T., Xu, P., Meers, C., Zhang, Y., Balachander, S., Park, V.S., Marsili, S., et al. (2024). RNA-mediated double-strand break repair by end-joining mechanisms. Nature Communications 2024 15:1 *15*, 1–24. 10.1038/s41467-024-51457-9.

74. Chen, F., Xu, W., Tang, M., Tian, Y., Shu, Y., He, X., Zhou, L., Liu, Q., Zhu, Q., Lu, X., et al. (2024). hnRNPA2B1 deacetylation by SIRT6 restrains local transcription and safeguards genome stability. Cell Death & Differentiation 2024, 1–15. 10.1038/s41418-024-01412-4.

75. Simon, M.N., Dubrana, K., and Palancade, B. (2024). On the edge: how nuclear pore complexes rule genome stability. Curr. Opin. Genet. Dev. 84, 102150. 10.1016/J.GDE.2023.102150.

76. Chung, D.K.C., Chan, J.N.Y., Strecker, J., Zhang, W., Ebrahimi-Ardebili, S., Lu, T., Abraham, K.J., Durocher, D., and Mekhail, K. (2015). Perinuclear tethers license telomeric DSBs for a broad kinesin- and NPC-dependent DNA repair process. Nature Communications 2015 6:1 *6*, 1–13. 10.1038/ncomms8742.

77. Horigome, C., Bustard, D.E., Marcomini, I., Delgoshaie, N., Tsai-Pflugfelder, M., Cobb, J.A., and Gasser, S.M. (2016). PolySUMOylation by Siz2 and Mms21 triggers relocation of DNA breaks to nuclear pores through the Slx5/Slx8 STUbL. Genes Dev. 30, 931–945. 10.1101/GAD.277665.116.

78. Chen, B., Ge, T., Jian, M., Chen, L., Fang, Z., He, Z., Huang, C., An, Y., Yin, S., Xiong, Y., et al. (2023). Transmembrane nuclease NUMEN/ENDOD1 regulates DNA repair pathway choice at the nuclear periphery. Nature Cell Biology 2023 25:7 *25*, 1004–1016. 10.1038/s41556-023-01165-1.

79. 78. von Mikecz, A. (2006). The nuclear ubiquitin-proteasome system. J. Cell Sci. 119, 1977–1984. 10.1242/JCS.03008.

80. Krogan, N.J., Lam, M.H.Y., Fillingham, J., Keogh, M.C., Gebbia, M., Li, J., Datta, N., Cagney, G., Buratowski, S., Emili, A., et al. (2004). Proteasome involvement in the repair of DNA double-strand breaks. Mol. Cell 16, 1027–1034. 10.1016/j.molcel.2004.11.033.

81. Cron, K.R., Zhu, K., Kushwaha, D.S., Hsieh, G., Merzon, D., Rameseder, J., Chen, C.C., D’Andrea, A.D., and Kozono, D. (2013). Proteasome Inhibitors Block DNA Repair and Radiosensitize Non-Small Cell Lung Cancer. PLoS One 8, e73710. 10.1371/JOURNAL.PONE.0073710.

82. Murakawa, Y., Sonoda, E., Barber, L.J., Zeng, W., Yokomori, K., Kimura, H., Niimi, A., Lehmann, A., Guang, Y.Z., Hochegger, H., et al. (2007). Inhibitors of the Proteasome Suppress Homologous DNA Recombination in Mammalian Cells. Cancer Res. 67, 8536–8543. 10.1158/0008-5472.CAN-07-1166.

83. Gudmundsdottir, K., Lord, C.J., and Ashworth, A. (2007). The proteasome is involved in determining differential utilization of double-strand break repair pathways. Oncogene 2007 26:54 *26*, 7601–7606. 10.1038/sj.onc.1210579.

84. 83. Levy-Barda, A., Lerenthal, Y., Davis, A.J., Chung, Y.M., Essers, J., Shao, Z., Van Vliet, N., Chen, D.J., Hu, M.C.T., Kanaar, R., et al. (2011). Involvement of the nuclear proteasome activator PA28γ in the cellular response to DNA double-strand breaks. Cell Cycle 10, 4300–4310. 10.4161/CC.10.24.18642.

85. Korsholm, L.M., Gál, Z., Nieto, B., Quevedo, O., Boukoura, S., Lund, C.C., and Larsen, D.H. (2020). Recent advances in the nucleolar responses to DNA double-strand breaks. Nucleic Acids Res. 48, 9449–9461. 10.1093/NAR/GKAA713.

86. Zhang, X., Khan, S., Jiang, H., Antonyak, M.A., Chen, X., Spiegelman, N.A., Shrimp, J.H., Cerione, R.A., and Lin, H. (2016). Identifying the functional contribution of the defatty-acylase activity of SIRT6. Nature Chemical Biology 2016 12:8 *12*, 614–620. 10.1038/nchembio.2106.

87. Toiber, D., Stein, D., Portillo, M., Kopatch, S.K.-, Stein, D., Lachberg, Y., Eremenko, E., Smirnov, D., Einav, M., Khrameeva, E., et al. (2024). SIRT6 regulates protein synthesis and folding through nucleolar remodeling. 10.21203/RS.3.RS-4215918/V1.

88. Ogawa, L.M., and Baserga, S.J. (2017). Crosstalk between the nucleolus and the DNA damage response. Mol. Biosyst. 13, 443–455. 10.1039/C6MB00740F.

89. González-Arzola, K. (2024). The nucleolus: Coordinating stress response and genomic stability. Biochimica et Biophysica Acta (BBA) - Gene Regulatory Mechanisms 1867, 195029. 10.1016/J.BBAGRM.2024.195029.

90. Scott, D.D., and Oeffinger, M. (2016). Nucleolin and nucleophosmin: nucleolar proteins with multiple functions in DNA repair1. 10.1139/bcb-2016-0068 S4, 419–432. 10.1139/BCB-2016-0068.

91. Braunstein, S., Badura, M.L., Xi, Q., Formenti, S.C., and Schneider, R.J. (2009). Regulation of Protein Synthesis by Ionizing Radiation. Mol. Cell. Biol. 29, 5645–5656. 10.1128/MCB.00711-09.

92. Angelov, D., Bondarenko, V.A., Almagro, S., Menoni, H., Mongélard, F., Hans, F., Mietton, F., Studitsky, V.M., Hamiche, A., Dimitrov, S., et al. (2006). Nucleolin is a histone chaperone with FACT-like activity and assists remodeling of nucleosomes. EMBO Journal 25, 1669–1679. 10.1038/SJ.EMBOJ.7601046/ASSET/A95CCE31-4962-4947-A332-D96DBF573E5B/ASSETS/GRAPHIC/EMBJ7601046-FIG-0007-M.JPG.

93. Kobayashi, J., Fujimoto, H., Sato, J., Hayashi, I., Burma, S., Matsuura, S., Chen, D.J., and Komatsu, K. (2012). Nucleolin Participates in DNA Double-Strand Break-Induced Damage Response through MDC1-Dependent Pathway. PLoS One 7, e49245. 10.1371/JOURNAL.PONE.0049245.

94. Lim, K.H., Park, J.J., Gu, B.H., Kim, J.O., Park, S.G., and Baek, K.H. (2015). HAUSP-nucleolin interaction is regulated by p53-Mdm2 complex in response to DNA damage response. Scientific Reports 2015 5:1 *5*, 1–12. 10.1038/srep12793.

95. A. Goldstein, M., Derheimer, F.A., Tait-Mulder, J., and Kastan, M.B. (2013). Nucleolin mediates nucleosome disruption critical for DNA double-strand break repair. Proc. Natl. Acad. Sci. U. S. 110, 16874–16879. 10.1073/PNAS.1306160110/SUPPL_FILE/PNAS.201306160SI.PDF.

96. Ginisty, H., Sicard, H., Roger, B., and Bouvet, P. (1999). Structure and functions of nucleolin. J. Cell Sci. 112, 761–772. 10.1242/JCS.112.6.761.

97. Ma, N., Matsunaga, S., Takata, H., Ono-Maniwa, R., Uchiyama, S., and Fukui, K. (2007). Nucleolin functions in nucleolus formation and chromosome congression. J. Cell Sci. 120, 2091–2105. 10.1242/JCS.008771.

98. Toiber, D., Stein, D., Portillo, M., Kopatch, S.K.-, Stein, D., Lachberg, Y., Eremenko, E., Smirnov, D., Einav, M., Khrameeva, E., et al. (2024). SIRT6 regulates protein synthesis and folding through nucleolar remodeling. 10.21203/RS.3.RS-4215918/V1.

99. Portillo, M., Eremenko, E., Kaluski, S., Garcia-Venzor, A., Onn, L., Stein, D., Slobodnik, Z., Zaretsky, A., Ueberham, U., Einav, M., et al. (2021). SIRT6-CBP-dependent nuclear Tau accumulation and its role in protein synthesis. Cell Rep. 35, 109035. 10.1016/J.CELREP.2021.109035/ATTACHMENT/657A1236-DC98-4107-A0BA-03735C6089AD/MMC9.PDF.

100. Tanwar, V.S., Jose, C.C., and Cuddapah, S. (2019). Role of CTCF in DNA damage response. Mutation Research/Reviews in Mutation Research 780, 61–68. 10.1016/J.MRREV.2018.02.002.

101. Larsen, D.H., and Stucki, M. (2016). Nucleolar responses to DNA double-strand breaks. Nucleic Acids Res. 44, 538–544. 10.1093/NAR/GKV1312.

102. Ui, A., Chiba, N., and Yasui, A. (2020). Relationship among DNA double-strand break (DSB), DSB repair, and transcription prevents genome instability and cancer. Cancer Sci. 111, 1443– 1451. 10.1111/cas.14404.

103. Dadi, S., Payet-Bornet, D., and Ferrier, P. (2013). ImmunoPrecipitation of Nuclear Protein with Antibody Affinity Columns. Bio. Protoc. 3. 10.21769/BIOPROTOC.319.

104. Schindelin, J., Arganda-Carreras, I., Frise, E., Kaynig, V., Longair, M., Pietzsch, T., Preibisch, S., Rueden, C., Saalfeld, S., Schmid, B., et al. (2012). Fiji: An open-source platform for biological-image analysis. Nat. Methods 9, 676–682. 10.1038/NMETH.2019;SUBJMETA=1647,245,631,794;KWRD=IMAGING,SOFTWARE.

105. Stirling, D.R., Swain-Bowden, M.J., Lucas, A.M., Carpenter, A.E., Cimini, B.A., and Goodman, A. (2021). CellProfiler 4: improvements in speed, utility and usability. BMC Bioinformatics 22, 1–11. 10.1186/S12859-021-04344-9/FIGURES/6.

106. North, B.J., Marshall, B.L., Borra, M.T., Denu, J.M., and Verdin, E. (2003). The Human Sir2 Ortholog, SIRT2, Is an NAD+-Dependent Tubulin Deacetylase. Mol. Cell 11, 437–444. 10.1016/S1097-2765(03)00038-8.

107. Britton, S., Coates, J., and Jackson, S.P. (2013). A new method for high-resolution imaging of Ku foci to decipher mechanisms of DNA double-strand break repair. J. Cell Biol. 202, 579–595. 10.1083/JCB.201303073.

108. Xie, A., Kwok, A., and Scully, R. (2009). Role of mammalian Mre11 in classical and alternative nonhomologous end joining. Nature Structural & Molecular Biology 2009 16:8 *16*, 814–818. 10.1038/nsmb.1640.

109. Wu, L., Luo, K., Lou, Z., and Chen, J. (2008). MDC1 regulates intra-S-phase checkpoint by targeting NBS1 to DNA double-strand breaks. Proc. Natl. Acad. Sci. U. S. A. 105, 11200–11205. 10.1073/PNAS.0802885105/ASSET/5832C28A-425C-48E1-8F47-4B6BF09712C9/ASSETS/GRAPHIC/ZPQ9990840920005.JPEG.

110. Gibson, D.G. (2011). Enzymatic Assembly of Overlapping DNA Fragments. Methods Enzymol. 498, 349–361. 10.1016/B978-0-12-385120-8.00015-2.

111. Tiscornia, G., Singer, O., and Verma, I.M. (2006). Production and purification of lentiviral vectors. Nature Protocols 2006 1:1 *1*, 241–245. 10.1038/nprot.2006.37.

112. Kaluski, S., Portillo, M., Besnard, A., Stein, D., Einav, M., Zhong, L., Ueberham, U., Arendt, T., Mostoslavsky, R., Sahay, A., et al. (2017). Neuroprotective Functions for the Histone Deacetylase SIRT6. Cell Rep. 18, 3052–3062. 10.1016/J.CELREP.2017.03.008.

113. Cox, J., Hein, M.Y., Luber, C.A., Paron, I., Nagaraj, N., and Mann, M. (2014). Accurate Proteome-wide Label-free Quantification by Delayed Normalization and Maximal Peptide Ratio Extraction, Termed MaxLFQ. Mol. Cell. Proteomics 13, 2513. 10.1074/MCP.M113.031591.

114. Tyanova, S., Temu, T., Sinitcyn, P., Carlson, A., Hein, M.Y., Geiger, T., Mann, M., and Cox, J. (2016). The Perseus computational platform for comprehensive analysis of (prote)omics data. Nature Methods 2016 13:9 *13*, 731–740. 10.1038/nmeth.3901.

115. Kammers, K., Cole, R.N., Tiengwe, C., and Ruczinski, I. (2015). Detecting significant changes in protein abundance. EuPA Open Proteom. 7, 11–19. 10.1016/J.EUPROT.2015.02.002.

116. Kammers, K., Cole, R.N., Tiengwe, C., and Ruczinski, I. (2015). Detecting Significant Changes in Protein Abundance. EuPA Open Proteom. 7, 11. 10.1016/J.EUPROT.2015.02.002.

117. Shannon, P., Markiel, A., Ozier, O., Baliga, N.S., Wang, J.T., Ramage, D., Amin, N., Schwikowski, B., and Ideker, T. (2003). Cytoscape: a software environment for integrated models of biomolecular interaction networks. Genome Res. 13, 2498–2504. 10.1101/GR.1239303.

118. Csárdi, G., Nepusz, T., Müller, K., Horvát, S., Traag, V., Zanini, F., and Noom, D. igraph for R: R interface of the igraph library for graph theory and network analysis. 10.5281/ZENODO.14347716.

119. De Domenico, M., Porter, M.A., and Arenas, A. (2015). MuxViz: A tool for multilayer analysis and visualization of networks. J. Complex Netw. 3, 159–176. 10.1093/comnet/cnu038.

120. Guimerà, R., and Amaral, L.A.N. (2005). Functional cartography of complex metabolic networks. Nature 2005 433:7028 *433*, 895–900. 10.1038/nature03288.

121. Futschik, M.E., and Carlisle, B. (2011). NOISE-ROBUST SOFT CLUSTERING OF GENE EXPRESSION TIME-COURSE DATA. .10.1142/S0219720005001375 *3*, 965–988. 10.1142/S0219720005001375.

122. Tsitsiridis, G., Steinkamp, R., Giurgiu, M., Brauner, B., Fobo, G., Frishman, G., Montrone, C., and Ruepp, A. (2023). CORUM: the comprehensive resource of mammalian protein complexes– 2022. Nucleic Acids Res. 51, D539–D545. 10.1093/NAR/GKAC1015.

123. Bader, G.D., and Hogue, C.W.V. (2003). An automated method for finding molecular complexes in large protein interaction networks. BMC Bioinformatics 4, 1–27. 10.1186/1471-2105-4-2/FIGURES/12.

124. Gu, Z., Eils, R., and Schlesner, M. (2016). Complex heatmaps reveal patterns and correlations in multidimensional genomic data. Bioinformatics 32, 2847–2849. 10.1093/BIOINFORMATICS/BTW313.

125. Pérez-Silva, J.G., Araujo-Voces, M., and Quesada, V. (2018). nVenn: generalized, quasi-proportional Venn and Euler diagrams. Bioinformatics 34, 2322–2324. 10.1093/BIOINFORMATICS/BTY109.

